# Dietary emulsifiers alter composition and activity of the human gut microbiota *in vitro*, irrespective of chemical or natural emulsifier origin

**DOI:** 10.1101/2020.06.28.174946

**Authors:** Lisa Miclotte, Chris Callewaert, Kim de Paepe, Leen Rymenans, Jeroen Raes, Andreja Rajkovic, John Van Camp, Tom Van de Wiele

## Abstract

The use of additives in food products has become an important public health concern. In recent reports, dietary emulsifiers have been shown to affect the gut microbiota, contributing to a pro-inflammatory phenotype and metabolic syndrome. So far, it is not yet known whether similar microbiome shifts are observable for a more diverse set of emulsifier types and to what extent these effects vary with the unique features of an individual’s microbiome.

To bridge this gap, we investigated the effect of five dietary emulsifiers on the fecal microbiota from 10 human individuals upon a 48 hour exposure. Community structure was assessed with quantative microbial profiling, functionality was evaluated by measuring fermentation metabolites and pro-inflammatory properties were assessed with the phylogenetic prediction algorythm PICRUSt, together with a TLR5 reporter cell assay for flagellin. A comparison was made between two mainstream chemical emulsifiers (carboxymethylcellulose and P80), a natural extract (soy lecithin) and biotechnological emulsifiers (sophorolipids and rhamnolipids).

While fecal microbiota responded in a donor-dependent manner to the different emulsifiers, profound differences between emulsifier were observed. Rhamnolipids, sophorolipids and soy lecithin eliminated 91% ± 0%, 89% ± 1% and 87% ± 1% of the viable bacterial population after 48 hours, yet they all selectively increased the proportional abundance of putative pathogens. Moreover, profound shifts in butyrate (−96% ± 6 %, −73% ± 24% and −34 ± 25% respectively) and propionate (+13% ± 24 %, +88% ± 50% and +29% ± 16% respectively) production were observed for these emulsifiers. Phylogenetic prediction indicated higher motility, which was, however, not confirmed by increased flagellin levels using the TLR5 reporter cell assay.

We conclude that dietary emulsifiers can severely impact the gut microbiota and this seems to be proportional to their emulsifying strength, rather than emulsifier type or origin. As biotechnological emulsifiers were especially more impactful than chemical emulsifiers, caution is warranted when considering them as more natural alternatives for clean label strategies.

## 1 Introduction

The current obesity crisis and related health conditions are increasingly associated with the overconsumption of so-called ultra-processed food products (Waterlander et al. 2018; Broussard and Devkota 2016; Branca et al. 2019; Rauber et al. 2018; Carlos Augusto Monteiro et al. 2017). Food additives are characteristic elements of said products (Carocho et al. 2014; C. A. Monteiro et al. 2013; Carlos Augusto Monteiro et al. 2017) and are added to enhance, amongst others, shelf life, palatablility, texture, color and nutritional value. However, the health impact of certain food additives has always been questioned (Miclotte and Van de Wiele 2019; Carocho et al. 2014; Payne, Chassard, and Lacroix 2012), and at this moment, the use of food additives in food products is one of the main public concerns about food in Europe (EFSA 2019).

Diet is known to have a strong and fast impact on the gut microbiota (Martínez Steele et al. 2017; Musso, Gambino, and Cassader 2010; Ding et al. 2019), which is generally considered an important parameter of gut and overall health (Bischoff 2011; Ding et al. 2019; Musso, Gambino, and Cassader 2010). An unbalanced gut microbiota is being related to several physical and mental illnesses and conditions (Ding et al. 2019). With respect to obesity and NCDs, a dysbiosed gut microbiota is characterized by a lower alpha diversity and is related to increased harvest from food and decreased fatty acid oxidation, glucose tolerance, production of satiety hormones and intestinal barrier integrity (Musso, Gambino, and Cassader 2010).

Recently, research has emerged that ties the consumption of additives to health markers through the gut microbiota. Dietary emulsifiers in particular have been proposed to display a destabilizing impact on gut health. Chassaing et al (2015, 2017) found *in vivo* that sodium carboxymethylcellulose (CMC) and polysorbate 80 (P80) increase gut microbial motility and lower mucus layer thickness, yielding an increased production of pro-inflammatory compounds, low-grade gut inflammation and weight gain. Another study has linked glycerol monolaureate (GML) with signatures of metabolic syndrome together with alterations of gut microbiota composition, among which decreased abundance of *Akkermansia muciniphila* and increased abundance of *Escherichia coli* (Jiang et al. 2018). The latter study is particularly relevant since GML is one of the World’s most widely used dietary emulsifiers (E471).

Knowledge that is currently still lacking from literature regarding the impact of dietary emulsifiers is what characteristics of an emulsifier determine its destabilizing effects, whether alternative, more natural emulsifiers could be safer and to what extent the unique features of an individual’s microbiome play a role in the purported effects on the microbiome.

The present study describes the effects of five dietary emulsifiers: CMC, P80, soy lecithin, sophorolipids and rhamnolipids. The first two, CMC and P80 are synthetic emulsifiers that have been used for years and both are considered safe for human oral consumption. CMC is a water soluble anionic polymer with water-binding properties, due to which it is used as a thickener, emulsifier or water-retainer in applications like pharmaceuticals, food products, paper, cosmetics, detergents, etc. (Biswal and Singh 2004; Hercules Inc. and Aqualon 1999). In Europe, CMC can be used in many food products at *quantum satis* levels ^1^(European Commision 2014) and in the USA, CMC caries the GRAS-status (generally recognized as safe) for applications in food ^2,3^(FDA 2020, 2019b). P80 is a member of the polysorbates, a group of non-ionic surfactants with applications mainly in the food, cosmetics and pharmaceutical industries (Nielsen et al. 2016; FMI 2020). With an acceptable daily intake of 25 mg/kg BW/day (Aguilar et al. 2015), EFSA allows it’s use in products like sauces, soups, chewing gum, coconut milk, dairy products and usually at maximal concentrations of 10-10000 mg/kg, depending on the product ^4^ (European Commision n.d.). Also the U.S. Food and Drug Administration (FDA) limits the use of P80 to 4 – 10000 mg/kg, depending on the product ^5^ (FDA 2019a).

Soy lecithin is a mixture of phospoholipids (at least 60%), triglycerides, sterols and carbohydrates obtained by extraction from soybeans. It is more widely used than CMC and P80, primarily in bakery products, ice creams, chocolate etc ^6^ (European Commision 2018). Lecithins are allowed by EFSA in most food applications in *quantum satis* levels, and also the FDA considers soy lecithin a GRAS compound^7^ (Carocho et al. 2014; FDA 2019c). Even though soy lecithin is considered safe or even beneficial for health (Ehehalt et al. 2010; Mourad et al. 2010; Stremmel et al. 2010), the impact of soy lecithin on the gut microbiota has never been studied. Since this compound is one of the most extensively used food emulsifiers, it was incorporated in this research.

Finally, rhamnolipids and sophorolipids are two biotechnological emulsifiers of microbial origin. Due to their advantageous properties with respect to (eco)toxicity and waste stream reuse (Van Bogaert, Zhang, and Soetaert 2011; Haba et al. 2003; Costa et al. 2017) they are currently under consideration as novel food additives (Costa et al. 2017; Nitschke and Silva 2018; Cameotra and Makkar 2004). Their strong emulsifying capacities (Van Bogaert, Zhang, and Soetaert 2011; Costa et al. 2017) and more natural origin could qualify them as adequate alternatives for emulsifiers of chemical origin, which the food industry is currently seeking to replace under the umbrella of the ‘clean label’ trend (Asioli et al. 2017; Nitschke and Silva 2018; Costa et al. 2017). However, given their strong antimicrobial properties, an evaluation at the level of the gut microbiota is highly warranted before such applications can be legalized.

Here we investigated the effects of the five above-mentioned dietary emulsifiers on human fecal microbiota through 48h *in vitro* batch incubations. This set of emulsifiers enables the comparison of the previously-studied chemical emulsifiers with the natural extract lecithin and with biosurfactants. In order to take into account interindividual variability in microbiome composition as a possible determinant of the putative impact from emulsifiers, we separately assessed microbial incubations from ten different individuals.

## 2 Materials and Methods

### 2.1 Experimental design

Fecal material from 10 human individuals was collected and separately incubated for 48h with the five emulsifiers at three concentrations (0.005% (m/v), 0.05% (m/v) and 0.5% (m/v)). Emulsifier concentrations were chosen based on the maximal legal concentration in food products (EFSA & FDA), which comply with commonly applied concentrations in food products (Msagati 2012; Mallet 1992; Adams et al. 2004). Each donor incubation series also included a control condition, in which a sham treatment with an equivalent volume of distilled water was performed.

The emulsifiers used during this study were sodium carboxymethylcellulose (CMC), polysorbate (P80), soy lecithin, sophorolipids and rhamnolipids. CMC (catalogue number 419303 – average molecular weight of 250 000 g/mol, degree of substitution of 0.9), P80 (P4780 - suitable for cell culture) and rhamnolipids (RL) (R90 - 90% pure) were obtained from Sigma Aldrich, St. Louis, MO. Soy lecithin was obtained from Barentz Unilecithin (UNILEC – ISL non GMO IP) and sophorolipids were obtained from the UGent Inbio group from the Centre for Synthetic Biology. The latter were described as 75% (w/v) solutions and their composition was determined to be mainly lactonic, diacetylated C18:1 SL.

Donors 2 and 6 reported to follow a vegetarian and vegan diet. All other donors consumed an omnivorous diet. The age of the four female and six male donors varied from 23 to 53 years old. None of the donors received any antibiotic treatment in the 3 months prior to their donation. Experimental work with fecal microbiota from human origin was approved by the ethical committee of the Ghent University hospital under registration number BE670201836318.

### 2.2 Batch incubation

Before incubation, the five emulsifiers were supplemented to amber penicillin bottles containing 40 mL of autoclaved low-sugar nutritional medium (per L: 0.25 g gum arabic, 0.5 g pectin,0.25 g xylan, 1 g starch, 3 g yeast extract, 1 g proteose peptone, 2 g pig gastric mucin; all from Sigma Aldrich, St. Louis, MO). The amounts of emulsifiers to add were calculated to obtain concentrations of 0.005% (m/v), 0.05% (m/v) and 0.5% (m/v) in a final volume of 50 mL (the volume obtained after addition of the fecal slurry). The bottles were stored in a 4°C fridge until use (for maximum 3 days).

At the start of the batch experiments, the penicillin flasks containing nutritional medium and emulsifiers were brought to room temperature to provide an ideal growth environment for the fecal bacteria. Fresh fecal samples were then collected in airtight plastic lidded containers. Anaerogen™ sachets (Oxoid Ltd., Basingstoke, Hampshire, UK) were used to sequester O_2_. The samples were stored at 4°C until use for a maximum of 3 hours. A fecal inoculum was then prepared as described in De Boever, Deplancke, and Verstraete (2000), by mixing 20% (w/v) fecal material into a 0.1 M anaerobic phosphate buffer (pH 6.8) supplemented with 1 g/L sodium thioglycanate as a reducing agent. Into each penicillin bottle, 10 mL of fecal inoculum was added, after which the bottles were closed with butyl rubber stoppers and aluminium caps. The headspaces were flushed with a N_2_/CO_2_ (80/20)-gas mixture using gas exchange equipment to obtain anaerobic conditions and incubated in an IKA® KS 4000 I Control shaker at 200 rpm at 37°C. During the course of the experiments, the pH was followed up every day with a Prosense QP108X pH-electrode connected to a Consort C3020 multi parameter analyzer to ensure stable and viable growth conditions (pH remained within 5.5 – 6.8).

Aliquots were taken on three timepoints: immediately after the start of the incubation (T0) (2-3 hours after combining the fecal inoculum with the medium containing emulsifier), after 24 hours of incubation (T1) and after 48 hours of incubation (T2). Samples were taken for short chain fatty acid (SCFA-) analysis (1 mL), 16S rRNA gene amplicon sequencing (1 mL), for flagellin detection (500 µL) and for immediate fluorescent cell staining and flow cytometry (100 µL). SCFA-, flagellin- and sequencing samples were stored at −20°C until analysis.

### 2.3 Intact/damaged cell counts

To assess the impact of the emulsifiers on total and intact cell concentrations, cell staining with SYBR® green and propidium iodide was performed after which cells were counted on an Accuri C6+ Flow cytometer from BD Biosciences Europe. The combination of these two cell stains is frequently used to distinguish intact bacterial cells from cells damaged at the level of the cell membrane, since SYBR® green enters any cell rapidly, while propidium iodide, being a larger molecule, enters intact cells much slower and mainly stains damaged cells within commonly applied incubation times (Van Nevel et al. 2013). Samples were analyzed immediately after sampling to preserve the intact cell community. Dilutions up to 10^−4^ and 10^−5^ were prepared in 96-well plates using 0.22 µm filtered 0.01 M phosphate buffered saline (PBS) (HPO_4_^2-^ / H_2_PO_4_^-^, 0.0027 M KCl and 0.137 M NaCl, pH 7.4, at 25 °C) and these were subsequently stained with SYBR® green combined with propidium iodide (SGPI, 100x concentrate SYBR® Green I, Invitrogen, and 50x 20 mM propidium iodide, Invitrogen, in 0.22 μm-filtered dimethyl sulfoxide) (Van Nevel et al. 2013; Props et al. 2016a). After 25 minutes of incubation, the intact and damaged cell populations were measured immediately with the flow cytometer, which was equipped with four fluorescence detectors (530/30 nm, 585/40 nm, >670 nm and 675/25 nm), two scatter detectors and a 20 mW 488 nm laser. The flow cytometer was operated with Milli-Q (Merck Millipore, Belgium) as sheath fluid. The blue laser (488 nm) was used for the excitation of the stains and a minimum of 10,000 cells per sample were measured for accurate quantification. Settings used were an FLH-1 limit of 1000, a measurement volume of 25 µL and the measurement speed was set to ‘fast’. Cell counts were obtained by gating the intact and damaged cell populations in R (version 3.6.2) according to the Phenoflow-package (v1.1.6) (Props et al. 2016b). Gates were verified using data from negative control samples (only 0.22 µm filtered 0.01 M PBS) (Figure S1).

### 2.4 SCFA-analysis

The SCFA-concentrations were determined by means of diethyl ether extraction and capillary gas chromatography coupled to a flame ionization detector as described by De Paepe et al. (2017); Anderson et al. (2017). Briefly, 1 mL aliquots were diluted 2x with 1 mL milli-Q water and SCFA were extracted by adding approximately 400 mg NaCl, 0.5 mL concentrated H_2_SO_4_, 400 µL of 2-methyl hexanoic acid internal standard and 2 mL of diethyl ether before mixing for 2 min in a rotator and centrifuging at 3000 *g* for 3 minutes. Upper layers were collected and measured using a GC-2014 capillary gas chromatograph (Shimadzu, Hertogenbosch, the Netherlands), equipped with a capillary fatty acid-free EC-1000 Econo-Cap column (Alltech, Lexington, KY, US), 25 m × 0.53 mm; film thickness 1.2 μm, and coupled to a flame ionization detector and split injector. One sample (donor 9, timepoint 2 – 0.05% (m/v) CMC) returned only zero values, presumably due to a technical error. This sample was therefore omitted prior to computational analyses.

### 2.5 Amplicon sequencing

Samples from T0 and T2 were selected for Illumina 16S rRNA gene amplicon sequencing. The samples (1 mL) were first centrifuged for 5 min at 30 130 *g* in an Eppendorf 5430 R centrifuge to obtain a cell pellet. After removing the supernatant, the pellets were subjected to DNA-extraction (Vilchez-Vargas et al. 2013; De Paepe et al. 2017). The pellets were dissolved in 1 mL Tris/HCl (100 mM, pH = 8.0) supplemented with 100 mM EDTA, 100 mM NaCl, 1% (w/v) polyvinylpyrrolidone and 2% (w/v) sodium dodecyl sulfate after which 200 mg glass beads (0.11 mm Sartorius, Gottingen, Germany) were added and the cells were lysed for 5 min at 2000 rpm in a FastPrep VR-96 instrument (MP Biomedicals, Santa Ana, CA). The beads were then precipitated by centrifugation for 5 min at 30 130 *g* and the supernatant was collected. Purification of DNA took place by extraction of cellular proteins with 500 µL phenol-chloroform-isoamilyc alcohol 25-24-1 at pH7 and 700 µL 100% chloroform. The DNA was precipitated by adding 1 volume of ice-cold isopropyl alcohol and 45 µL sodium acetate and cooling for at least 1h at −20°C. Isopropyl alcohol was then separated from DNA by centrifugation for 30 min at 4°C and at 30 130 *g* and the pellet was dried by pouring off the supernatant. It was resuspended in 100 mL 1x TE buffer (10 mM Tris, 1 mM EDTA) for storage at −20°C.

DNA-quality was verified by electrophoresis in a 1.5% (w/v) agarose gel (Life technologies, Madrid, Spain) and DNA-concentration was determined using a QuantiFluor® dsDNA kit (detection limit: 50 pg/mL; sensitivity: 0.01 – 200 ng/µL) and Glomax® –Multi+ system (Promega, Madison, WI) with the blue fluorescence optical kit installed (Ex: 490nm, Em: 510–570 nm).

Library preparation and next generation 16S rRNA gene amplicon sequencing were performed at the VIB nucleomics core (VIB, Gasthuisberg Campus, Leuven, Belgium) as described in Tito et al. (2017). The V4 region of the 16S rRNA gene was amplified using the bacterial 515F (GTGYCAGCMGCCGCGGTAA) and the 806R (GGACTACNVGGGTWTCTAAT) primers, which were modified with both Illumina adapters as well as adapters for directional sequencing. Sequencing was then performed on an Illumina MiSeq platform (Illumina, Hayward, CA, USA) according to manufacturer’s guidelines.

One sample (donor 3, timepoint 2, 0.05% (m/v) sophorolipids) failed to sequence. The sequencing data have been submitted to the National Center for Biotechnology Information (NCBI) database under the accession number PRJNA630547.

Processing of amplicon data was carried out using mothur software version 1.40.5 and guidelines (Kozich et al. 2013). First, contigs were assembled, resulting in 14 977 727 sequences, and ambiguous base calls were removed. Sequences with a length of 291 or 292 nucleotides were then aligned to the silva_seed nr.123 database, trimmed between position 11 895 and 25 318 (Quast et al. 2013). After removing sequences containing homopolymers longer than 9 base pairs, 92% of the sequences were retained resulting in 2 957 626 unique sequences. A preclustering step was then performed, allowing only three differences between sequences clustered together and chimera.vsearch was used to remove chimeras, retaining 79% of the sequences. The sequences were then classified using a naïve Bayesian classifier against the Ribosomal Database Project (RDP) 16S rRNA gene training set version 16, with a cut-off of 85% for the pseudobootstrap confidence score. Sequences that were classified as *Archaea, Eukaryota, Chloroplasts*, unknown or *Mitochondria* at the kingdom level were removed. Finally, sequences were split at the order level into taxonomic groups using the opticlust method with a cut-off of 0.03. The data was classified at a 3% dissimilarity level into OTUs resulting in a .shared (count table) and a .tax file (taxonomic classification).

For the entire dataset of 319 samples, 95 511 OTUs were detected in 175 genera. An OTU was in this manuscript defined as a collection of sequences with a length between 291 and 292 nucleotides and with 97% or more similarity to each other in the V4 region of their 16S rRNA gene after applying hierarchical clustering.

### 2.6 Cell culture for flagellin detection

Murine TLR5-expressing HEK 293 cells (InvivoGen), which are designed to respond to bacterial flagellin in cell culture medium, were cultured according to the manufacturer’s guidelines. Cells were grown from an in house created frozen stock in Dulbecco’s modified Eagle’s (DMEM) growth medium (4.5 g/L glucose, 10% fetal bovine serum, 50 U/mL penicillin, 50 µg/mL streptomycin, 2 mM L-glutamine) supplemented with 100 µg/mL Normocin™ and maintained in culture in DMEM growth medium supplemented with 100 µg/mL Normocin™, 10 µg/mL of blasticidin and 100 µg/mL of zeocin™. Medium was refreshed every two days and cells were passaged when reaching 70-80% confluency.

Assays for flagellin detection were performed as instructed by Invivogen, using cells from passage 4-9. Samples from donor 3, 5 and 7 were selected for this assay based on a high, intermediate and low metabolic response to the emulsifiers, as measured by the SCFA-levels. Before combining with the HEK-blue cells, the samples were purified to obtain only the bacterial cells by first diluting them 1/4 in UltraPure™ DNase/RNase-Free distilled water (Invitrogen), then centrifuging twice at 4 226 *g* for 10 min, with a washing step using 0.22 µm filtered -/- PBS in between. The resulting cell pellet was dissolved into 0.22 µm filtered -/- PBS. A standard curve (125 ng/mL – 1.95 ng/mL), prepared from recombinant flagellin from *Salmonella typhymurium* (RecFLA-ST, Invivogen) in sterile water was also added to the plate in triplicate. After an incubation for 23h, absorbances were obtained using a Tecan Infinite F50 plate reader at 620 nm.

As a check for the viability of the cell culture after combination with the samples, a resazurin assay was performed. To this end, the supernatant from the cell culture plate used for the flagellin assay was discarded after the first incubation phase. The cells were then washed using 0.22 µm filtered -/- PBS. As a positive control, 3 wells were spiked with 20 µL of a 5% Triton solution. The rest of the wells received 20 µL -/- PBS after which 180 µL of a of 0,01 mg/mL resazurin solution was added to all wells. After 3 hours of incubation at 37°C and 10% CO_2_, cell activity was measured using a Glomax®-Multi1 system (Promega, Madison, WI) with filter the green fluorescence optical kit (Ex: 525 nm, Em: 580–640 nm).

### 2.7 Data analysis and statistics

Data visualization and processing was performed in R version 3.4.2 (2017-09-28) (R Core Team, 2016) and Excel 2016. All hypothesis testing was done based on a significance level of 5% (α = 0.05).

#### 2.7.1 Cell counts and SCFA

After loading the cell count table in R, total cell counts were calculated as the sum of the intact and damaged cell counts. The data were explored by calculating intact/damaged cell count ratio’s and percentages of intact cells at different timepoints. For plotting, both the 10 000x and 100 000x dilutions were taken into account. Boxplots of total and intact cell counts, as well as intact/damaged ratio’s were created using ggplot2 (v3.2.1) in which the stat_compare_means function was used to check the significance of the emulsifier effect, by means of a Kruskal-Wallis test.

Statistical analysis of SCFA-levels was similar to that of the cell counts. Production levels of acetate, propionate and butyrate over 48h (C_T2_– C_T0_) were first calculated and then boxplots were created using ggplot2. Significance of the effect of the emulsifier treatment was tested with stat_compare_means using Wilcoxon Rank Sum tests with Holm correction and Kruskal-Wallis test for overall group comparison.

#### 2.7.2 Amplicon sequencing data

The shared and taxonomy files resulting from the mothur pipeline were loaded into R for further processing. Singletons (OTUs occurring only once over all samples) were removed, resulting in 36 496 OTUs being retained (McMurdie and Holmes 2014). Rarefaction curves were created to evaluate the sequencing depth (Figure S2) (Oksanen et al. n.d.). As the number of 16S rRNA gene copies present within bacteria differs between species, a copy number correction of the reads was carried out by first classifying the representative sequences of the OTUs (also obtained via the mothur pipeline) using the online RDP classifier tool, then obtaining both a copy number corrected read classification and a non copy number corrected one, calculating the copy number by dividing both and finally dividing the acquired read counts in the shared file by the calculated copy numbers.

Both relative and absolute abundances of the OTUs and genera were calculated from the copy number corrected read counts and were explored via bar plots using ggplot2 (v3.2.1). Relative abundances were calculated as percentages of the total read counts per sample. Absolute abundances were calculated (Quantitative microbial profiling) by multiplying the total cell counts obtained via flow cytometry with the relative abundances of the OTUs (similar to Vandeputte et al. 2017).

Overall community composition was visualized using Principle Coordinate Analysis (PcoA) on the abundance based Jaccard distance matrix using the cmdscale-function in the stats (v3.6.2) package. To investigate the effects of the individual contraints on the microbial community a series of distance based redundancy analyses (dbRDA) was then performed on the scores obtained in the PCoA on the Jaccard distance matrix using the capscale function in the vegan (v2.5-6) package. Permutation tests were used to evaluate the significance of the models and of the explanatory variables (De Paepe et al. 2018). The global model included the factors Emulsifier, Emulsifier concentration, Timepoint and Donor as explanatory variables and the absolute abundances of the genera as explanatory variables. In a first dbRDA this full model was included, to investigate the share of variance explained by each constraint variable. The timepoint factor was distinguished as the factor causing the largest share of variance and since its effect was of little interest to us it was partialled out in further dbRDAs. To check for the effect of the donor variable on the microbial community, a second and third dbRDA were performed, with and without conditioning of the donor variable. The final model then visualized the effects of the treatments (defined by factors emulsifier and emulsifier concentration). The results of the dbRDAs were plotted as type II scaling correlation triplots showing the two first constrained canonical axes (labelled as dbRDA Dim 1/2) and the proportional constrained eigenvalues representing their contribution to the total (both constrained and unconstrained) variance.

The chao1, chaobunge2002, ACE-1, Shannon, Simpson, InvSimpson and Pielou diversity indices were calculated for the microbial community after 48h incubation based on the copy number corrected OTU-table using the SPECIES (v1.0) package and the diversity function in the vegan (v2.5-6) package. Indices were plotted using ggplot2 and significances were tested using pairwise Wilcoxon Rank Sum tests with Holm correction (ggpubr package v.0.2.4).

To evaluate differential abundance of genera between emulsifier treatments and control, the DESeq2 package (v 1.24.0) was applied on the copy number corrected count-table at genus level. In order to streamline the DESeq-analysis, pre-filtering according to McMurdie and Holmes 2014 was first applied on the copy number corrected count-table, after which a genus-level table was created using the aggregate function (stats package v3.6.3). In the generalized linear model, the factor Timepoint, Donor and Treatment – a concatenation of the emulsifiers and their concentrations – were included. A likelihood ratio test was employed within the DESeq function on the reduced model, containing only the factors Donor and Timepoint, to test for the significance of the model. Low count genera were subjected to an empirical Bayesian correction (Love, Huber, and Anders 2014). For pairwise comparison of treatments versus controls, Wald tests were used after shrinkage of the Log2FoldChange values by means of the lfcShrink function. P-values were adjusted by means of a Benjamini-Hochberg procedure (Love, Huber, and Anders 2014). Results were visualized in volcanoplots, displaying the – log(adjusted p-value) versus the Log2FoldChange of each genus. Additionally, box plots were created showing the log-transformed pseudocounts extracted by the plotCounts function for each genus that showed significant differential abundance. Since for CMC and P80 no significantly altered genera were found, these emulsifiers were omitted from the boxplots.

Finally, to summarize relations between the emulsifier treatments and the intact cell counts, the SCFA-data and the 16S rRNA sequencing data, a partial redundancy analysis was carried out performed using the rda function in the vegan package (v2.5.6). The acetate, propionate and butyrate concentrations, the intact cell counts and the relative abundance of the genera as response variables and the factors Emulsifier, Emulsifier concentration, Donor and Timepoint as explanatory variables. Since the response variables carried different units, they were first centered around their mean using the scale function (base R v3.6.2). The factors Donor and Timepoint were partialled out to visualize solely the effect of the emulsifier treatments. The statistical significance of the effects was tested via a permutation tests and the results were plotted in a type II correlation triplot showing the first two constrained canonical axes (RDA1/2) annotated with their proporiotnal eigenvalues representing their contribution to the constrained variance. The sites were calculated as weighed sums of the scores of the response variables.

#### 2.7.3 Metagenome prediction

Indications of functionality from phylogenetic information were obtained using PICRUSt (Phylogenetic Investigation of Communities by Reconstruction of Unobserved States) (Langille et al. 2013). An OTU-table was first generated against the Greengenes reference database (v13.8) using a closed ref OTU-picking strategy. The obtained OTU-table was then run through PICRUSt’s normalize_by_copy_number.py script (Langille et al. 2013), which divides the abundance of each OTU by its inferred 16S copy number (the copy number is inferred from the closest genome representative for a 16S Greengenes reference sequence). The metagenome was then predicted using Kyoto Encyclopedia of Genes and Genomes (KEGG) database (Kanehisa et al. 2012). The prediction provided an annotated table of predicted gene family counts for each sample, where gene families were grouped by KEGG Orthology identifiers. Significantly different L2-level pathways across emulsifier concentration were visualized in boxplots using the ggplot2 package (v3.2.1). Kruskal-Wallis tests were performed for overall comparison of emulsifier concentrations within L2-pathways for each emulsifier and Wilcoxon rank sum tests were used for pairwise comparison of emulsifier concentrations *vs* control. Also relying on PICRUSt (Langille et al. 2013), the BugBase tool (Ward et al. 2017) was used to determine the relative degree of biofilm formation, oxygen utilization, pathogenic potential, oxidative stress tolerance and Gram-stain of the bacteria in the samples.

#### 2.7.4 Flagellin concentration

Initial data processing was executed in Excel 2016. First, a four parametric logistic model was fitted to the standard curve using the 4PL-Curve Calculator from aatbio.com (AAT Bioquest n.d.). Given our observation that the emulsifiers decreased HEK cell activity, flagellin concentrations were normalized by use of absorbance values obtained from the resazurin assays: flagellin concentrations were divided by the ratio of the absorbance values from the samples over the average absorbance values for the standard curve of the same plate. Graphs were created using ggplot2. Since donors were observed separately and no replicate experiments per donor were performed, no statistical tests were performed for the flagellin data.

### 2.8 Donor diversity analysis

We sought to assess the overall susceptibility of the 10 donors to the effects of the emulsifiers. Since literature describes no workflow for this purpose, we elaborated our own. Donors were ranked in terms of their susceptibility to the emulsifiers using several parameters: The 48h production of the three most abundant SCFA (acetate, propionate and butyrate), the intact cell counts at T2 and the relative and absolute abundance of the most abundant OTU in the OTU -table, *Escherichia/Shigella*, at T2. These calculations were performed in Excel 2016.

First, to correct for the batch-effect, the control-values were subtracted from the treatment-values for each donor. Next the corrected treatment values were summated for each donor, to obtain a single value that expressed the donor’s susceptibility and these values were then used to rank the donors from least to most susceptible for every parameter, visualized in bar graphs. This workflow was followed for each parameter.

### 2.9 Comparison of equivalent emulsifier concentrations

Due to their stronger emulsifying properties, rhamnolipids and sophorolipids could reportedly be used in lower concentrations in food products than conventional chemical emulsifiers (Nitschke and Silva 2018). Therefore, we sought to compare the effects of the chemical emulsifiers, CMC and P80, versus those of the biosurfactants, rhamnolipids and sophorolipids, with regards to their impacts on the gut microbiota at equivalent emulsifying concentrations. Wilcoxon Rank Sum test were executed in R using the compare_means function on the same parameters we used to evaluate donor diversity (see 3.7). As equivalent concentrations we considered a 10x lower concentration of biosurfactants compared to the chemical emulsifiers, given that this is what industry reports (Van Haesendonck and Vanzeveren 2006). Hence we compared the condition of 0.5% (m/v) of chemical emulsifiers with that of 0.05% (m/v) of biosurfactancs and 0.05% (m/v) of chemical emulsifiers with 0.005% (m/v) of biosurfactants.

## 3 Results

### 3.1 Community structure is altered by addition of dietary emulsifiers

#### 3.1.1 Intact/damaged cell counts

Analysis of intact and damaged cell populations with flow cytometry (SGPI-staining defines damage at the level of the cell membrane (Wlodkovic, Skommer, and Darzynkiewicz 2009; Buysschaert et al. 2018; Falcioni, Papa, and Gasol 2008), was used as a proxy for emulsifier toxicity. First, total and intact cell counts in the controls dropped by 14% ± 2% and 21% ± 3%, respectively, after 48h *in vitro* incubation (Table S1 and S2). When considering the impact of the emulsifiers, we observed that higher concentrations of rhamnolipids, sophorolipids and soy lecithin resulted in significantly lower total and intact cell counts (Figure 1 and 2). At 0.5% (m/v) rhamnolipids, intact cells decreased by 91% ± 0% after 48h compared to the control sample of T0. At 0.5% (m/v) sophorolipids, about 89% ± 1%, was lost and at 0.5% (m/v) soy lecithin, 87% ± 1% was lost. The toxic effects were immediate for the sophorolipids and rhamnolipids, while for soy lecithin this decreasing effect only became significantly apparent after 24h (T1) (Figure 2). The impact of CMC and P80 towards the cell population was less pronounced. CMC even increased the total cell counts (not significantly) at higher concentrations, although the fraction of living cells remained unaffected for all CMC-conditions.

**Figure 1:**
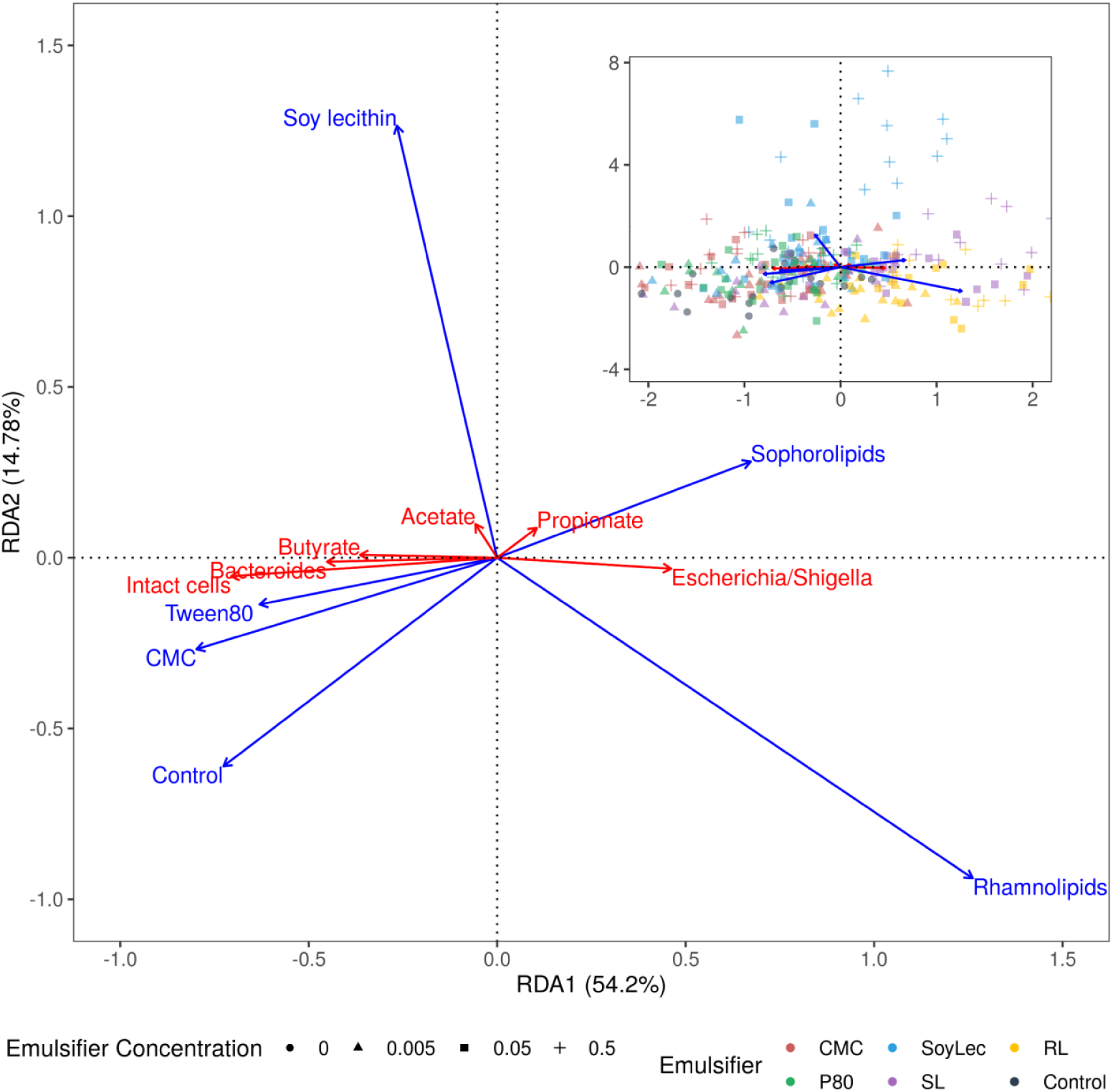
Type II scaling triplot obtained using partial redundancy analysis of the microbial community detected after 48h of in vitro batch incubations of fecal material from 10 human donors in sugar depleted medium supplemented with 5 emulsifiers at 4 concentrations. In the main figure, the intact cell count, the SCFA-levels and the relative abundance of the top two genera are shown as response variables (red arrows) and emulsifier and emulsifier concentration are given as explanatory variables (blue arrows). The top right figure also displays the samples as sites.

**Figure 2:**
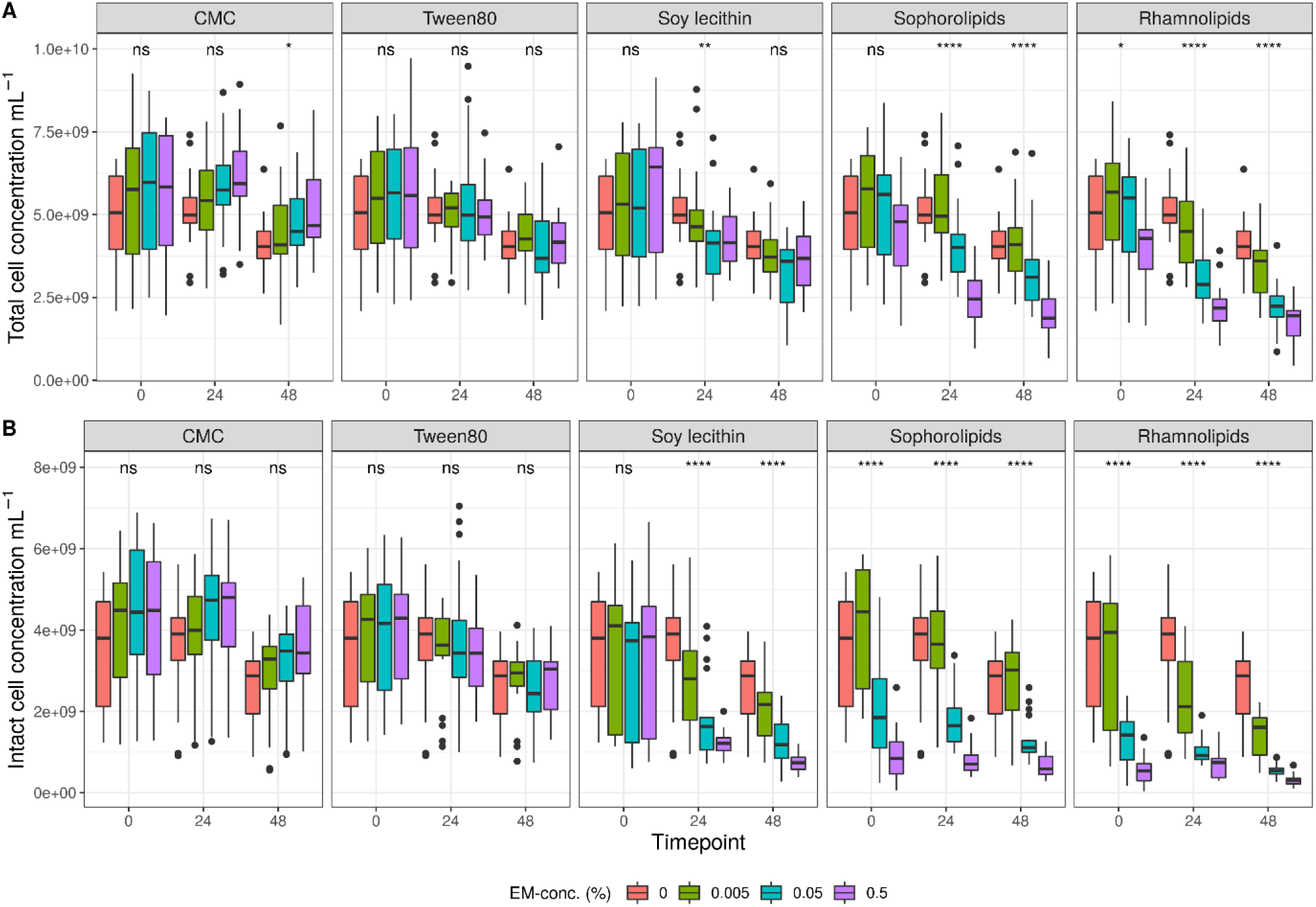
Average total (A) and intact (B) bacterial cell counts (cells/mL) detected during in vitro batch incubations of fecal material from 10 donors with sugar depleted medium supplemented with 5 emulsifiers at 4 concentrations. Samples were taken upon incubation (T0; 2-3h after inoculation) as well as after 24h (T1) and 48h (T2) of incubation. Asterisks indicate significant differences detected with a Kruskal Wallis test (α = 0.05).

#### 3.1.2 Microbial community

The impact from the emulsifiers towards microbial community structure was assessed with 16S rRNA gene amplicon sequencing. First, the *in vitro* conditions had an impact on microbiota composition. While each donor showed a unique profile of microbial genera at the start of the experiment more similar microbial community profiles were obtained upon incubation, primarily due to an increase in *Escherichia/Shigella* abundance from 0.02% ± 0.02% to 16% ± 25% (Fig S3 – S5).

Independent from the *in vitro* effects, clear differences were noted between emulsifier treatments and controls, which were both emulsifier- and donor-dependent (Figure 3, Figure S4 and S5). Where the effects of rhamnolipids and sophorolipids were most outspoken, the impact of soy lecithin was intermediary and CMC and P80 had the smallest impacts (Figure 1). This was evidenced by significant drops in diversity indices upon incubation with rhamnolipids, sophorolipids and to a lesser extent soy lecithin (Figure 4). DESeq-analysis further revealed significant differential relative abundance of 36 genera, from which 23 were increased and 13 were suppressed, compared to the control condition (Figure 5 and 6). Rhamnolipids triggered the strongest changes, with the three most increased genera being unclassified *Enterobacteriaceae* (L2FC = 3.85; Padj <0.0001), *Fusobacterium* (L2FC = 2.75; Padj < 0.0001) and *Escherichia*/*Shigella* (L2FC = 2.49; Padj <0.0001) and the three most suppressed ones being unclassified *Bacteroidetes* (L2FC = −2.19; Padj = < 6,323E-4), *Barnesiella* (L2FC = −2.09; Padj <0,009) and *Bacteroides* (L2FC = −2.02; Padj <0,002). The top three most increased genera for sophorolipids were *Escherichia*/*Shigella* (L2FC = 1.86; Padj <0,043), *Acidaminococcus* (L2FC = 1.80; Padj <0.0001) and *Phascolarctobacterium* (L2FC = 1.68; Padj = <0.0001) and the three most decreased were unclassified *Bacteroidetes* (L2FC = −1.97; Padj <0.0001), *Barnesiella* (L2FC = −1.70; Padj = <0.0001) and *Bacteroides* (L2FC = −1.53; Padj = 3,034E-06). The top three most increased genera by soy lecithin were *Acidaminococcus* (L2FC = 1.23; Padj =0,016), *Porphyromonadaceae*_*unclassified* (L2FC = 1.19; Padj =0,017) and *Sutterella* (L2FC = 1.19; Padj = 0,004). Two significantly decreased genera were *Flavonifractor* (L2FC = −1.04; Padj = 0,009) and *Pseudoflavonifractor* (L2FC = −0.95; Padj = 0,015) (Figure 5 and 6).

**Figure 3:**
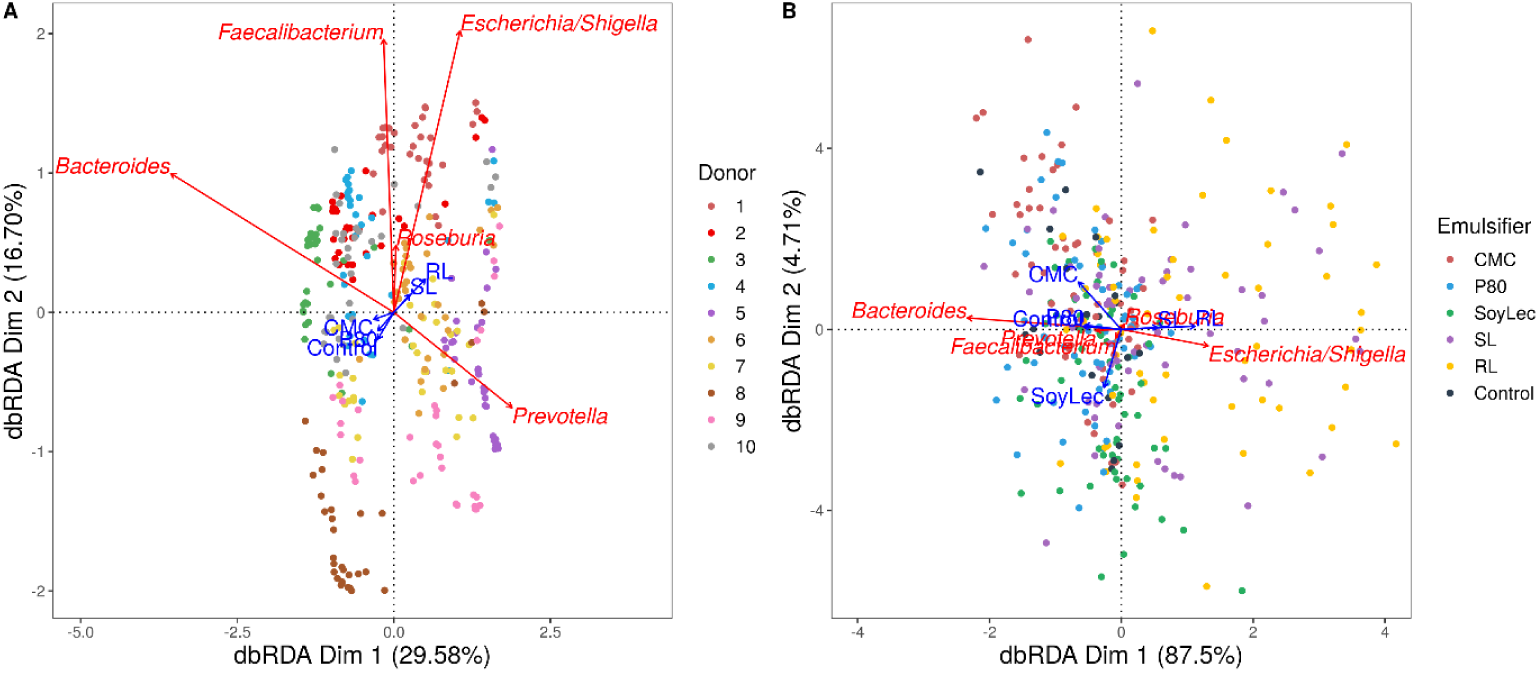
Type II scaling triplots obtained using partial distance based redundancy analysis of the microbial community composition detected using 16S rRNA gene amplicon sequencing after 48h of in vitro batch incubations of fecal material from 10 donors with sugar depleted medium supplemented with 5 emulsifiers at 4 concentrations. Samples were taken upon incubation (T0; 2-3h after inoculation) as well as after 24h (T1) and 48h (T2) of incubation. Factors Donor, Emulsifier and Emulsifier concentration were set as explanatory variables (blue arrows) and absolute abundances of genera as response variables (red arrows). Only the top five genera were displayed for adequate visibility. A: The factor timepoint was partialled out. B: The factors donor and timepoint were partialled out.

**Figure 4:**
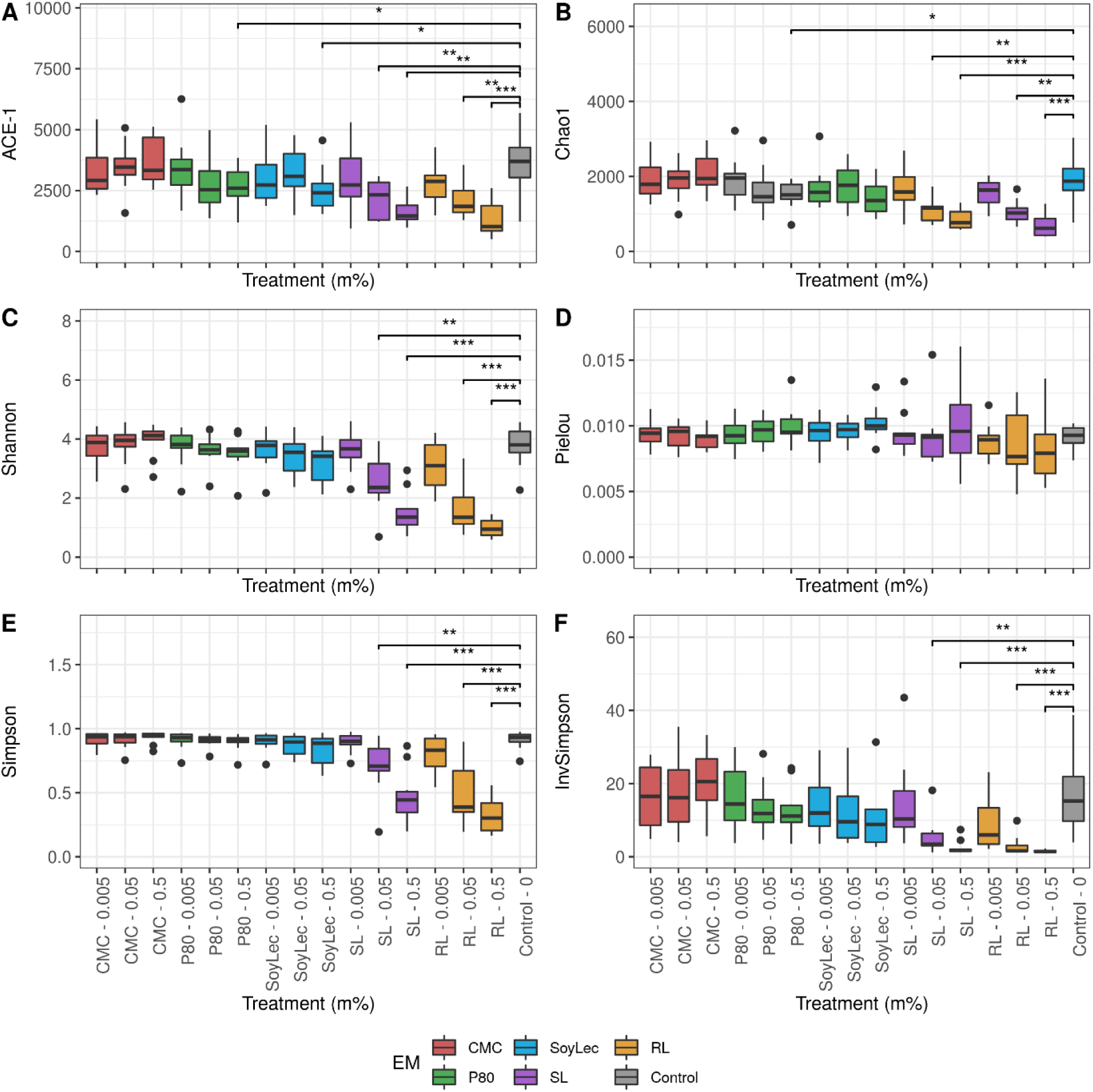
Diversity parameters of gut microbial community obtained after 48h of in vitro batch incubations of fecal material from 10 donors with sugar depleted medium supplemented with 5 emulsifiers at 4 concentrations. Samples were taken upon incubation (T0; 2-3h after inoculation) as well as after 24h (T1) and 48h (T2) of incubation. Asterisks represent significant differences with control based on Wilcoxon Rank Sum tests with Holm correction (α = 0.05).

**Figure 5:**
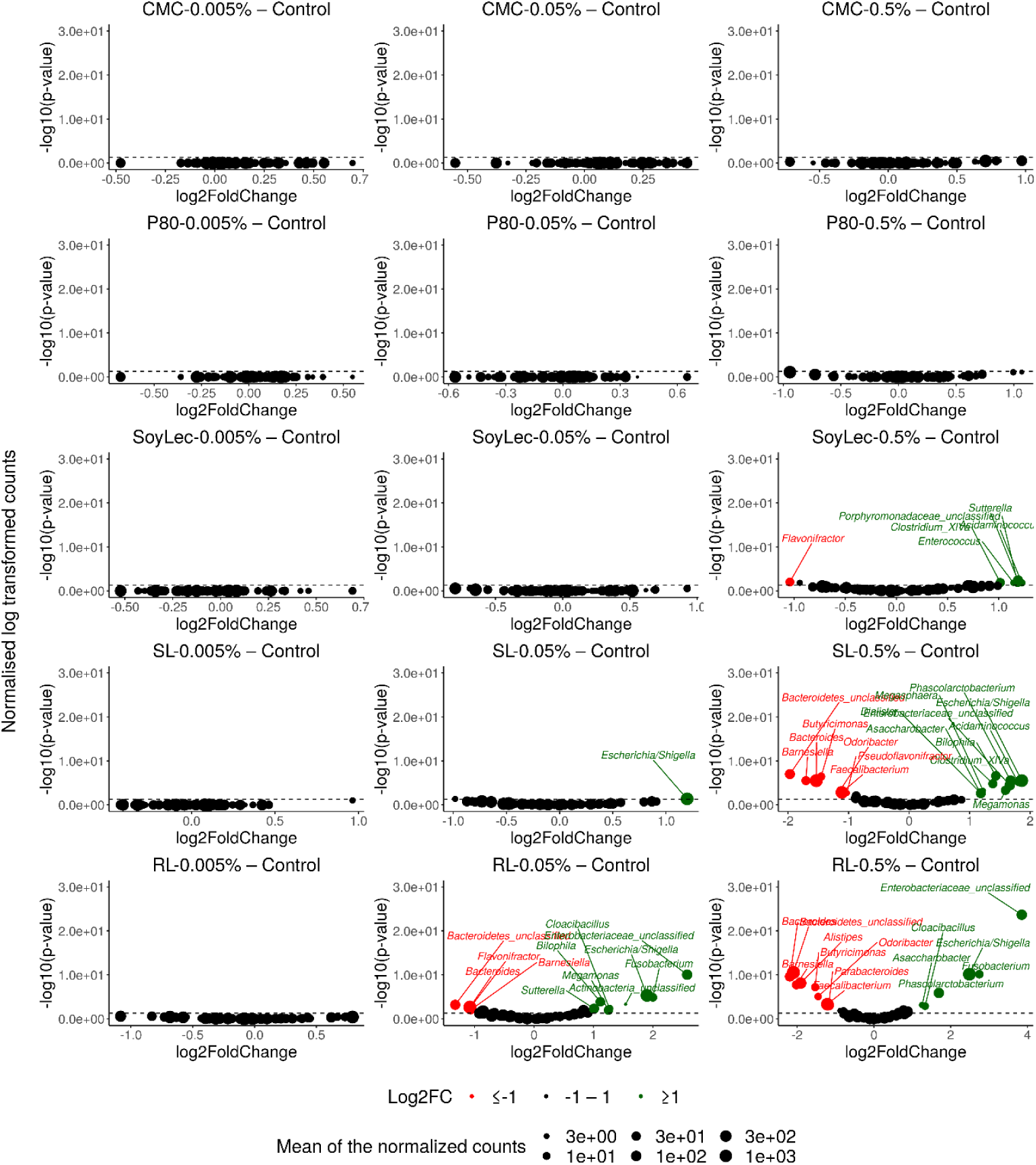
Volcanoplots indicating gut microbial community alterations after 48h of in vitro batch incubations of fecal material from 10 donors with sugar depleted medium supplemented with 5 emulsifiers at 4 concentrations. Samples were taken upon incubation (T0; 2-3h after inoculation) as well as after 24h (T1) and 48h (T2) of incubation. Log2FoldChange (L2FC) of genus abundances for all emulsifier treatments versus the control are presented on the x-axis and the log transformed adjusted p-value is presented on the y-axis. Significantly increased or decreased genera are indicated in respectively green and red. The dashed line represents the significance threshold of α = 0.05.

**Figure 6:**
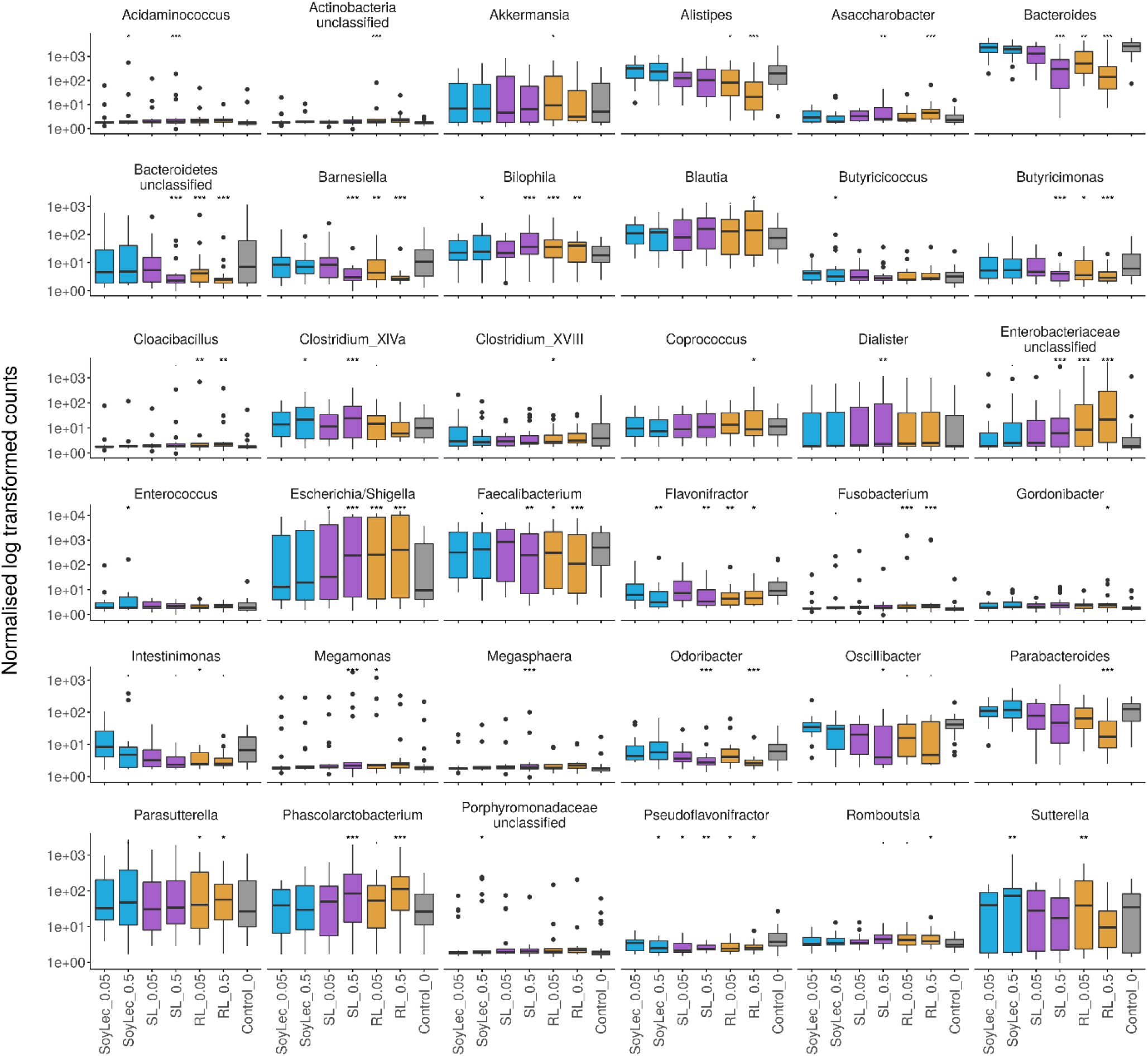
Copy number corrected counts of significantly in- or decreased genera, obtained from DESeq analysis in R (version 3.4.2), after 48h of exposure of gut microbial communities from 10 donors to soy lecithin, sophorolipids and rhamnolipids during in vitro batch incubations. Asterisks represent significant differences with the control based on Wald tests (α = 0.05).

### 3.2 Functional analysis

#### 3.2.1 SCFA

Short chain fatty acids were analyzed to study how exposure to dietary emulsifiers affects the general microbial metabolic activity. We observed that SCFA-production was significantly and differently affected by rhamnolipids, sophorolipids and soy lecithin, while no changes were observed for P80 and CMC (Figure 1 and 7). The strongest impacts were noted for rhamnolipids, which, at 0.5% (m/v), significantly decreased total SCFA production by about 36% ± 5% (P_Wilcox_ <0.0001) compared to the control condition. This decrease was mainly attributed to a 32% ± 7% decrease in acetate production (P_Wilcox_ <0.0001) compared to the control. Rhamnolipids at 0.5% (m/v) also reduced butyrate production by 96% ± 6% compared to the control condition (P_Wilcox_ <0.0001) while propionate production remained unaffected. Interestingly, incubation with 0.5% (m/v) sophorolipids also resulted in a decrease in butyrate production by 73% ± 24% compared to the control (P_Wilcox_ <0.0001), while propionate production increased by 88% ± 50% (P_Wilcox_ = 2.1 e-04). Soy lecithin at 0.5% (m/v) significantly increased propionate production by 29% ± 18% on average (P_Wilcox_ = 0.0089) and decreased butyrate production non-significantly by 34% ± 25% on average (P_Wilcox_ = 0.035). No profound shifts in microbial fermentation activity were observed for incubations with CMC and P80.

**Figure 7:**
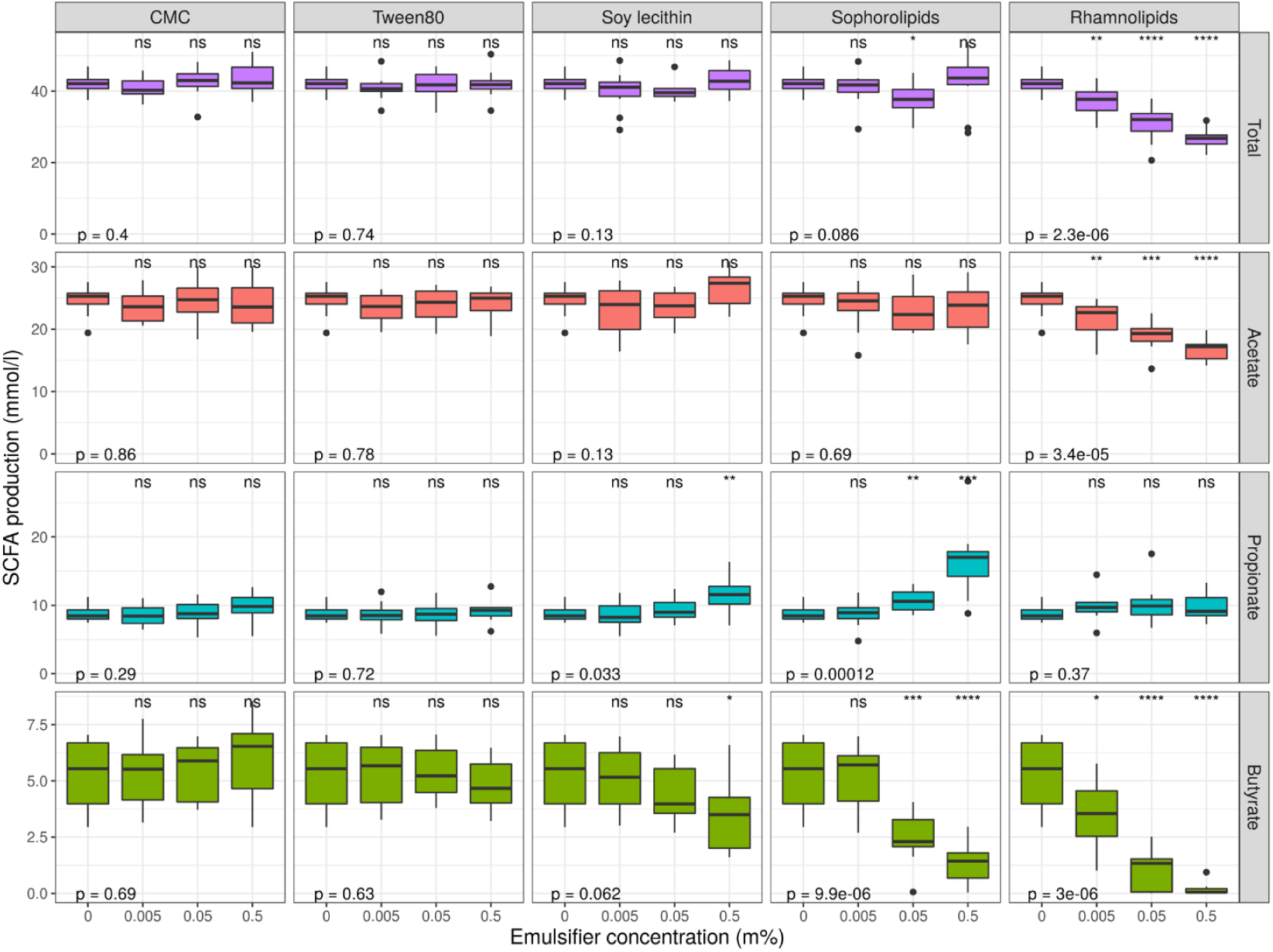
Short chain fatty acid production levels over 48h of incubation of fecal material from 10 donors in sugar depleted medium supplemented with 5 emulsifiers at 4 concentrations. Samples were taken upon incubation (T0; 2-3h after inoculation) as well as after 24h (T1) and 48h (T2) of incubation. Asterisks indicate significant differences with the control (0% (m/v)), calculated with Wilcoxon Rank Sum tests with Holm correction. P-values indicate results of general Kruskal-Wallis tests (α = 0.05).

#### 3.2.2 Metagenomic prediction

Other emulsifier related functional shifts were explored via metagenomic prediction using PICRUSt. These analyses predicted suppressing effects of rhamnolipids, sophorolipids on the pathways “Biosynthesis of other secondary metabolites”, “Cell growth and death” and “Signalling molecules and interaction” (Figure S6). Possible significantly upregulated L2-pathways were “Cell motility”, “Cellular Processes and signalling”, “Genetic information processing”, “Lipid metabolism”, “Membrane transport”, “Metabolism”, “Signal transduction”, “Transcription” and “Xenobiotics degradation”.

Plugging the data into the Bugbase-webtool revealed a significantly stimulating effect of sophorolipids and rhamnolipids on the formation of biofilms and mobile elements, stress tolerance and increased abundance of potential pathogens, gram negative and facultative anaerobic bacteria all properties related to the Proteobacteria phylum. This coincides with our observation of an increased abundance of *Escherichia/Shigella* (Supplementary Data 1).

#### 3.2.3 Flagellin levels

In order to validate the PICRUSt prediction of a higher motility potential a HEK-blue mTLR5 reporter cell assay was used for the detection of bacterial flagellin. The response of the flagellin concentrations to the emulsifiers was found to be largely donor-dependent and inconsistent shifts were observed in function of incubation time (Figure 8). Shifts in flagellin levels upon emulsifier dosage were variable, meaning that the prediction of higher motility by PICRUSt could not be substantiated.

**Figure 8:**
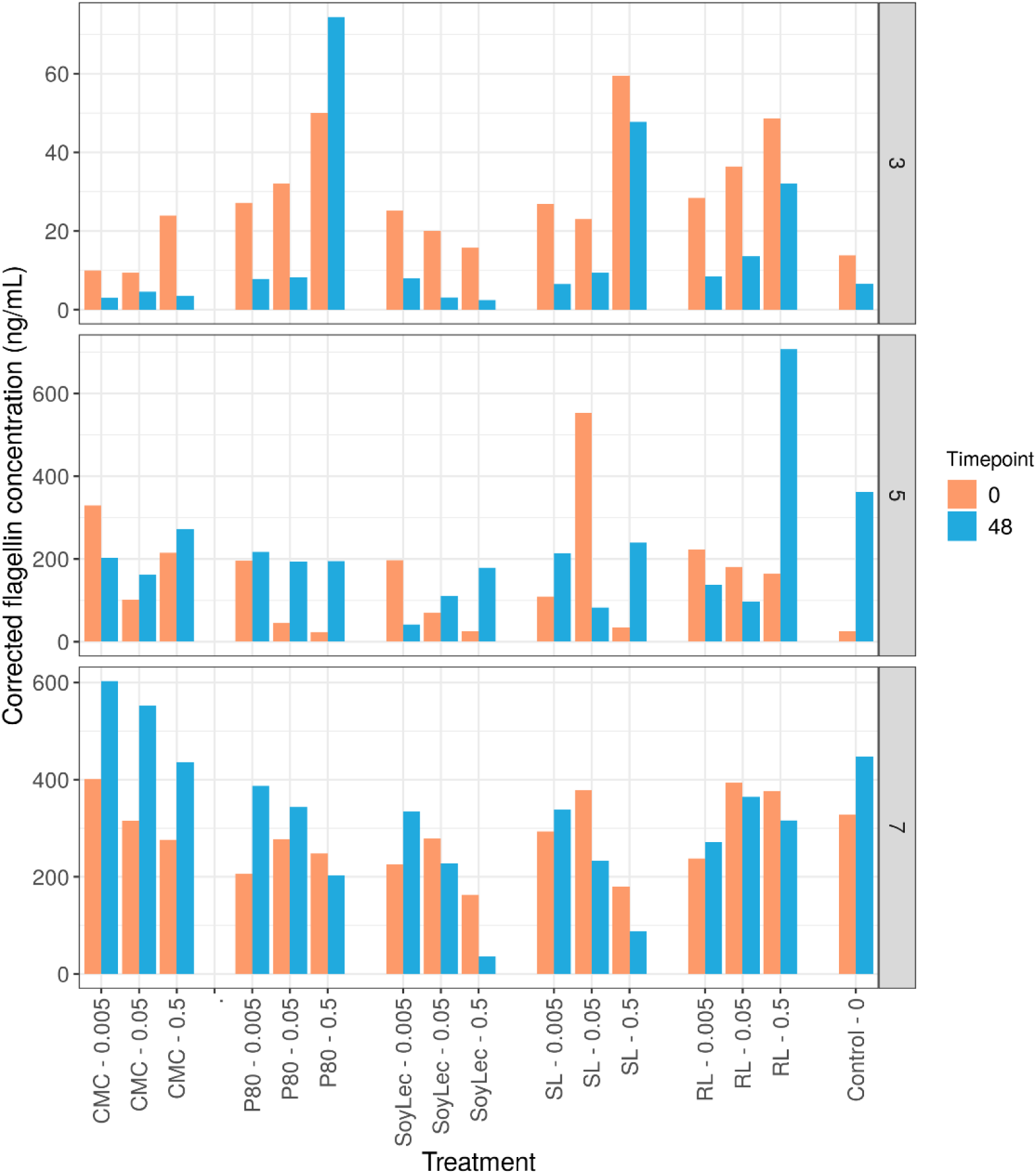
Flagellin concentrations obtained using mTLR5 HEK blue reporter cells for three donors at the start (T0) and the end (T2) of in vitro 48h batch experiments of fecal material from 10 donors in sugar depleted medium supplemented with 5 emulsifiers at 4 concentrations. Donors for the flagellin assay were selected based on their low (D3), high (D7) and intermediate (D5) metabolic response to the emulsifiers.

### 3.3 Donor Diversity

For all endpoints, inter-individual variability was observed in response to the *in vitro* incubations and emulsifier treatments. In terms of community structure, Figure 3A shows that each donor clusters separately. This clustering was found to be significant (P_dbRDA_ < 0.05).

To assess in more detail whether there was coherence in the read-outs with respect to donor susceptibility, we ranked the donors according to their response on the most relevant parameters, *i*.*e*. intact cell counts, production of the most important SCFA (acetate, propionate and butyrate) and the absolute and relative abundance of *Escherichia/Shigella*.

We observed that the susceptibility of the donors to the effects of the emulsifiers depended on the targeted parameter (Figure 9). Some donors consistently ranked as highly susceptible (D9) or less susceptible (D6 and D1), while for other donors, ranking was more variable over the different parameters.

**Figure 9:**
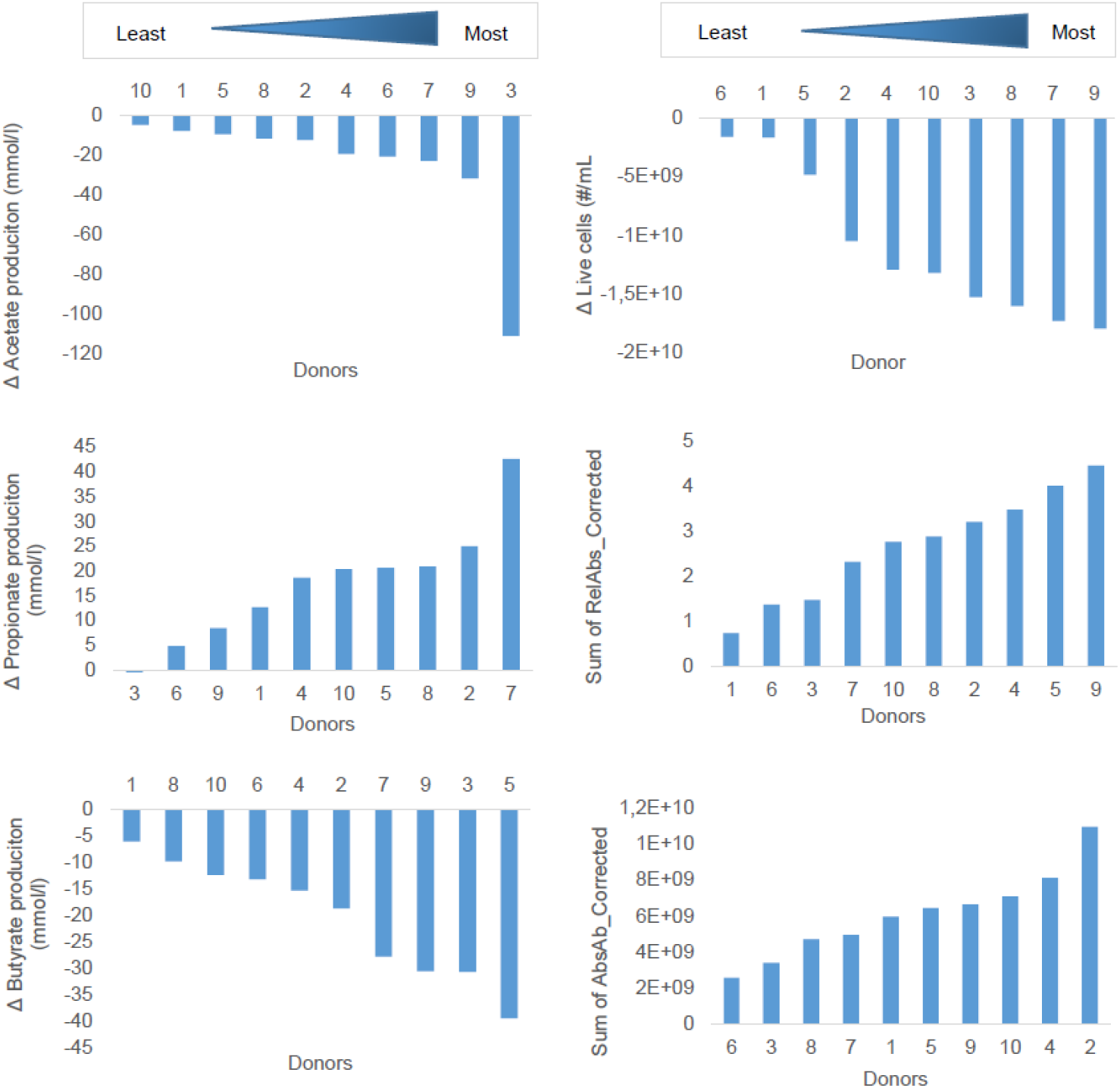
Ranking of donors of fecal material for 48h batch incubations in sugar depleted medium supplemented with 5 emulsifiers at 4 concentrations. Donors were ranked along the principal parameters impacted by the emulsifiers. Measures were calculated based on cumulative (sum of all treatments) change in 48h production of SCFA or cummulative 48h change in living cell counts, relative or absolute abundances of *Escherichia/Shigella*.

### 3.4 Equivalent emulsifier concentrations

We sought to compare the effects of biosurfactants *versus* those of conventional chemical emulsifiers. When comparing 0.5% (m/v) of chemical emulsifier with 0.05% (m/v) of biosurfactant – the concentrations most representative of the currently applied levels of dietary emulsifiers – the previously described effects of the biosurfactants (decreased levels of acetate, butyrate, intact and total cell concentration and increased abundance of propionate and increased abundance of *Escherichia/Shigella)* were significant compared to the effects of the chemical emulsifiers (Table S3). When comparing 0.05% (m/v) of chemical emulsifier with 0.005% (m/v) of biosurfactant, the effects were not significantly different, except for the effects of rhamnolipids on the cell count data.

## 4 Discussion

We found dietary emulsifiers to significantly alter human gut microbiota towards a composition and functionality with potentially higher pro-inflammatory properties. While donor-dependent differences in microbiota response were observed, our *in vitro* experimental setup showed these effects to be primarily emulsifier-dependent. Rhamnolipids and sophorolipids had the strongest impact with a sharp decrease in intact cell counts, an increased abundance in potentially pathogenic genera like *Escherichia/Shigella* and *Fusobacterium*, a decreased abundance of beneficial Bacteroidetes and *Barnesiella* and a predicted increase in flagellar assembly and general motility. The latter was not substantiated through direct measurements, though. The effects were less pronounced for soy lecithin, while chemical emulsifiers P80 and CMC showed the smallest effects. Short chain fatty acid production, with butyrate production in particular, was also affected by the respective emulsifiers, again in an emulsifier- and donor-dependent manner.

One of the most profound impacts of emulsifier treatment towards gut microbiota was the decline in intact microbial cell counts. The degree of microbiome elimination in this study seems comparable to what has been observed for antibiotic treatments (Guirro et al. 2019; Francino 2016). Since antibiotics are considered detrimental for gut ecology, this may serve as a warning sign with respect to emulsifier usage. Emulsifiers also act as surfactants, which are known for their membrane solubilizing properties (Jones 1999). The fact that the observed decline in microbial viability was dependent on emulsifier dose and on the emulsifying potential of the supplemented compound, as measured by the aqueous surface tension reduction (Table 1), leads us to conclude that the dietary emulsifiers attack the bacterial cells principally at the level of the cell membrane.

**Table 1:**
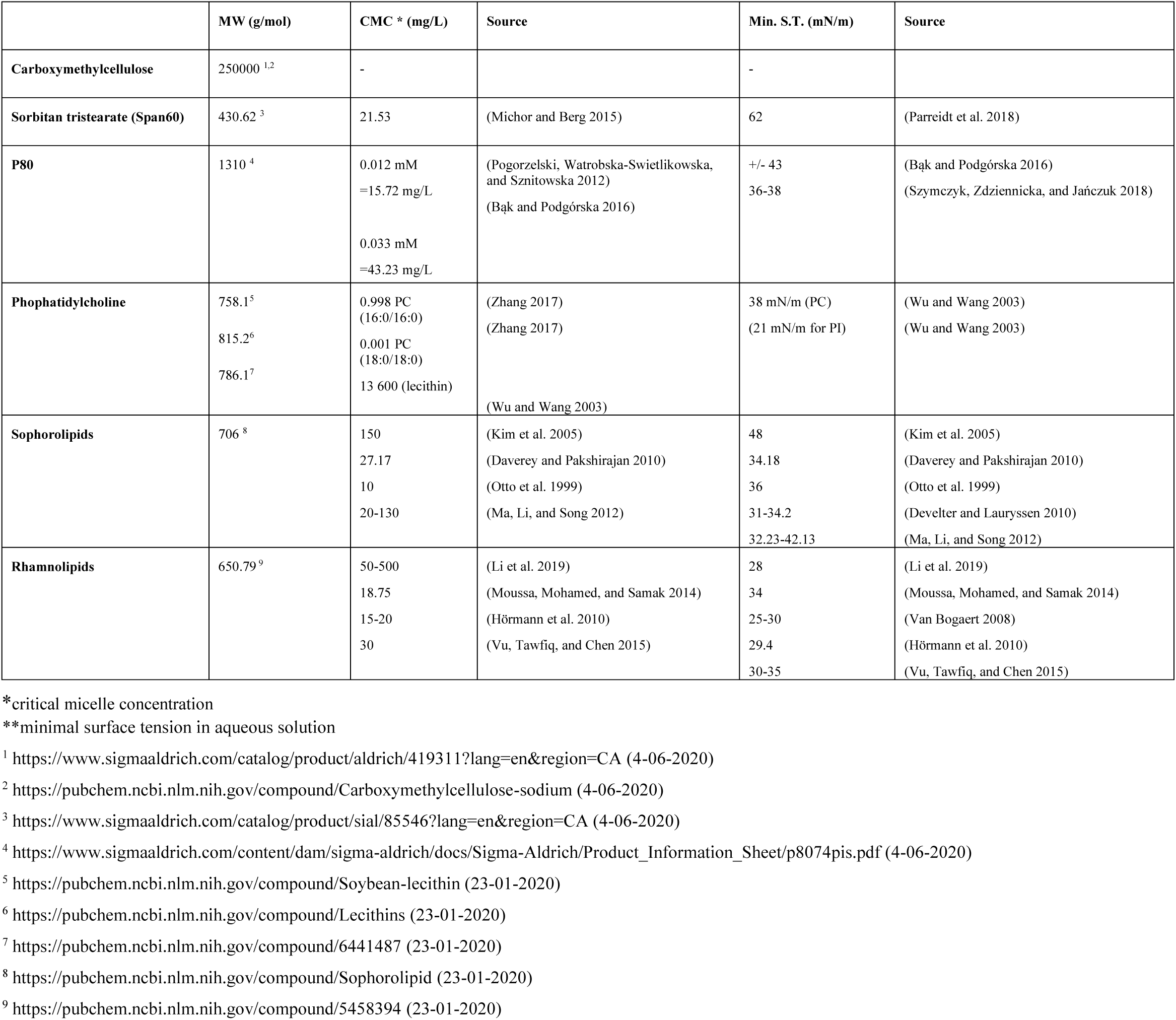
Emulsifier characteristics of CMC, Span60, P80, phosphatidylcholine (major component of soy lecithin), sophorolipids and rhamnolipids.

|  | MW (g/mol) | CMC * (mg/L) | Source | Min. S.T. (mN/m) | Source |
| --- | --- | --- | --- | --- | --- |
| **Carboxymethylcellulose** | 250000 <sup>1,2</sup> | - |  | - |  |
| **Sorbitan tristearate (Span60)** | 430.62 <sup>3</sup> | 21.53 | (Michor and Berg 2015) | 62 | (Parreidt et al. 2018) |
| **P80** | 1310 <sup>4</sup> | 0.012 mM<br>=15.72 mg/L<br>0.033 mM<br>=43.23 mg/L | (Pogorzelski, Watrobska-Swietlikowska, and Sznitowska 2012)<br>(Bąk and Podgórska 2016) | +/- 43<br>36-38 | (Bąk and Podgórska 2016)<br>(Szymczyk, Zdziennicka, and Jańczuk 2018) |
| **Phosphatidylcholine** | 758.1<sup>5</sup><br>815.2<sup>6</sup><br>786.1<sup>7</sup> | 0.998 PC (16:0/16:0)<br>0.001 PC (18:0/18:0)<br>13 600 (lecithin) | (Zhang 2017)<br>(Zhang 2017)<br>(Wu and Wang 2003) | 38 mN/m (PC)<br>(21 mN/m for PI) | (Wu and Wang 2003)<br>(Wu and Wang 2003) |
| **Sophorolipids** | 706 <sup>8</sup> | 150<br>27.17<br>10<br>20-130 | (Kim et al. 2005)<br>(Daverey and Pakshirajan 2010)<br>(Otto et al. 1999)<br>(Ma, Li, and Song 2012) | 48<br>34.18<br>36<br>31-34.2<br>32.23-42.13 | (Kim et al. 2005)<br>(Daverey and Pakshirajan 2010)<br>(Otto et al. 1999)<br>(Develter and Lauryssen 2010)<br>(Ma, Li, and Song 2012) |
| **Rhamnolipids** | 650.79 <sup>9</sup> | 50-500<br>18.75<br>15-20<br>30 | (Li et al. 2019)<br>(Moussa, Mohamed, and Samak 2014)<br>(Hörmann et al. 2010)<br>(Vu, Tawfiq, and Chen 2015) | 28<br>34<br>25-30<br>29.4<br>30-35 | (Li et al. 2019)<br>(Moussa, Mohamed, and Samak 2014)<br>(Van Bogaert 2008)<br>(Hörmann et al. 2010)<br>(Vu, Tawfiq, and Chen 2015) |

\*critical micelle concentration

We also found this antimicrobial effect from the tested emulsifiers to be highly selective, confirming previous observations by Moore (1997) who showed that the effects of surfactants are dependent on the bacterial species. We found the *Escherichia/Shigella* genus to be particularly resistant against the surfactants antimicrobial effects. This agrees with Kramer et al. (1984) and Nickerson and Aspedon (1992) who demonstrated the surfactant resistance of enteric bacteria *Enterobacter cloaca* and *E. coli* against sodium dodecyl sulphate. They showed that this resistance is widespread among *Enterobacteriaceae*, that it’s energy-dependent and that exposure to sodium dodecyl sulphate altered the expression of 19 proteins, of which three were later tentatively identified as clpP, clpB and clpX intracellular proteases (Rajagopal, Sudarsan, and Nickerson 2002). Also membrane-derived oligosaccharides, present in the periplasm of gram negative bacteria, were found to be essential to the detergent-resistant properties of *E. coli* (Rajagopal et al. 2003). Even though this could explain the increased abundance in this study for *Enterobacteriaceae* and *Escherichia/Shigella*, the mechanism by which other species are (un)affected by the emulsifiers is not known.

Interestingly, our prediction of altered functionality using PICRUSt analysis indicated a potential increase in levels of motility-genes, even though we could not confirm increased flagellin levels by direct measurement. Increased motility has been observed upon addition of emulsifiers before (Lock et al. 2018). This finding may again be related to the increased relative abundance of *Escherichia/Shigella*, since these are known flagella (gene)-bearers (Mittal et al. 2003; Tominaga et al. 2001; Girón 1995). Increased flagellin levels constitute a potential health risk as flagellin is considered an important virulence factor: this particular microbe-associated molecular pattern may trigger inflammation upon binding to TLR5. A higher degree of flagellation would thus represent more motile bacteria and result in a gut microbiome that is able to more aggressively penetrate the mucus layer and subsequently reach gut epithelial cells (Chassaing et al. 2015; Ramos, Rumbo, and Sirard 2004). Chassaing et al. (2015; 2017) demonstrated possible consequences for the host: an increased inflammatory response in both gut and body, contributing to increased adiposity and weight gain.

Effects towards microbial metabolic functionality, as measured by SCFA-levels, were emulsifier-dependent. Nevertheless, consistent shifts in SCFA-profiles were observed for all three emulsifiers for which significant alterations were visible, *i*.*e*. a decrease in butyrate and an increase in propionate production. For sophorolipids, the strong increase in propionate production can be related to the increased abundance of *Phascolarctobacterium*. This genus is known to produce propionate from succinate produced by *E. coli* cells (Del Dot, Osawa, and Stackebrandt 1993). For rhamnolipids, we assume that similar cross-feeding interactions occurred, even though propionate levels remained stable. We hypothesize that an increased propionate production by the *Escherichia/Phascolarctobacterium* consortium is counteracted by the higher antimicrobial activities from the rhamnolipids at increasing concentrations. The fact that L-rhamnose is a well-known propionate precursor (Reichardt et al. 2014) and that rhamnolytic pathways have been observed in *E. coli* and other Gammaproteobacteria (Rodionova et al. 2013) further support this idea. The increase in acetate and propionate production observed with soy lecithin can be linked to the metabolism of glycerol (De Weirdt et al. 2010). This is in agreement with the upregulated abundances of the *Enterococcus* and *Clostridia* genera at higher levels of soy lecithin, since these genera are known to metabolize glycerol (Bradbeer 1965; Bizzini et al. 2010) as well as choline (Martínez-del Campo et al. 2015). Also the increased abundance of the *Acidaminoccus* genus would correspond with the increased acetate production (Chang et al. 2010). These findings indicate that observed shifts in SCFA-production can be attributed to shifts in microbial composition.

Whether the altered SCFA-production and -levels are positive or negative for host health is ambiguous. On the one hand, decreased butyrate levels can be considered negative, since butyrate is known to protect the gut epithelium from inflammation and cancerous growth (Liu et al. 2018; Cani 2017; Canani et al. 2011). On the other hand, propionate is also considered a health-promoting SCFA (Hosseini et al. 2011; Weitkunat et al. 2016; Louis and Flint 2017). Propionate production is related to decreased lipogenesis in the liver and is supposed to enhance satiety mechanisms, which would lower the chance of developing obesity (Hosseini et al. 2011; Weitkunat et al. 2016). It is, however, questionable whether such benefit may predominate the purported negative effects from the observed antimicrobial effects and the increase in pro-inflammatory markers, such as flagellated microbiota and a drop in butyrate production.

Another point relating to health effects entails that rhamnolipids, sophorolipids and – to a lesser extent - soy lecithin significantly decreased diversity parameters. Decreased microbiome diversity frequently occurs with compromised health conditions such as obesity, insulin resistance, dyslipideamia (Turnbaugh et al. 2009; Le Chatelier et al. 2013), type 1 diabetes (Patterson et al. 2016), heart failure (Luedde et al. 2017) and inflammatory bowel disease (Arumugam et al. 2011; Benoit Chassaing et al. 2015). The increased prevalence of the *Escherichia/Shigella* genus has also been observed with multiple metabolic conditions (Shin, Whon, and Bae 2015; Lecomte et al. 2015; Lupp et al. 2007). Increased *Enterobacteriaceae* abundance was previously linked with an increased gut permeability (Pedersen et al. 2018), the consumption of high fat diets (Fei and Zhao 2013; He et al. 2018; Lecomte et al. 2015), colitis (Lupp et al. 2007), cardiovascular disease (Jie et al. 2017), diabetes (Deschasaux et al. 2019; Allin, Nielsen, and Pedersen 2015) and even undernourishment and iron deficiency anemia (Shin, Whon, and Bae 2015; Muleviciene et al. 2018). With respect to soy lecithin, metabolism of phosphatidylcholine by the gut microbiota has been linked to cardiovascular disease (Wang et al. 2011; Tang and Hazen 2014). We also found an increased abundance of the genus *Sutterella*, a bacterium that has been linked to autism spectrum disorders (Wang et al. 2013) and gut inflammation, notably by IgA-degradation (Kaakoush 2020; Moon et al. 2015). Overall, we can thus conclude that dietary emulsifier consumption may result in profound microbiota shifts, putatively contributing to adverse health outcomes.

An important consideration is whether the observed *in vitro* effects would also take place in an *in vivo* setting. This will depend on a number of diet- and host-related factors. First, we found that the observed effects were concentration- and emulsifier-dependent. Choosing an emulsifier-concentration combination that minimizes adverse microbial impacts, without compromising food technical properties, could thus be a strategy to mitigate the harmful effects of food emulsifiers. Second, emulsifier concentration will continuously alter during gastrointestinal digestion, but the impact of the dilution with accompanying food products, excretion of digestive fluids or absorption of water from the gut lumen on the final concentration reaching the gut microbiota has so far not yet been studied. Third, digestion by human enzymes will alter chemical structure of the emulsifiers. While CMC is resistant to breakdown by human digestive enzymes (Joint FAO/WHO Expert Committee on Food Additives 1973), P80 and soy lecithin can be hydrolyzed by pancreatic lipases. For P80, only the polyethylene-sorbitan unit may reach the colon (Aguilar et al. 2015). Soy lecithin is mostly absorbed as lysolecithin and free fatty acids, but its detection in faeces indicates that some fraction reaches the colon (Mortensen et al. 2017), where the choline and glycerol moieties are metabolized by the gut microbiota (Tang and Hazen 2014). For both of these compounds, it will thus be necessary to verify if and to what extent the observed effects occur *in vivo* as well. With respect to rhamnolipids and sophorolipids, no literature is currently available on their digestion by human enzymes. Preliminary data from our side, however, have indicated only one alteration, a deacetylation of the sophorose units of the sophorolipids (supplementary data 2). These compounds may thus readily reach the colon and interact with the endogenous microbiota. Finally, other dietary constituents in the gut, primarily lipids and bile salts, will interact with emulsifiers (Naso et al. 2019). The mucus layer overlying the gut epithelium and the pH-fluctuations throughout the gut are other elements that may affect emulsifier- microbiota interactions.

A last important element in the putative health impact from dietary emulsifiers concerns interindividual variability. An individual’s unique microbiota and metabolism are important determinants of the potential health effects dietary emulsifiers could cause. While the overall effects from the different emulsifiers towards microbiota composition and functionality were quite consistent in our study, important interindividual differences in susceptibility of the microbiota were noted. Understanding what underlying factors and determinants drive this interindividual variability will be crucial to future health risk assessment of novel and existing dietary emulsifiers.

Food additives have come under scrutiny with respect to their impact on human health. Additives like colorants, artificial sweeteners, nitrites (NaNO_2_) and high fructose corn syrup have been associated with hyperactivity, cancer development, gastric cancer and obesity, respectively. (Arnold, Lofthouse, and Hurt 2012; Carocho et al. 2014; Bryan et al. 2012; Payne, Chassard, and Lacroix 2012). In answer to these public concerns, a clean label movement has developed in the food industry that aims to provide food products with a more natural image. In this light, we investigated whether rhamnolipids and sophorolipids, two biotechnological emulsifiers, would yield less of a detrimental impact on the gut microbiota than the mainstream chemical emulsifiers, CMC and P80. Our results showed however, that rhamnolipids and sophorolipids, of all emulsifiers in this study, had the strongest impact on microbiota composition and functionality, even when equivalent concentrations were taken into account. Further analysis of our data showed that the observed detrimental effects towards the microbiota can potentially be linked to their higher emulsifying potential. All of this indicates that rhamnolipids and sophorolipids are probably no appropriate alternatives to conventional emulsifiers unless they are used at substantially lower concentrations. More research must point out whether the effects prevail *in vivo* and whether concentrations can be kept low enough to avoid adverse health effects.

## Supporting information

Supplementary Data 1 - Bugbase

Supplementary Data 2 - RL&SL Digestion Chromatography

## 9 Author Contributions

This research was performed under promotorship of Prof. Tom Van de Wiele, Prof. John Van Camp and Prof. Andreja Raikovic. Lisa Miclotte was responsible for execution of all experiments, data visualisation and statistical analysis, as well as the assembly and submission of the manuscript. Chris Callewaert assisted in the execution of the PICRUSt algorythm. DESeq analysis was performed in collaboration with Kim De Paepe. Library preparation and 16S rRNA gene amplicon sequencing was performed by Leen Rymenans, collaborator withing Jeroen Raes’ group. All co-author have reviewed the manuscript before submission.

## 11 Acknowledgments

We thank prof. Inge Van Bogaert (Inbio, Ghent University) for providing us with sophorolipids samples. We also thank Jo Devrieze for providing feedback to our manuscript before submission.

## 12 Data Availability Statement

The 16S rRNA Illumina MiSeq sequencing data generated for this study can be found in the NCBI- database under accession number PJNR PRJNA63057

## Footnotes

1 European Commision. 2014. “00191 - Sodium Carboxymethyl Cellulose - E466.” 2014. https://webgate.ec.europa.eu/foods_system/main/index.cfm?event=substance.view&identifier=191.

2 FDA. 2019. “CFR - Code of Federal Regulations Title 21 - Sodium Carboxymethyl Cellulose.” 2019. https://www.accessdata.fda.gov/scripts/cdrh/cfdocs/cfCFR/CFRSearch.cfm?fr=182.1745.

3 FDA. 2020. “CARBOXYMETHYL CELLULOSE, SODIUM SALT.” 2020. https://www.accessdata.fda.gov/scripts/fdcc/?set=FoodSubstances&id=CARBOXYMETHYLCELLULOSESODIUMSALT&sort=Sortterm&order=ASC&startrow=1&type=basic&search=carboxymethyl.

4 European Commision. “00172 - Polysorbates -E433.” 2019. Accessed August 6, 2019. https://webgate.ec.europa.eu/foods_system/main/index.cfm?event=substance.view&identifier=172.

5 FDA. “CFR - Code of Federal Regulations Title 21 - Polysorbate 80.” 2019. Accessed August 6, 2019. https://www.accessdata.fda.gov/scripts/cdrh/cfdocs/cfCFR/CFRSearch.cfm?fr=172.840.

6 European Commision 2018. “00115 -Soy Lecithin - E322.” 2018. https://webgate.ec.europa.eu/foods_system/main/index.cfm?event=substance.view&identifier=115.

7 FDA. 2019. “CFR - Code of Federal Regulations Title 21 - Soy Lecithin.” 2019. https://www.accessdata.fda.gov/scripts/cdrh/cfdocs/cfCFR/CFRSearch.cfm?fr=184.1400.

