## Supplementary Figures and Tables for "Dietary emulsifiers alter composition and activity of the human gut microbiota *in vitro*, irrespective of chemical or natural emulsifier origin"

*Supplementary Material*

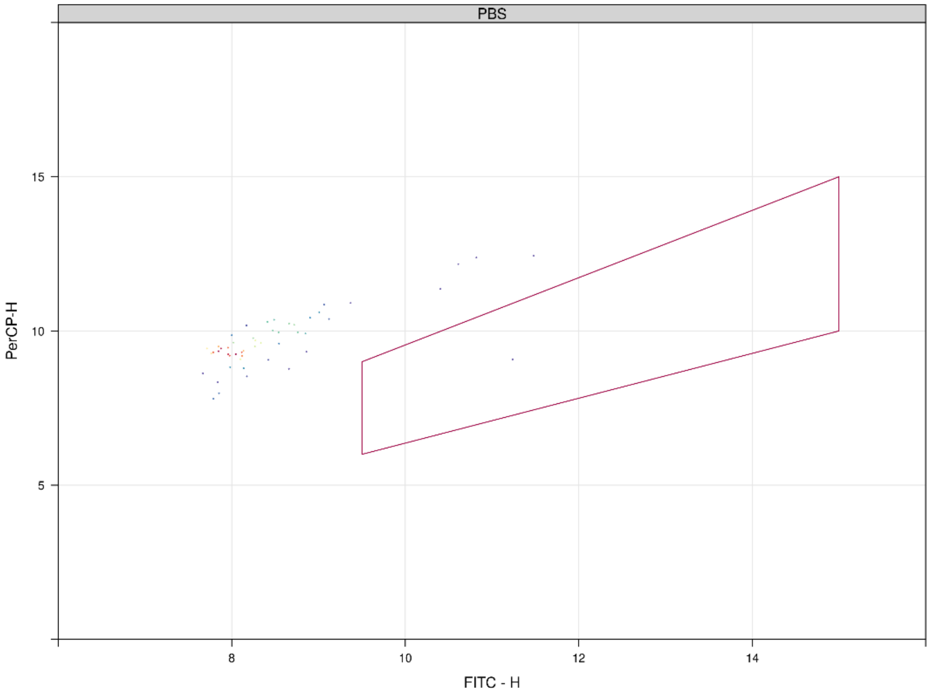


**Figure S1: Density plot of cell counts for the PBS control sample measured in conjuction with samples from for in vitro batch incubations of fecal material from 10 donors with sugar depleted medium supplemented with 5 emulsifiers at 4 concentrations. The gate represents intact cell counts. The lack of counts in this gate indicates limited background from the PBS-matrix.**

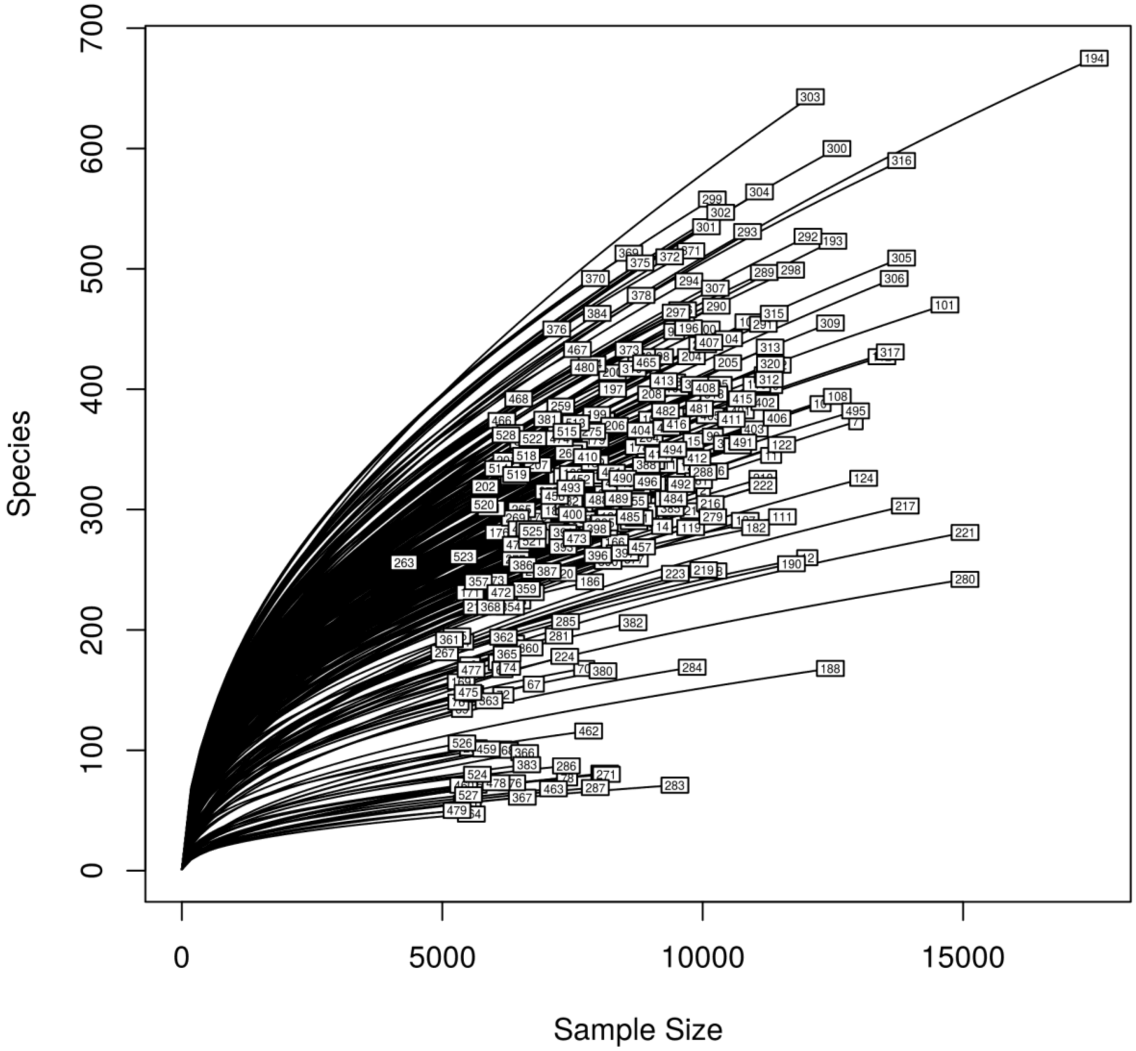


**Figure S2: Rarefaction curve of amplicon sequencing data after copynumber correction on the data from in vitro batch incubations of fecal material from 10 donors with sugar depleted medium supplemented with 5 emulsifiers at 4 concentrations.The numbers on the curves represent sample names which can be looked up in the NCBI submission with the accession code the accession code PRJNA63057.**

**Table S1: Intact cell counts (logarithmic) and percentage of surviving cells on the three measured timepoints for in vitro batch incubations of fecal material from 10 donors with sugar depleted medium supplemented with 5 emulsifiers at 4 concentrations. Samples were taken upon incubation (T0; 2-3h after inoculation) as well as after 24h (T1) and 48h (T2) of incubation. Concentrations of emulsifiers are given in % (m/v).**

| | 0h | | 24h | | 48h | | | |
|---|---|---|---|---|---|---|---|---|
| | Mean of Intact cells | Cell count / Cell count control-T0 | Mean of Intact cells | Cell count T1 / Cell count control-T0 | Mean of Intact cells | Cell count T2 / Cell count T0 | Cell count T2 / Cell count control-T0 | Cell count T2 / Cell count control-T2 |
| **CMC** | | | | | | | | |
| 0 | 9.48 ± 0.23 | 100% ± 3% | 9.52 ± 0.23 | 108% ± 4% | 9.38 ± 0.20 | 79% ± 3% | 79% ± 3% | 100% ± 3% |
| 0.005 | 9.56 ± 0.23 | 119% ± 4% | 9.55 ± 0.21 | 98% ± 3% | 9.40 ± 0.26 | 69% ± 3% | 82% ± 3% | 104% ± 4% |
| 0.05 | 9.59 ± 0.23 | 128% ± 4% | 9.59 ± 0.22 | 99% ± 3% | 9.44 ± 0.23 | 71% ± 2% | 91% ± 3% | 115% ± 4% |
| 0.5 | 9.56 ± 0.25 | 119% ± 4% | 9.60 ± 0.21 | 111% ± 4% | 9.49 ± 0.22 | 85% ± 3% | **101% ± 3%** | 128% ± 4% |
| **P80** | | | | | | | | |
| 0 | 9.48 ± 0.23 | 100% ± 3% | 9.52 ± 0.23 | 108% ± 4% | 9.38 ± 0.20 | 79% ± 3% | 79% ± 3% | 100% ± 3% |
| 0.005 | 9.54 ± 0.21 | 114% ± 4% | 9.51 ± 0.19 | 93% ± 3% | 9.39 ± 0.22 | 71% ± 2% | 81% ± 3% | 102% ± 3% |
| 0.05 | 9.55 ± 0.20 | 117% ± 4% | 9.50 ± 0.24 | 89% ± 3% | 9.36 ± 0.21 | 64% ± 2% | 75% ± 2% | 95% ± 3% |
| 0.5 | 9.57 ± 0.19 | 121% ± 4% | 9.51 ± 0.14 | 88% ± 2% | 9.42 ± 0.15 | 71% ± 2% | **86% ± 2%** | 109% ± 3% |
| **SoyLec** | | | | | | | | |
| 0 | 9.48 ± 0.23 | 100% ± 3% | 9.52 ± 0.23 | 108% ± 4% | 9.38 ± 0.20 | 79% ± 3% | 79% ± 3% | 100% ± 3% |
| 0.005 | 9.47 ± 0.27 | 96% ± 4% | 9.41 ± 0.21 | 88% ± 3% | 9.27 ± 0.19 | 64% ± 2% | 61% ± 2% | 78% ± 2% |
| 0.05 | 9.40 ± 0.34 | 82% ± 4% | 9.21 ± 0.21 | 65% ± 3% | 9.04 ± 0.25 | 44% ± 2% | 36% ± 1% | 46% ± 2% |
| 0.5 | 9.44 ± 0.29 | 92% ± 4% | 9.08 ± 0.11 | 43% ± 1% | 8.85 ± 0.13 | 25% ± 1% | **23% ± 1%** | 30% ± 1% |
| **SL** | | | | | | | | |
| 0 | 9.48 ± 0.23 | 100% ± 3% | 9.52 ± 0.23 | 108% ± 4% | 9.38 ± 0.20 | 79% ± 3% | 79% ± 3% | 100% ± 3% |
| 0.005 | 9.57 ± 0.18 | 122% ± 4% | 9.52 ± 0.21 | 89% ± 3% | 9.37 ± 0.23 | 64% ± 2% | 78% ± 3% | 98% ± 3% |
| 0.05 | 9.19 ± 0.37 | 51% ± 2% | 9.23 ± 0.15 | 110% ± 5% | 9.07 ± 0.16 | 76% ± 3% | 39% ± 1% | 49% ± 1% |
| 0.5 | 8.80 ± 0.46 | 21% ± 1% | 8.88 ± 0.17 | 120% ± 7% | 8.79 ± 0.19 | 99% ± 6% | **21% ± 1%** | 26% ± 1% |
| **RL** | | | | | | | | |
| 0 | 9.48 ± 0.23 | 100% ± 3% | 9.52 ± 0.23 | 108% ± 4% | 9.38 ± 0.20 | 79% ± 3% | 79% ± 3% | 100% ± 3% |
| 0.005 | 9.42 ± 0.33 | 86% ± 4% | 9.32 ± 0.21 | 79% ± 3% | 9.12 ± 0.20 | 50% ± 2% | 43% ± 1% | 55% ± 2% |
| 0.05 | 9.00 ± 0.37 | 33% ± 2% | 9.00 ± 0.12 | 100% ± 5% | 8.73 ± 0.13 | 53% ± 2% | 18% ± 0% | 22% ± 1% |
| 0.5 | 8.59 ± 0.43 | 13% ± 1% | 8.80 ± 0.22 | 162% ± 9% | 8.45 ± 0.20 | 72% ± 4% | **9% ± 0%** | 12% ± 0% |

**Table S2: Total cell counts (logarithmic) and percentage of remaining cells on the three measured timepoints for in vitro batch incubations of fecal material from 10 donors with sugar depleted medium supplemented with 5 emulsifiers at 4 different concentrations. Samples were taken upon incubation (T0; 2-3h after inoculation) as well as after 24h (T1) and 48h (T2) of incubation. Concentrations of emulsifiers are given in % (m/v).**

| | 0h | | 24h | | 48h | | | |
|---|---|---|---|---|---|---|---|---|
| | Mean of Total Cells | Cell count / Cell count control-T0 | Mean of Intact cells | Cell count T1 / Cell count control-T0 | Mean of Intact cells | Cell count T2 / Cell count T0 | Cell count T2 / Cell count control-T0 | Cell count T2 / Cell count control-T2 |
| **CMC** | | | | | | | | |
| 0 | 9.66 ± 0.16 | 100% + 2% | 9.70 ± 0.10 | 110% + 2% | 9.59 ± 0.10 | 86% + 2% | 86% + 2% | 100% + 1% |
| 0.005 | 9.71 ± 0.18 | 112% + 3% | 9.71 ± 0.12 | 101% + 2% | 9.62 ± 0.15 | 81% + 2% | 91% + 2% | 105% + 2% |
| 0.05 | 9.73 ± 0.16 | 118% + 3% | 9.75 ± 0.11 | 103% + 2% | 9.65 ± 0.12 | 83% + 2% | 98% + 2% | 114% + 2% |
| 0.5 | 9.71 ± 0.19 | 113% + 3% | 9.77 ± 0.10 | 115% + 3% | 9.71 ± 0.11 | 99% + 2% | 112% + 2% | 130% + 2% |
| **P80** | | | | | | | | |
| 0 | 9.66 ± 0.16 | 100% + 2% | 9.70 ± 0.10 | 110% + 2% | 9.59 ± 0.10 | 86% + 2% | 86% + 2% | 100% + 1% |
| 0.005 | 9.71 ± 0.14 | 112% + 3% | 9.69 ± 0.09 | 96% + 2% | 9.62 ± 0.10 | 82% + 1% | 92% + 2% | 107% + 2% |
| 0.05 | 9.71 ± 0.16 | 112% + 3% | 9.70 ± 0.15 | 98% + 2% | 9.58 ± 0.14 | 74% + 2% | 83% + 2% | 96% + 2% |
| 0,5 | 9.71 ± 0.17 | 113% + 3% | 9.70 ± 0.09 | 96% + 3% | 9.62 ± 0.09 | 80% + 2% | 90% + 2% | 105% + 1% |
| **SoyLec** | | | | | | | | |
| 0 | 9.66 ± 0.16 | 100% + 2% | 9.70 ± 0.10 | 110% + 2% | 9.59 ± 0.10 | 86% + 2% | 86% + 2% | 100% + 1% |
| 0.005 | 9.69 ± 0.17 | 106% + 3% | 9.67 ± 0.12 | 96% + 2% | 9.57 ± 0.11 | 76% + 2% | 81% + 2% | 94% + 1% |
| 0.05 | 9.68 ± 0.17 | 106% + 3% | 9.60 ± 0.12 | 82% + 2% | 9.48 ± 0.18 | 62% + 2% | 66% + 2% | 76% + 2% |
| 0.5 | 9.73 ± 0.17 | 118% + 3% | 9.63 ± 0.08 | 78% + 2% | 9.55 ± 0.12 | 65% + 1% | 77% + 2% | 89% + 1% |
| **SL** | | | | | | | | |
| 0 | 9.66 ± 0.16 | 100% + 2% | 9.70 ± 0.10 | 110% + 2% | 9.59 ± 0.10 | 86% + 2% | 86% + 2% | 100% + 1% |
| 0.005 | 9.72 ± 0.14 | 115% + 3% | 9.71 ± 0.12 | 97% + 2% | 9.60 ± 0.13 | 76% + 1% | 87% + 2% | 101% + 2% |
| 0.05 | 9.68 ± 0.16 | 105% + 2% | 9.59 ± 0.13 | 82% + 2% | 9.49 ± 0.15 | 64% + 1% | 67% + 2% | 78% + 1% |
| 0.5 | 9.60 ± 0.19 | 87% + 2% | 9.37 ± 0.16 | 59% + 2% | 9.26 ± 0.20 | 45% + 1% | 40% + 1% | 46% + 1% |
| **RL** | | | | | | | | |
| 0 | 9.66 ± 0.16 | 100% + 2% | 9.70 ± 0.10 | 110% + 2% | 9.59 ± 0.10 | 86% + 2% | 86% + 2% | 100% + 1% |
| 0.005 | 9.70 ± 0.16 | 110% + 2% | 9.64 ± 0.13 | 87% + 2% | 9.51 ± 0.13 | 64% + 1% | 71% + 2% | 82% + 1% |
| 0.05 | 9.65 ± 0.19 | 98% + 3% | 9.47 ± 0.14 | 67% + 2% | 9.32 ± 0.15 | 47% + 1% | 46% + 1% | 54% + 1% |
| 0.5 | 9.56 ± 0.17 | 80% + 2% | 9.33 ± 0.13 | 58% + 1% | 9.22 ± 0.20 | 45% + 1% | 36% + 1% | 42% + 1% |

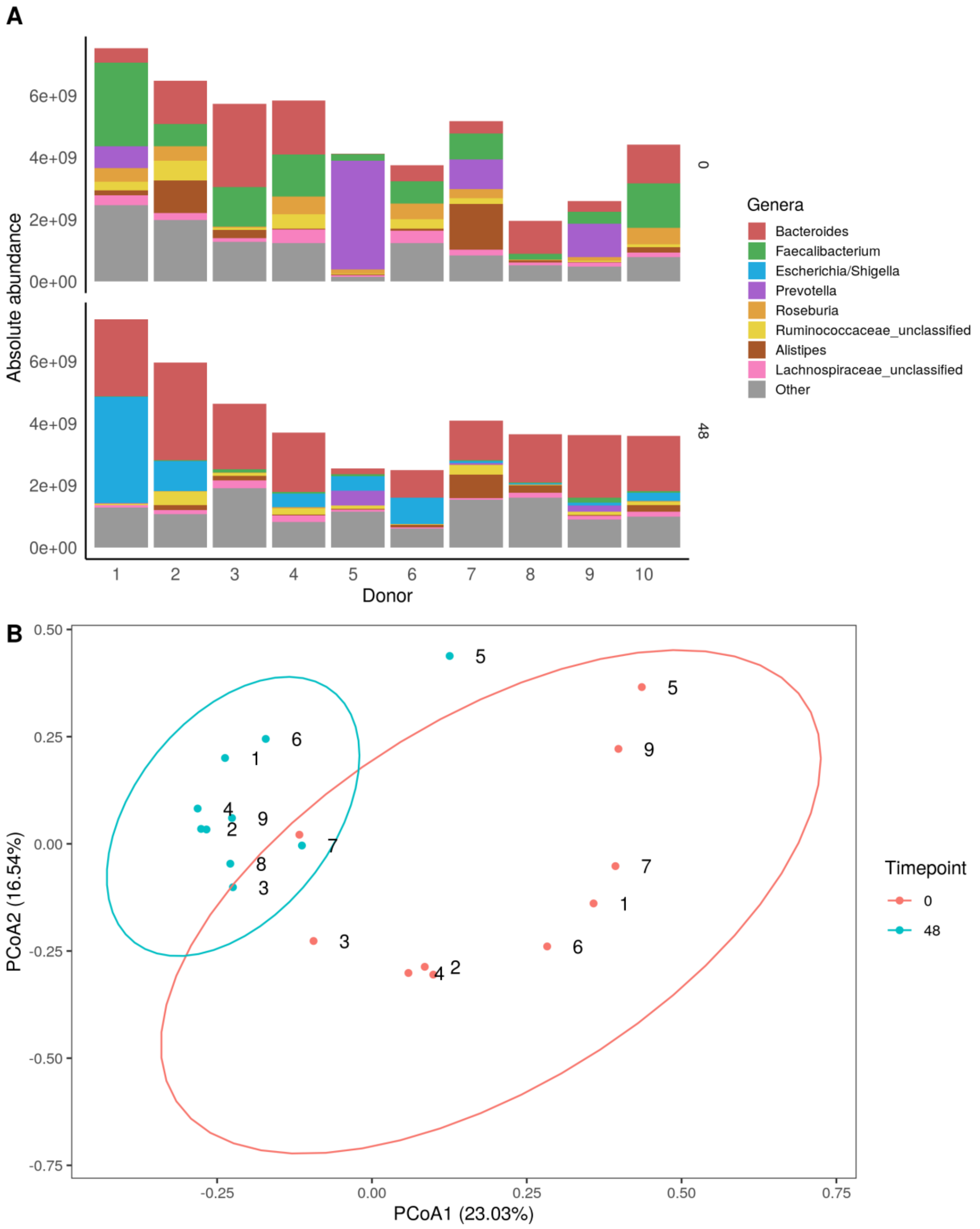


**Figure S3: A: Quantitative microbial profiling for control samples from 10 donors at the start (upper barplot) and at the end (lower barplot) of the in vitro batch incubations of fecal material with sugar depleted medium supplemented with 5 emulsifiers at 4 concentrations. Samples were taken upon incubation (T0; 2-3h after inoculation) as well as after 24h (T1) and 48h (T2) of incubation. B: Principle coordinate analysis of genus based QMP-data of control samples. Labels indicate donors.**

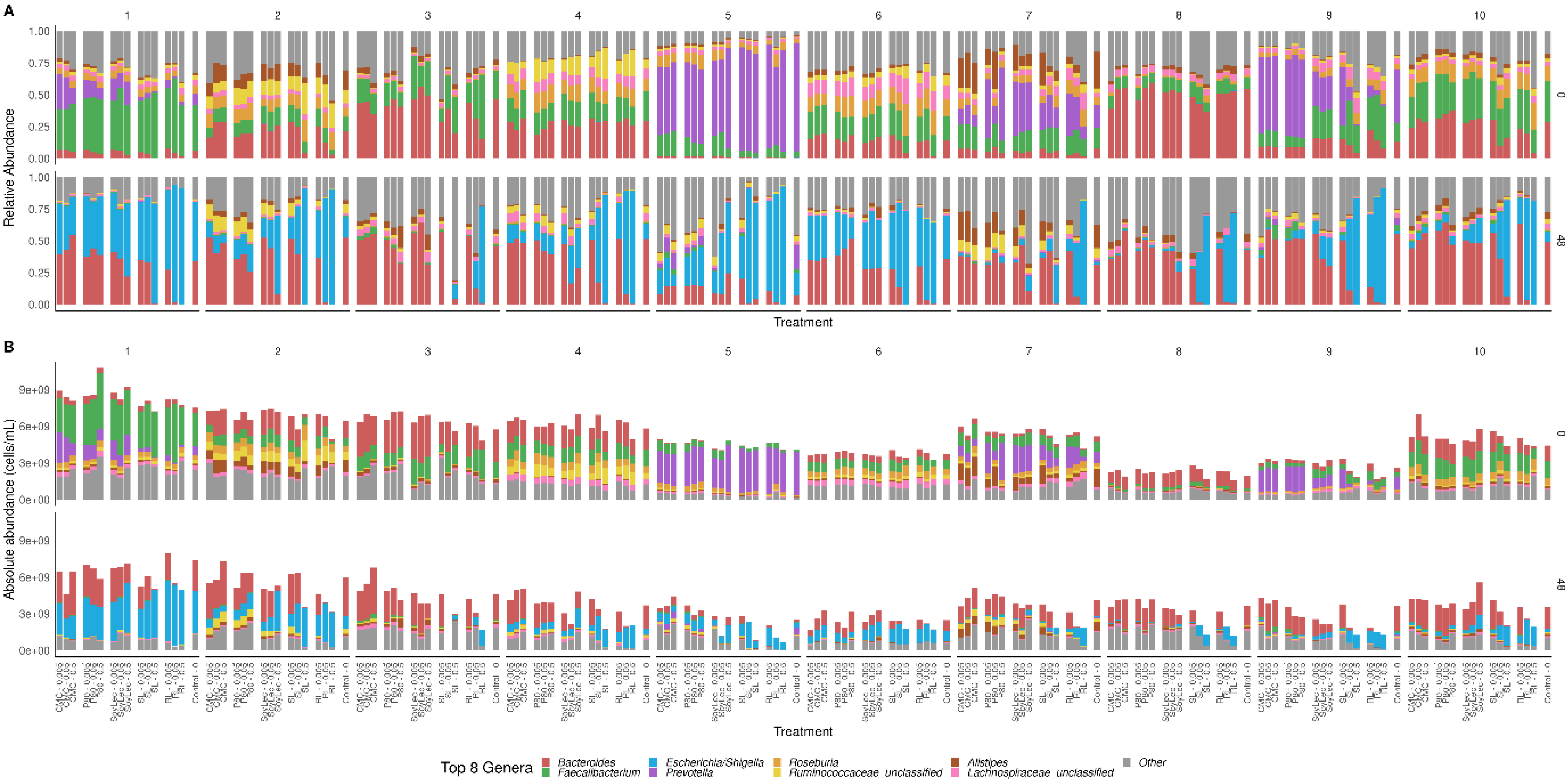


**Figure S4: Relative abundances (A) and absolute abundances (B) of top 8 genera detected at the start (0h) and at the end (48h) of the in vitro batch incubations of fecal material from 10 donors with sugar depleted medium supplemented with 5 emulsifiers at 4 concentrations. Samples were taken upon incubation (T0; 2-3h after inoculation) as well as after 24h (T1) and 48h (T2) of incubation.**

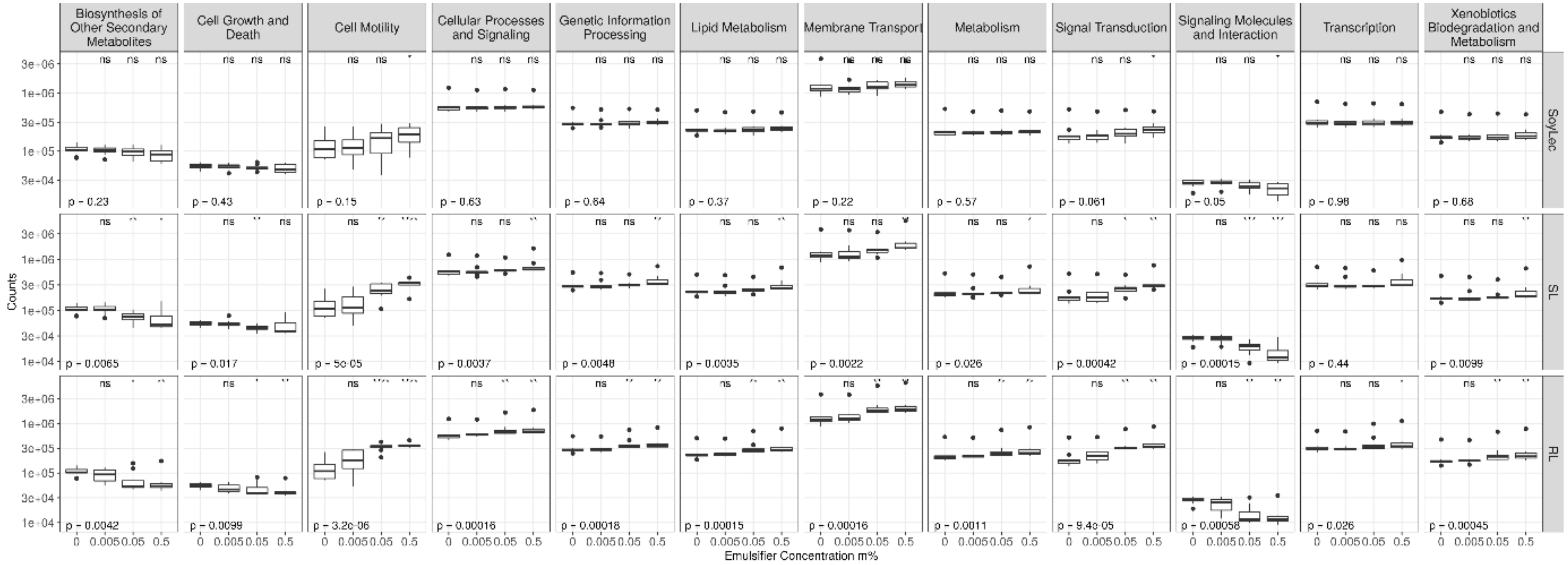


**Figure S5: Phenotypic prediction of gut microbial communities after 48h of in vitro batch incubations of fecal material from 10 donors with sugar depleted medium supplemented with 5 emulsifiers at 4 concentrations. Prediction of microbial functionalities was made using PICRUSt, based on the Kyoto Encyclopedia of Genes and Genomes (KEGG) database. Significantly different metagenomic pathways at KEGG L2-level are given for soy lecithin, sophorolipids and rhamnolipids. P-values represent results of Kruskal-Wallis test, asterisks represent results from Wilcoxon Rank Sum test of comparisons with control (α = 0.05).**

**Table S3: P-values resulting from Wilcoxon Rank Sum tests (α = 0.05) comparing the effect of equivalent emulsifier concentrations on short chain fatty acid concentrations, cell counts and the principal OTU detected using amplicon sequencing detected for in vitro batch incubations of fecal material from 10 donors with sugar depleted medium supplemented with 5 emulsifiers at 4 concentrations.**

**Acetate**

| Vs. | RL - 0.005 | SL - 0.005 | RL - 0.05 | SL - 0.05 |
|---|---|---|---|---|
| **CMC - 0.05** | 0,079 | 0,720 | | |
| **CMC - 0.5** | | | **0,001** | 0,393 |
| **P80 - 0.05** | 0,105 | 0,971 | | |
| **P80 - 0.5** | | | **0,002** | 0,481 |

**Total cell concentration**

| Vs. | RL - 0.005 | SL - 0.005 | RL - 0.05 | SL - 0.05 |
|---|---|---|---|---|
| **CMC - 0.05** | **0,000** | 0,201 | | |
| **CMC - 0.5** | | | **0,000** | **0,000** |
| **P80 - 0.05** | 0,221 | 0,718 | | |
| **P80 - 0.5** | | | **0,005** | 0,678 |

**Propionate**

| Vs. | RL - 0.005 | SL - 0.005 | RL - 0.05 | SL - 0.05 |
|---|---|---|---|---|
| **CMC - 0.05** | 0,211 | 0,968 | | |
| **CMC - 0.5** | | | 0,912 | 0,218 |
| **P80 - 0.05** | 0,190 | 0,853 | | |
| **P80 - 0.5** | | | 0,436 | 0,089 |

**Absolute abundance *Escherichia/Shigella***

| Vs. | RL - 0.005 | SL - 0.005 | RL - 0.05 | SL - 0.05 |
|---|---|---|---|---|
| **CMC - 0.05** | 0,123 | 0,481 | | |
| **CMC - 0.5** | | | **0,002** | **0,004** |
| **P80 - 0.05** | 0,143 | 0,912 | | |
| **P80 - 0.5** | | | **0,009** | **0,017** |

**Butyrate**

| Vs. | RL - 0.005 | SL - 0.005 | RL - 0.05 | SL - 0.05 |
|---|---|---|---|---|
| **CMC - 0.05** | **0,013** | 0,720 | | |
| **CMC - 0.5** | | | **0,000** | **0,000** |
| **P80 - 0.05** | **0,011** | 0,853 | | |
| **P80 - 0.5** | | | **0,000** | **0,000** |

**Relative abundance *Escherichia/Shigella***

| Vs. | RL - 0.005 | SL - 0.005 | RL - 0.05 | SL - 0.05 |
|---|---|---|---|---|
| **CMC - 0.05** | 0,063 | 0,579 | | |
| **CMC - 0.5** | | | **0,000** | **0,000** |
| **P80 - 0.05** | 0,063 | 0,853 | | |
| **P80 - 0.5** | | | **0,000** | **0,001** |

**Intact cell concentration**

| Vs. | RL - 0.005 | SL - 0.005 | RL - 0.05 | SL - 0.05 |
|---|---|---|---|---|
| **CMC - 0.05** | **0,000** | 0,102 | | |
| **CMC - 0.5** | | | **0,000** | **0,000** |
| **P80 - 0.05** | **0,000** | 0,620 | | |
| **P80 - 0.5** | | | **0,000** | **0,000** |
