## Supplementary figures and images for "Dietary emulsifiers alter composition and activity of the human gut microbiota *in vitro*, irrespective of chemical or natural emulsifier origin"

### Aerobic.pdf

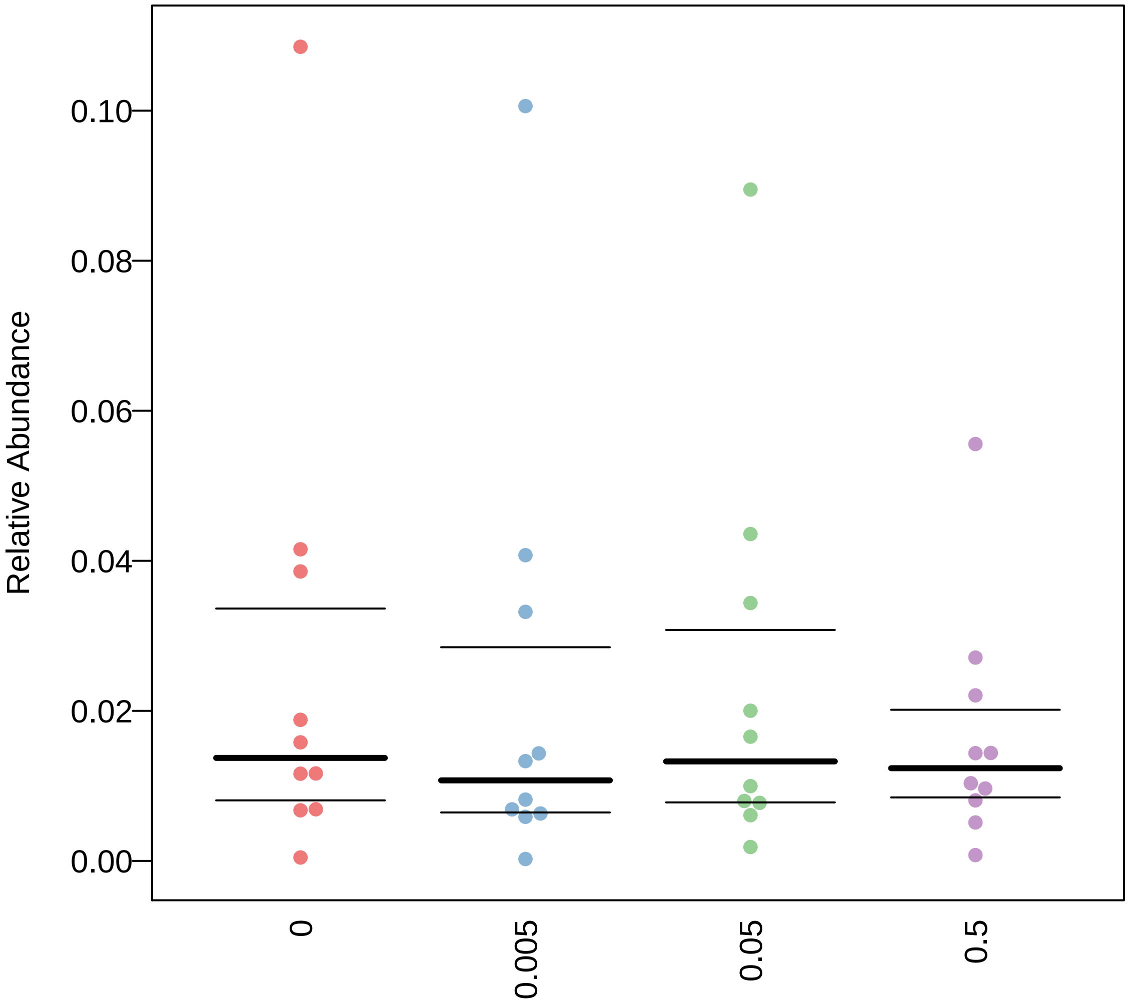

### Aerobic.pdf

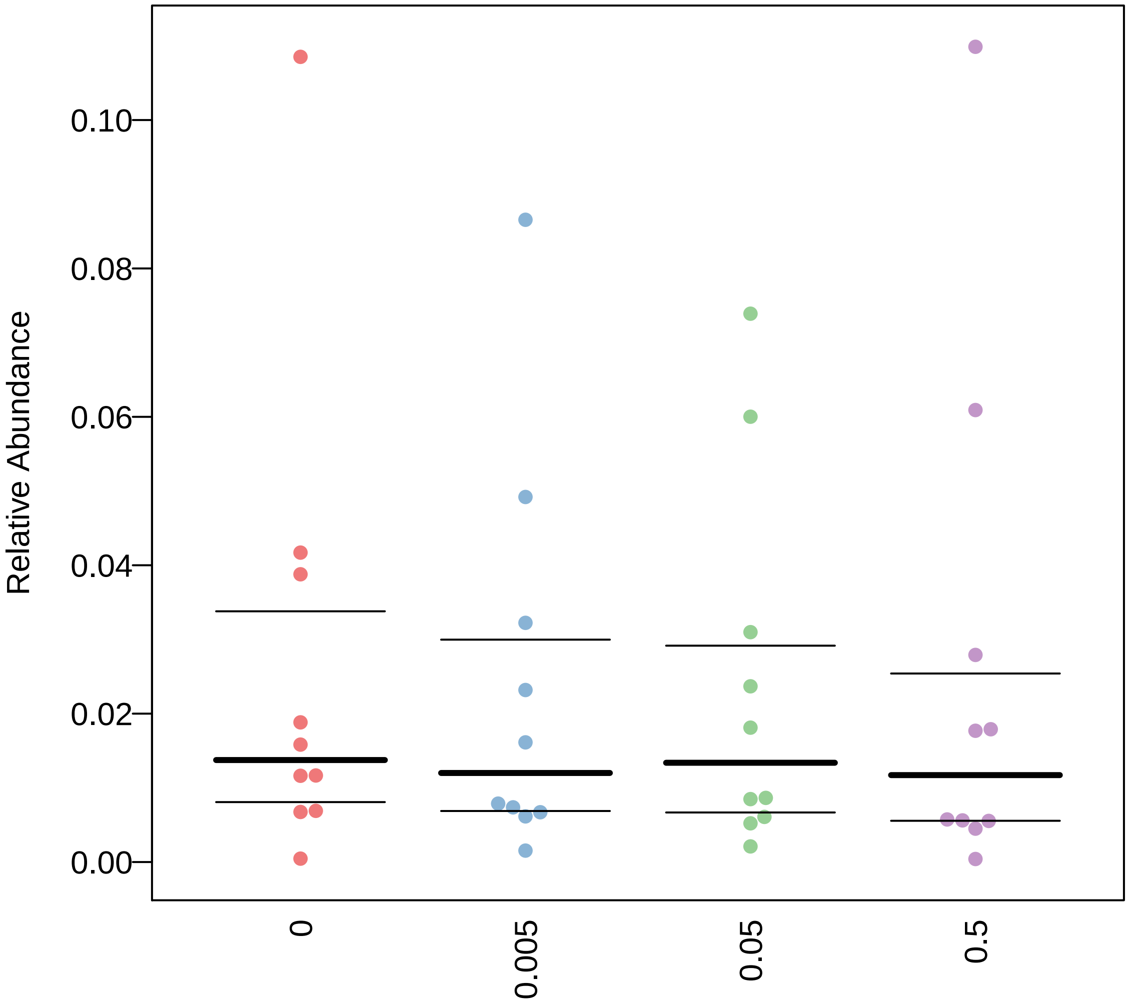

### Aerobic.pdf

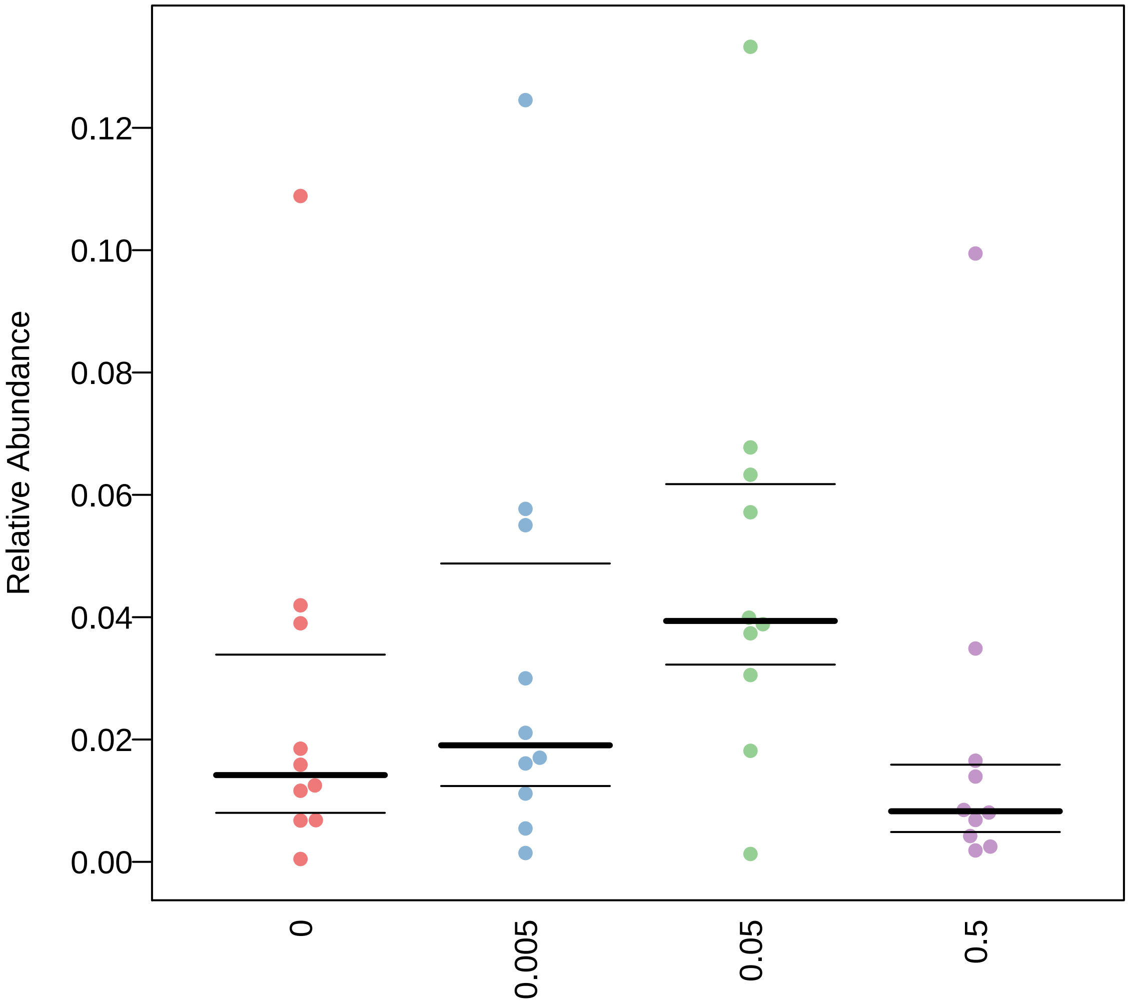

### Aerobic.pdf

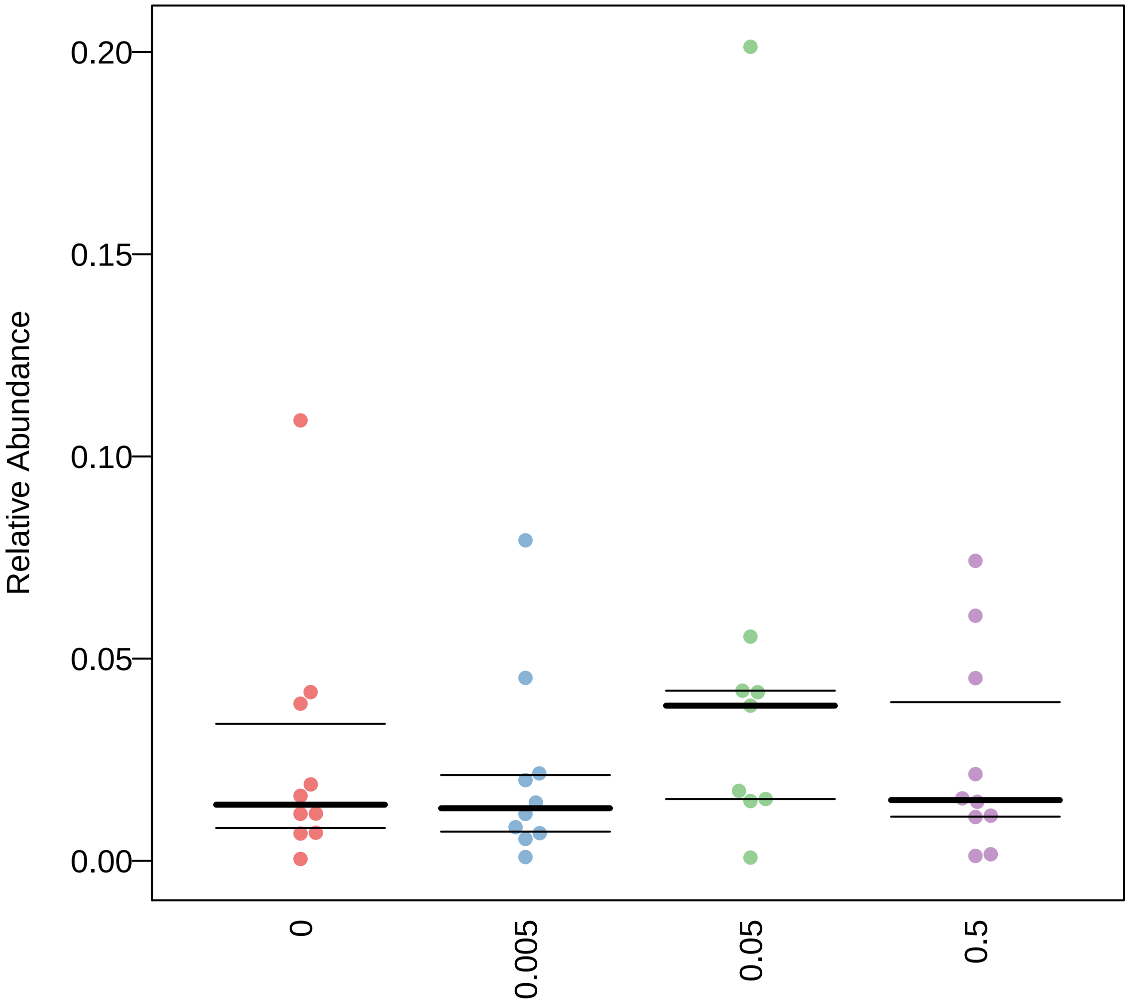

### Aerobic.pdf

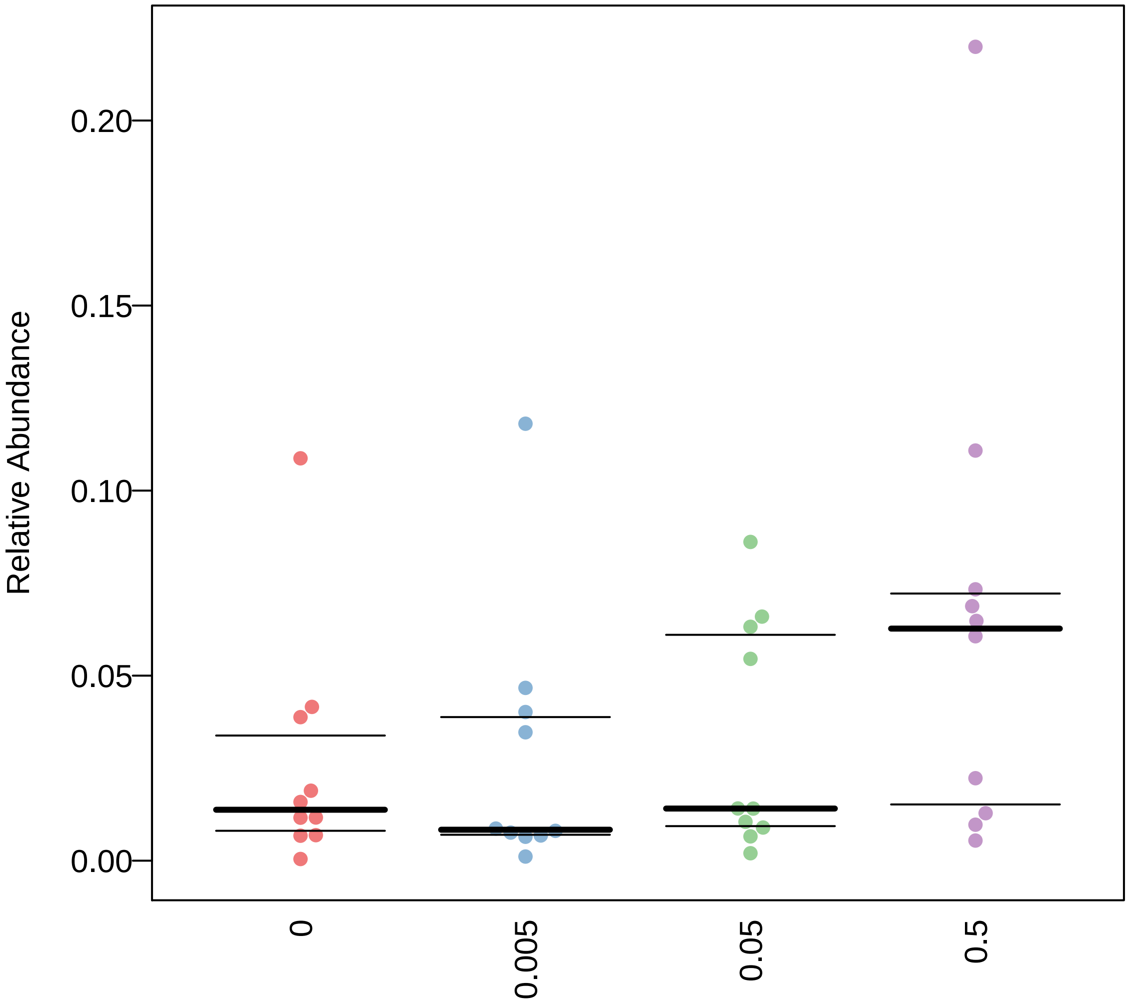

### Anaerobic.pdf

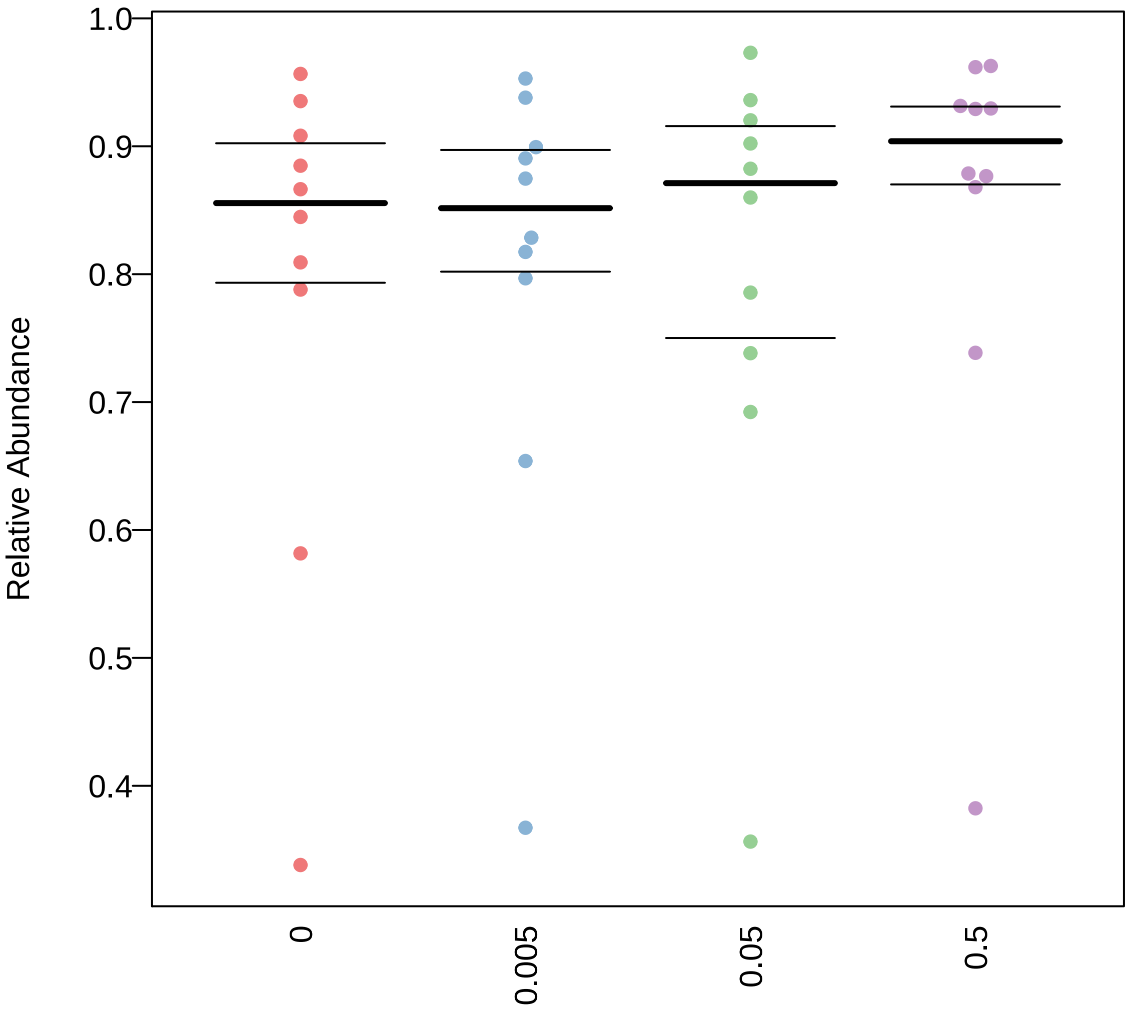

### Anaerobic.pdf

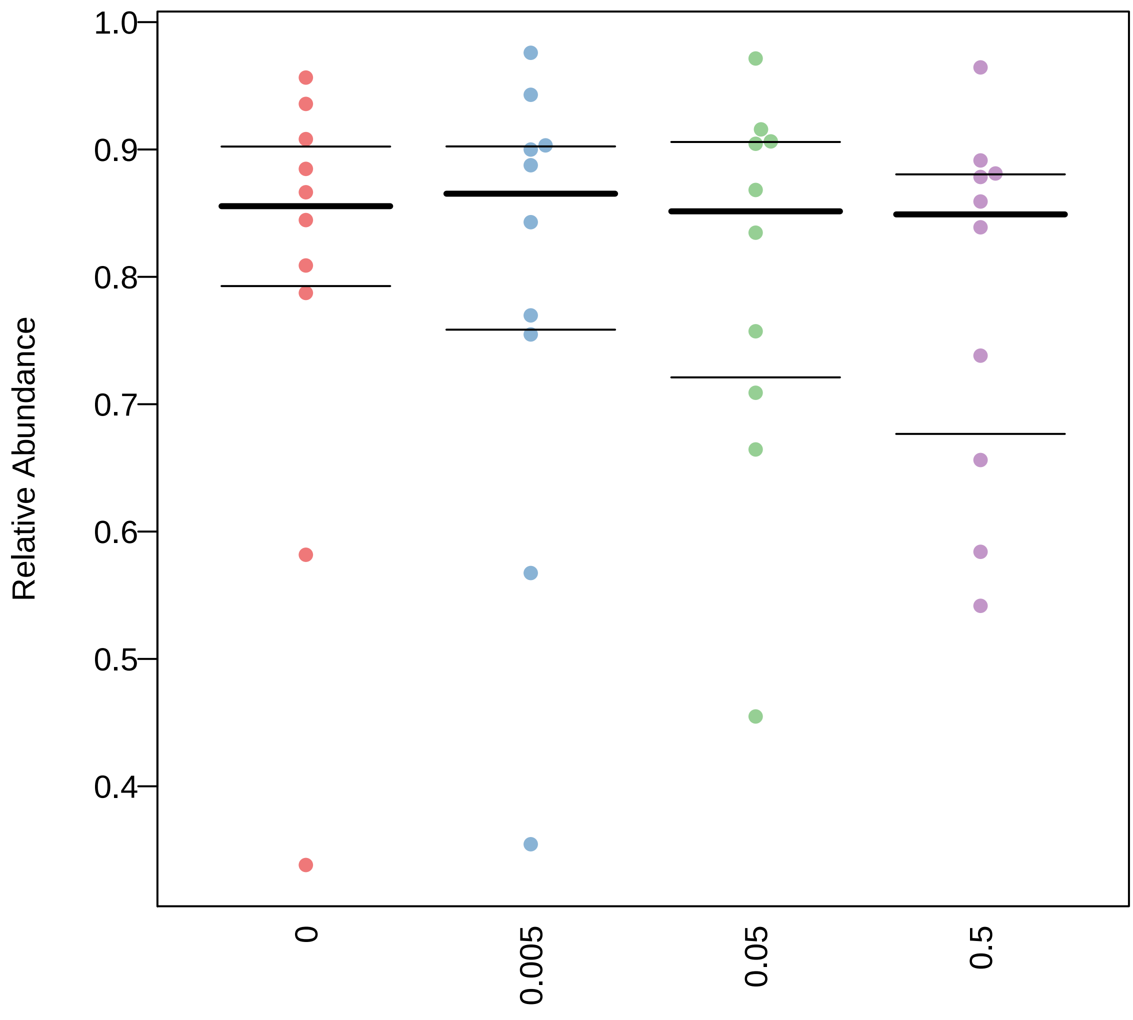

### Anaerobic.pdf

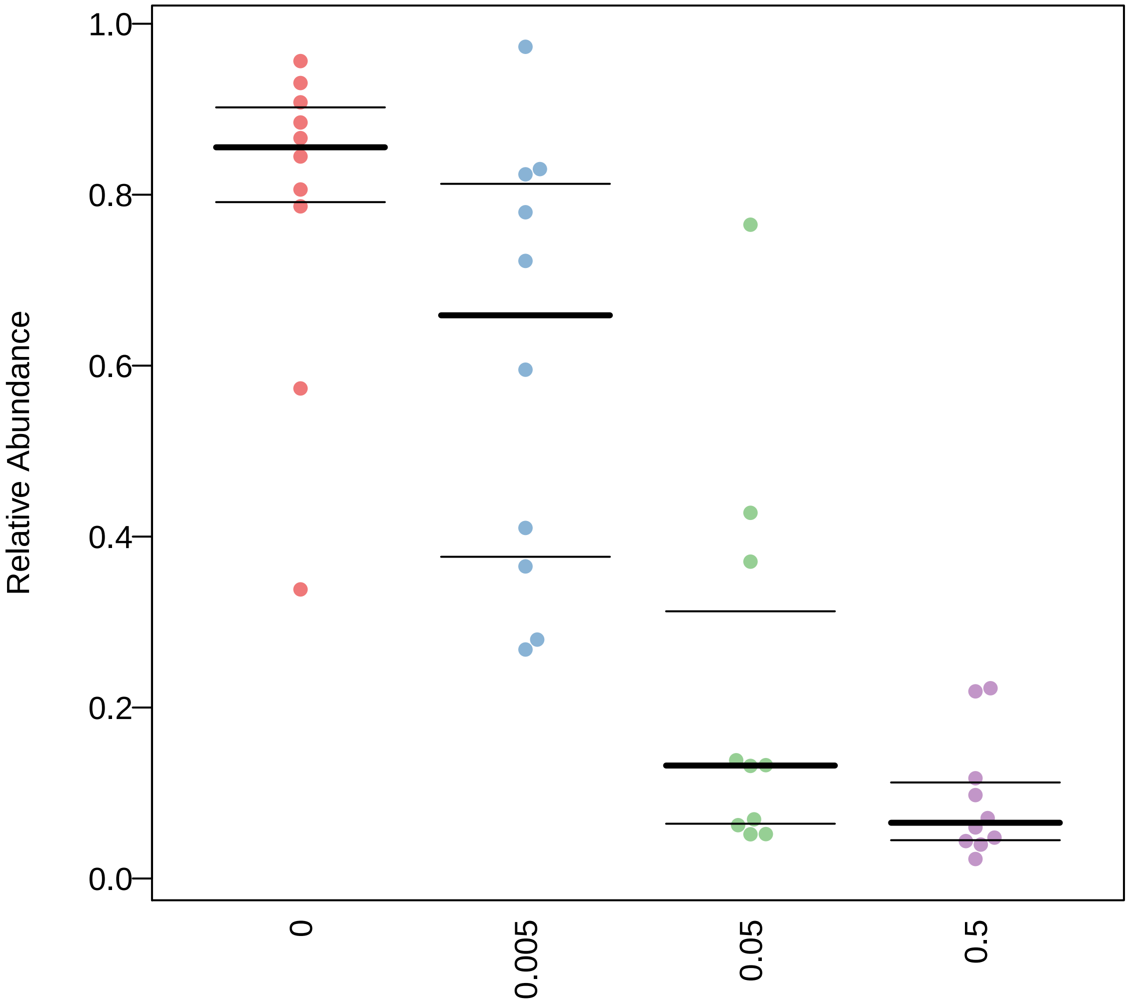

### Anaerobic.pdf

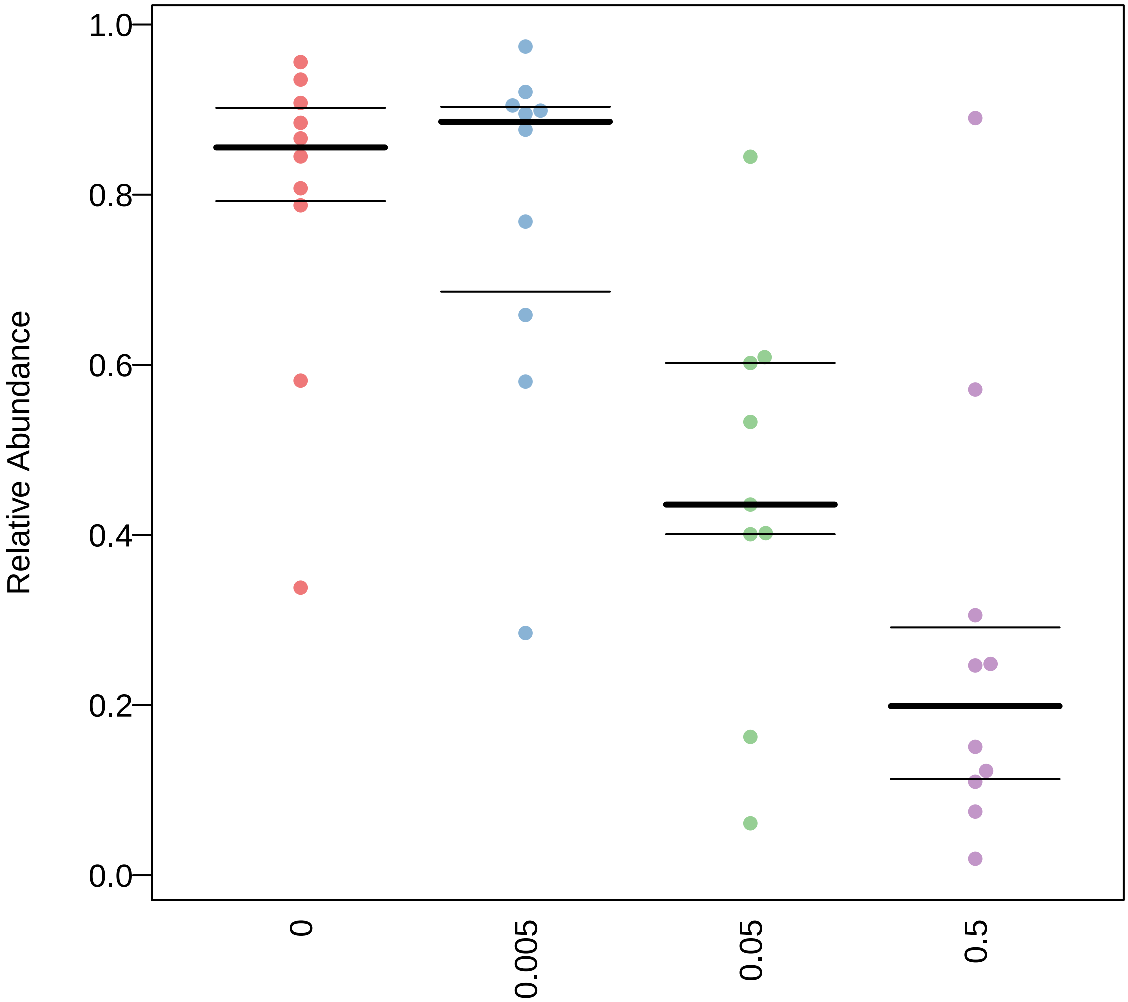

### Anaerobic.pdf

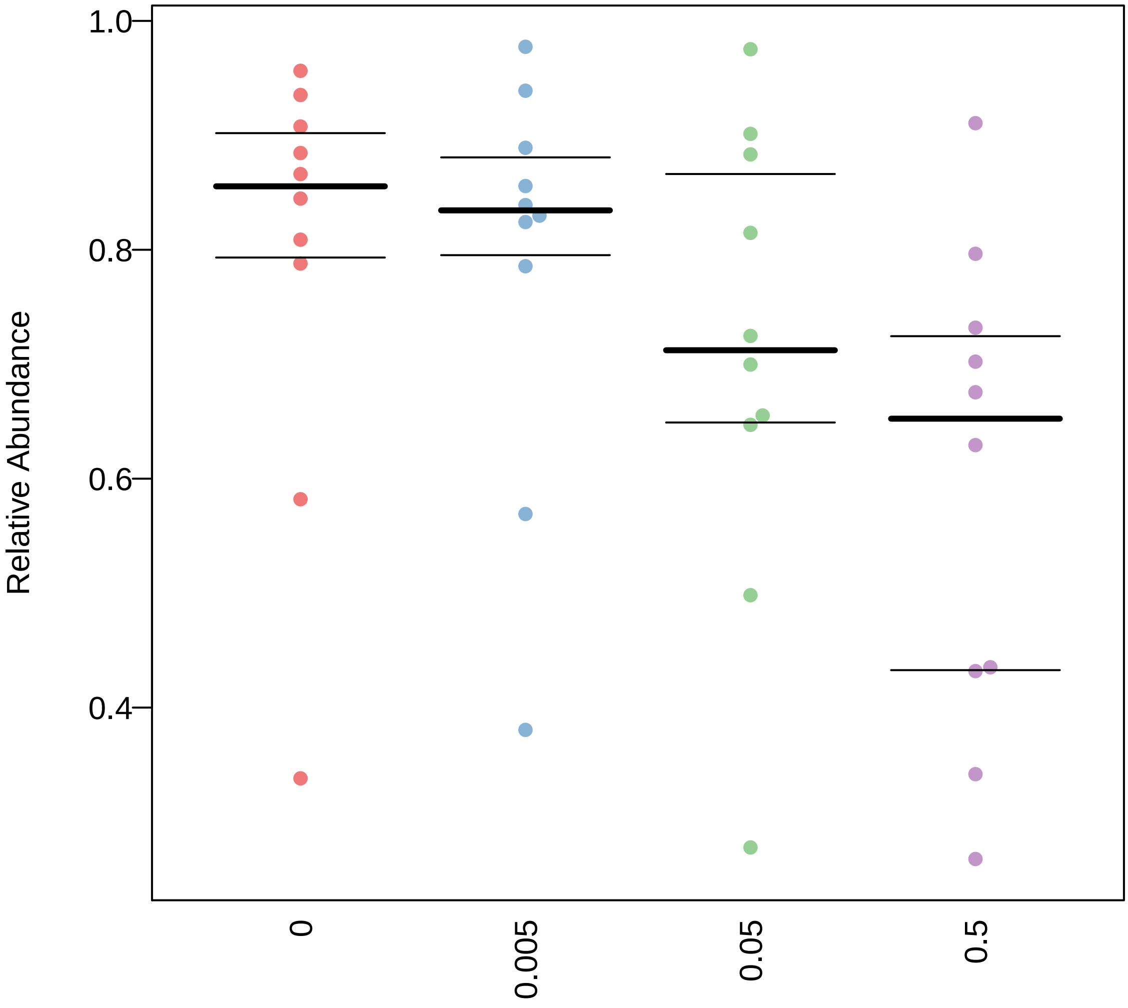

### Contains_Mobile_Elements.pdf

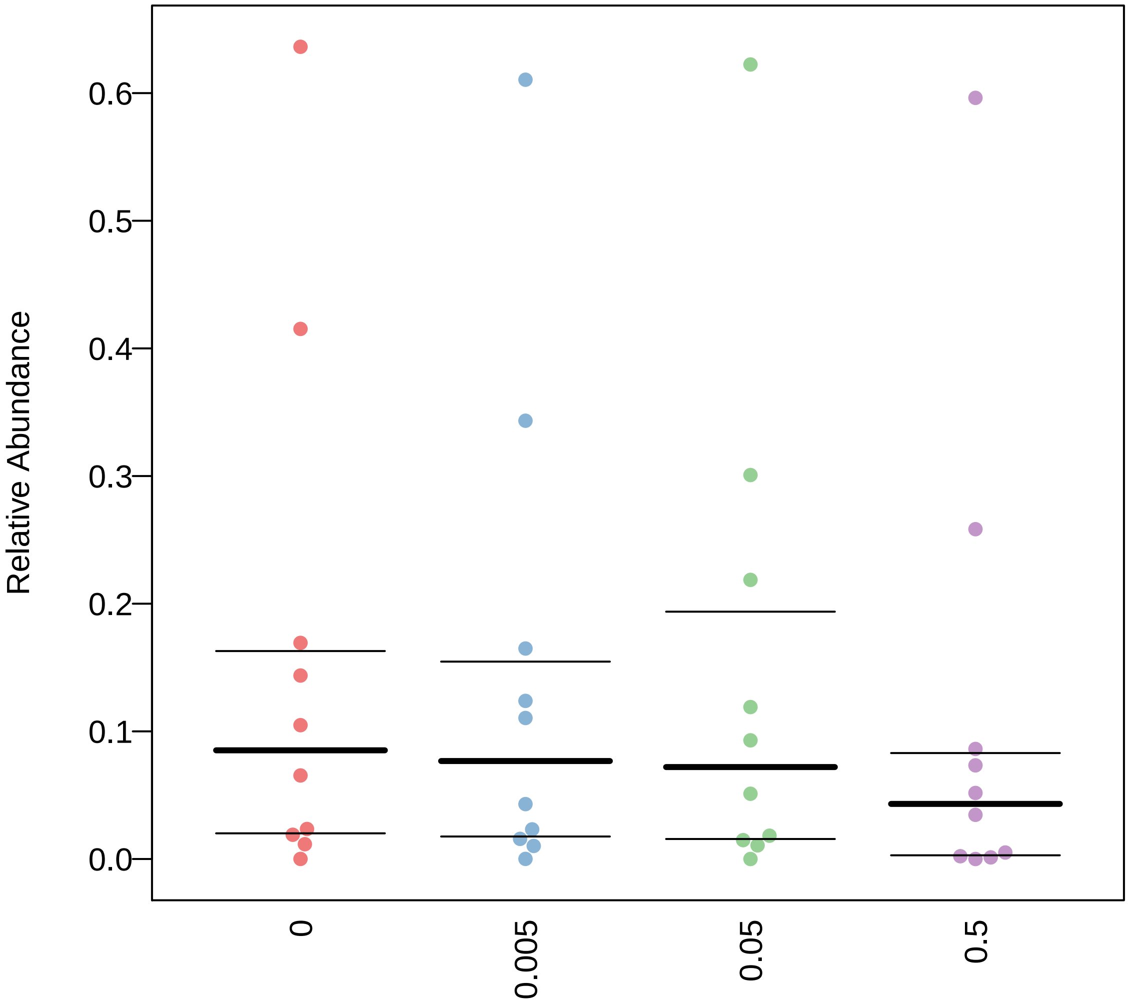

### Contains_Mobile_Elements.pdf

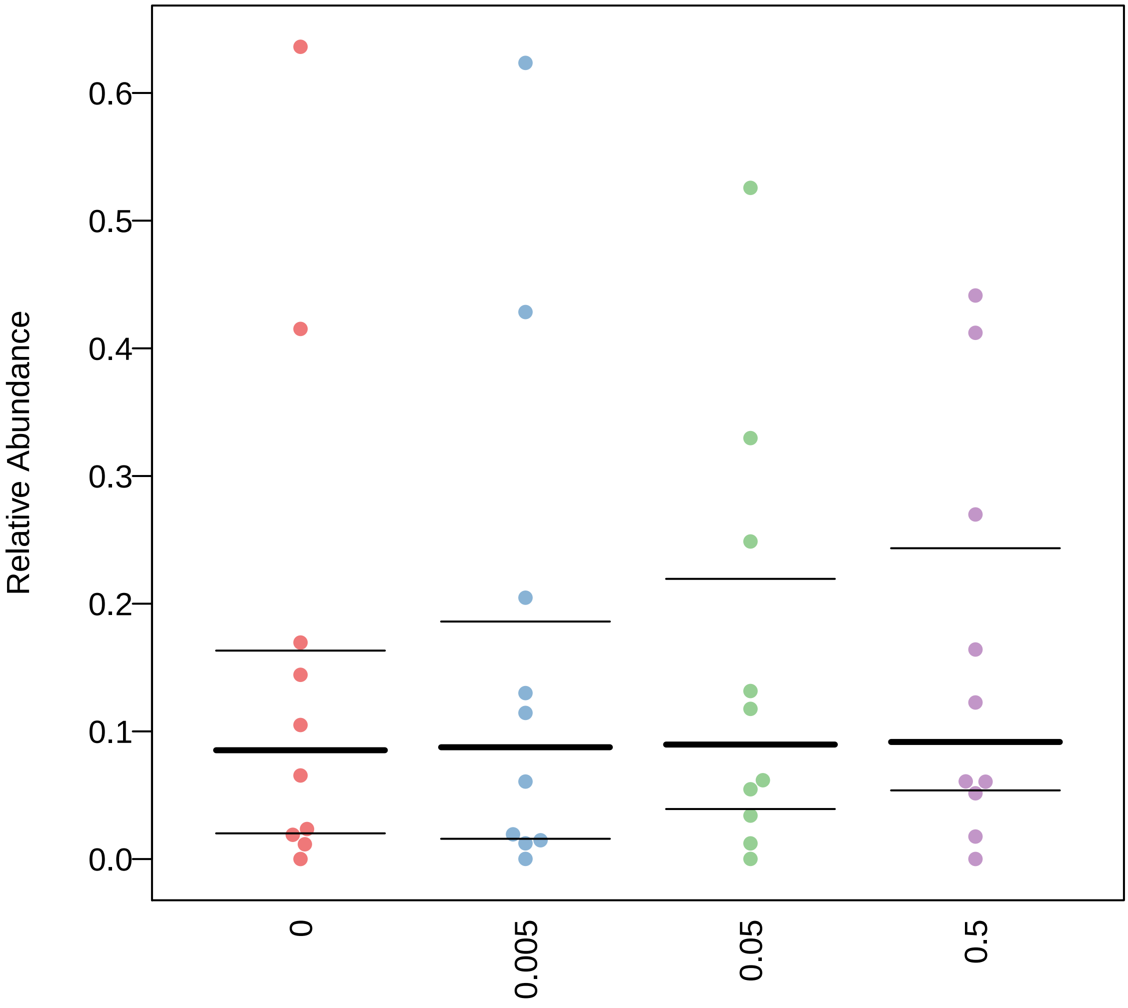

### Contains_Mobile_Elements.pdf

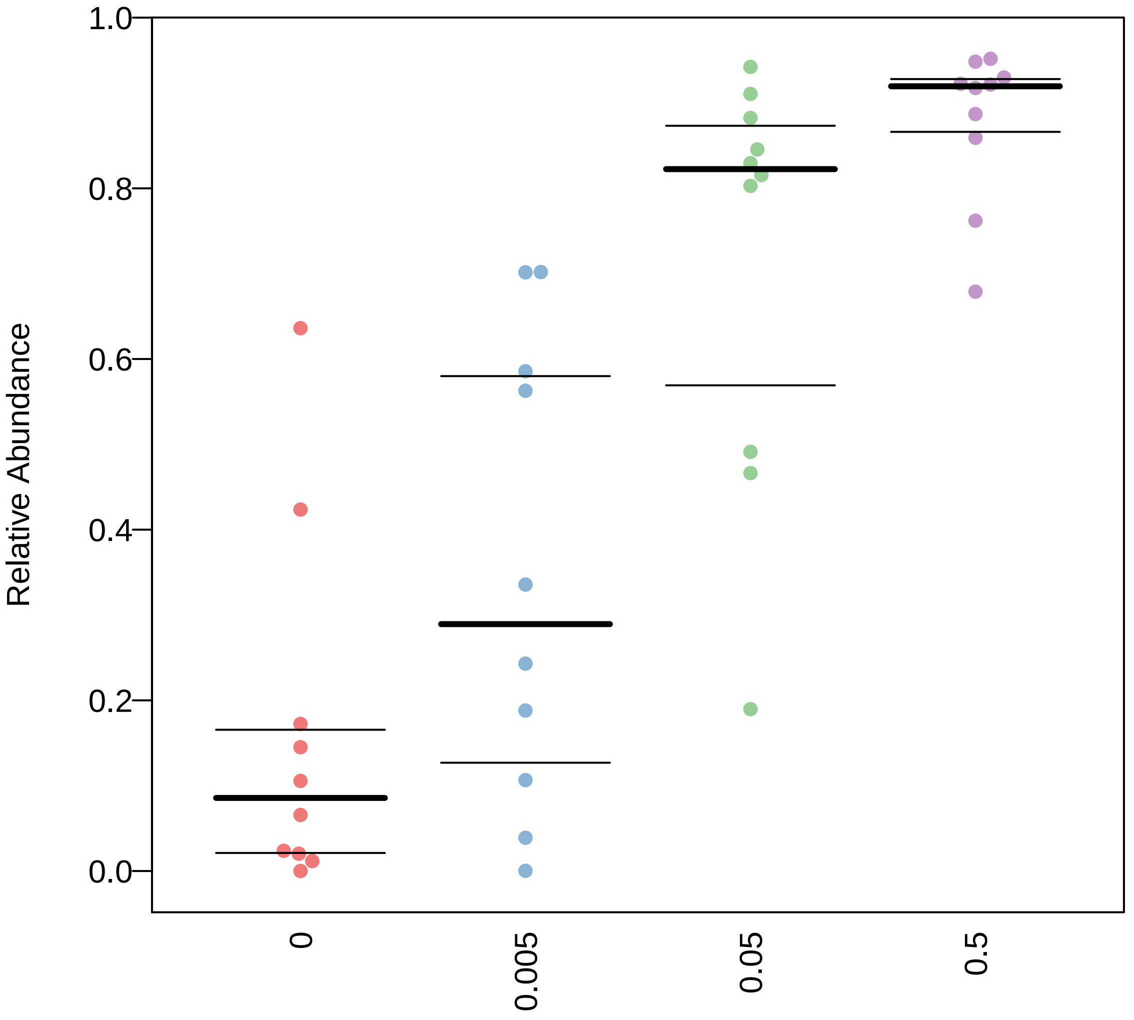

### Contains_Mobile_Elements.pdf

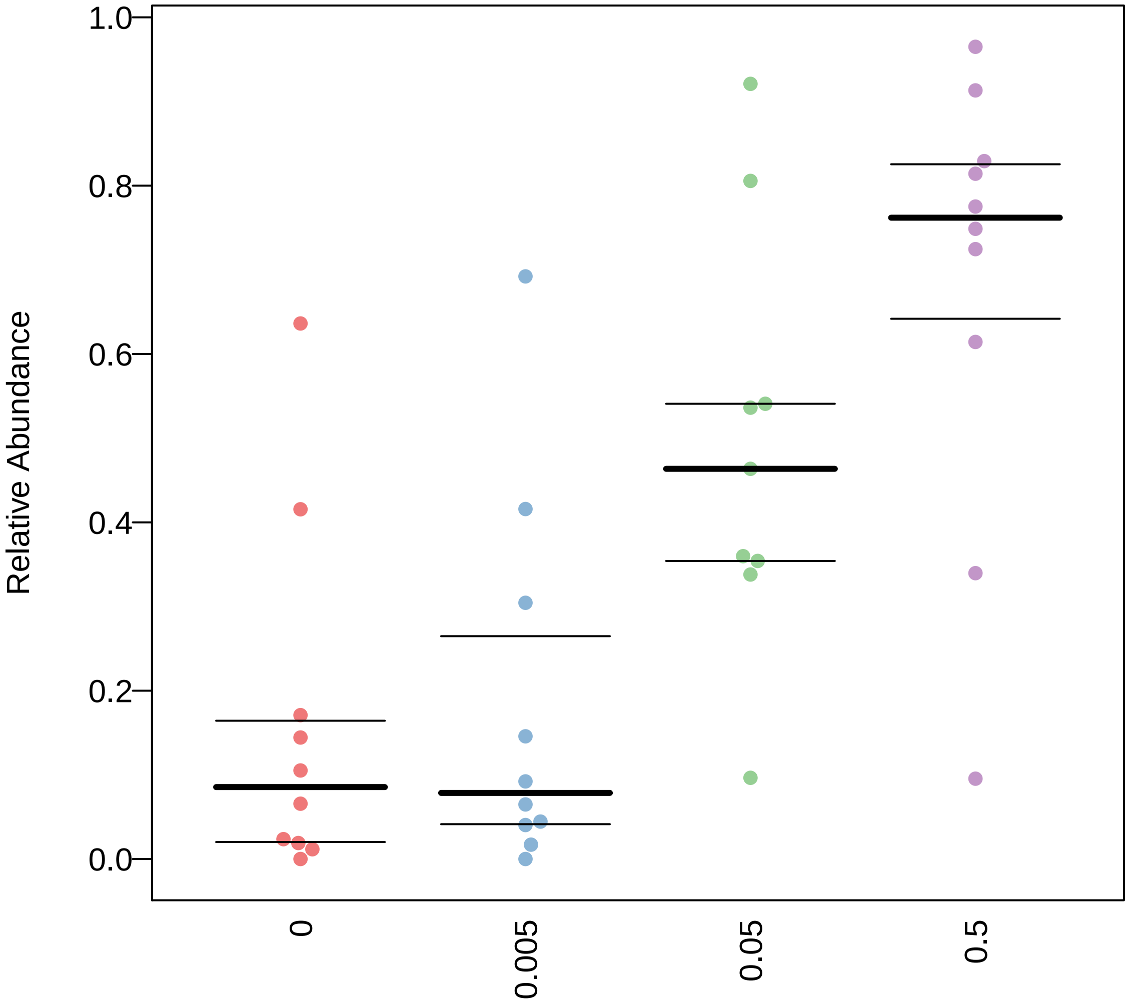

### Contains_Mobile_Elements.pdf

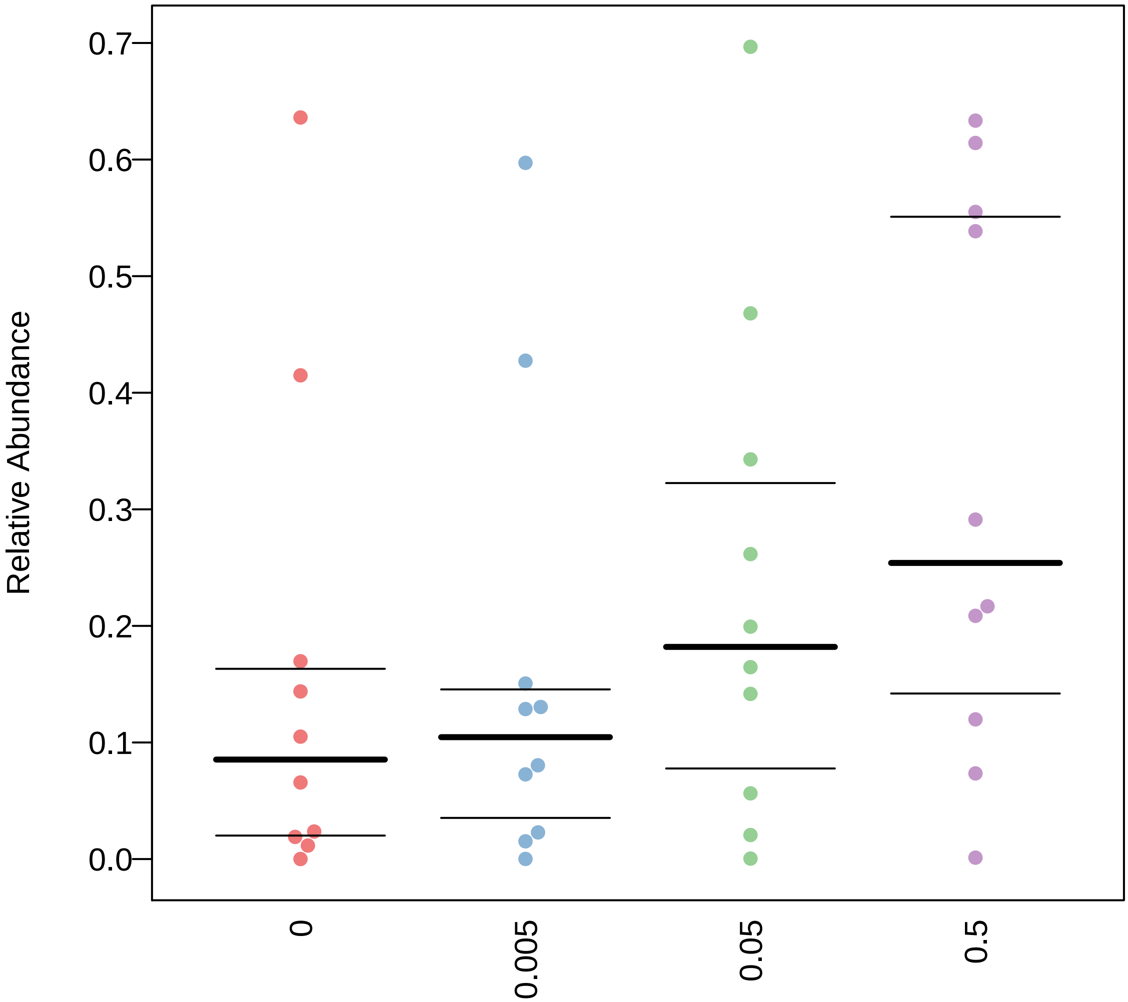

### Facultatively_Anaerobic.pdf

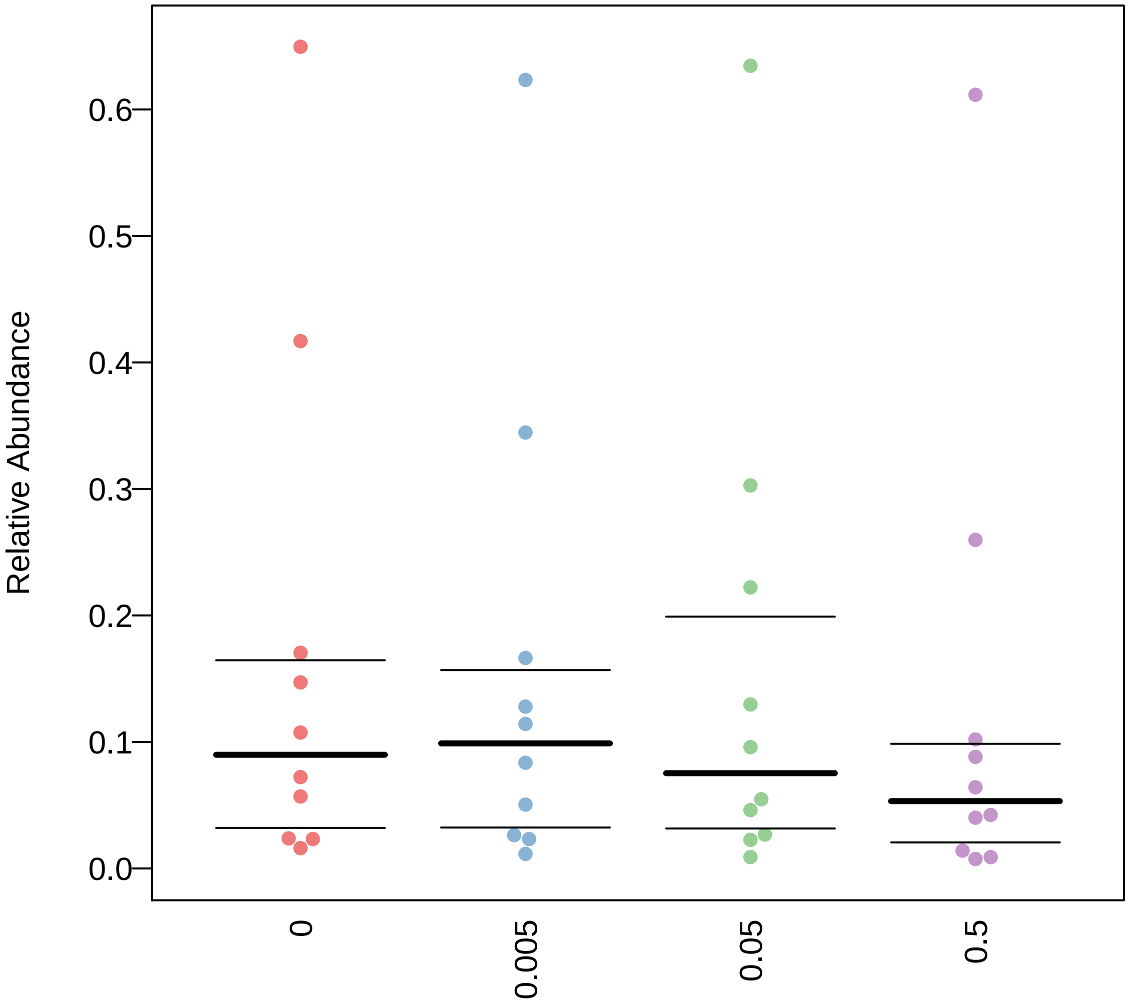

### Facultatively_Anaerobic.pdf

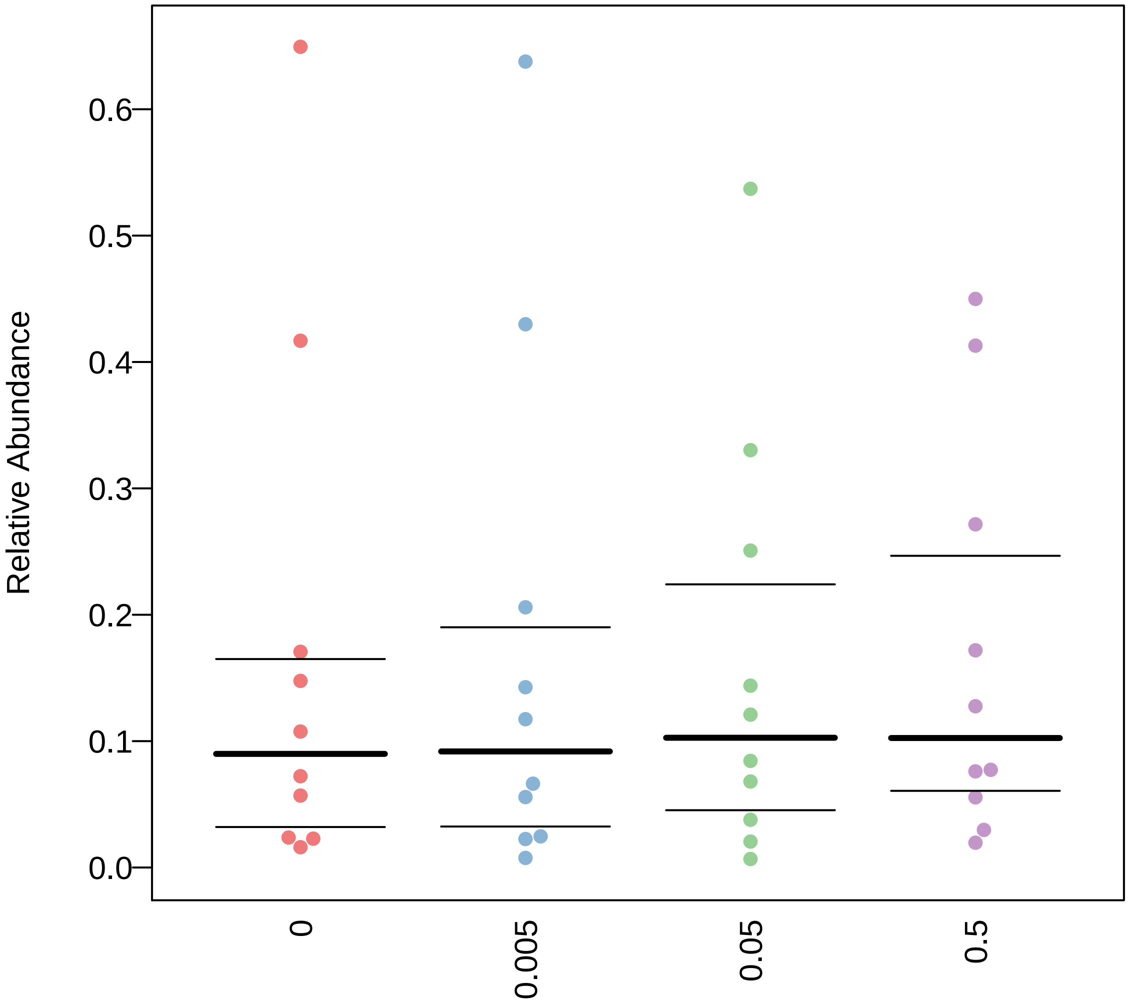

### Facultatively_Anaerobic.pdf

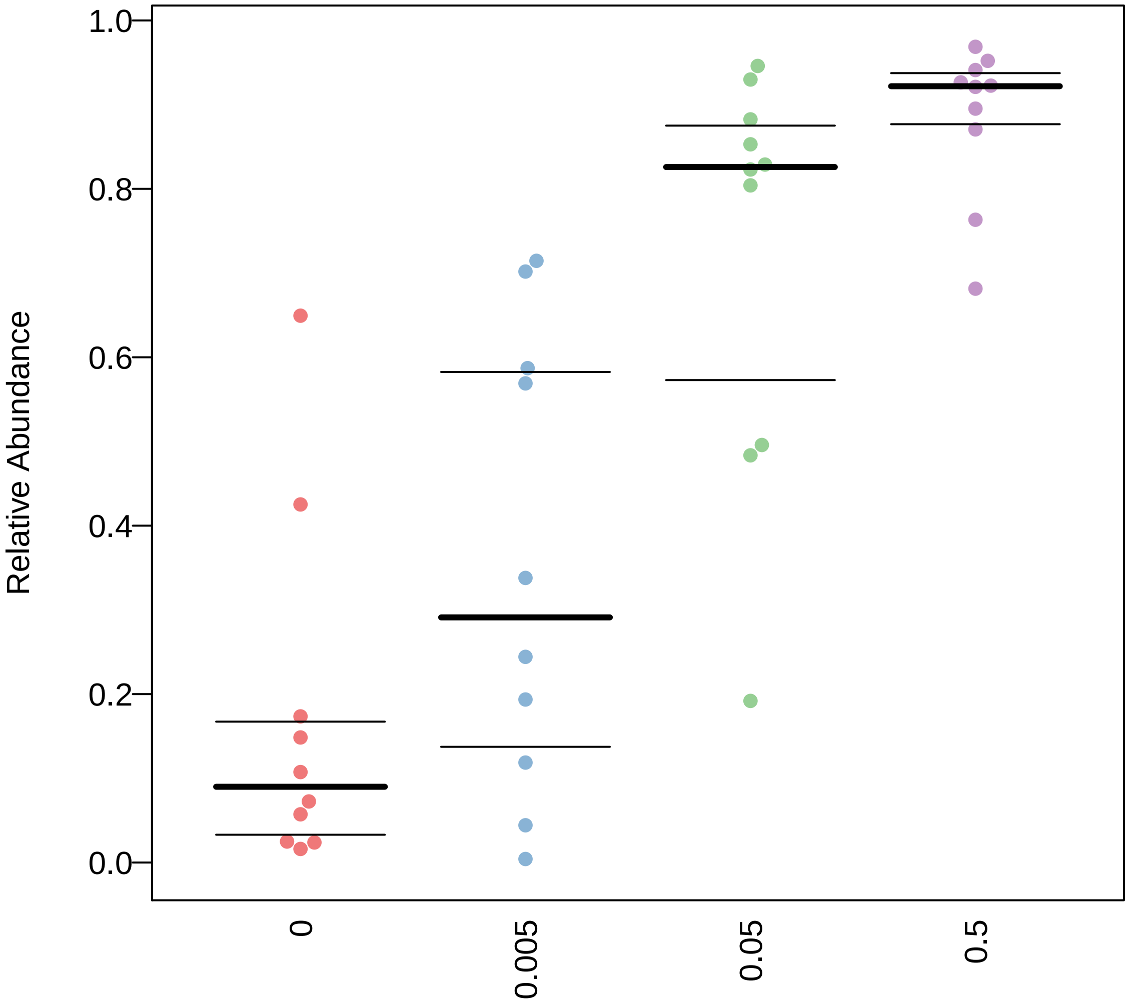

### Facultatively_Anaerobic.pdf

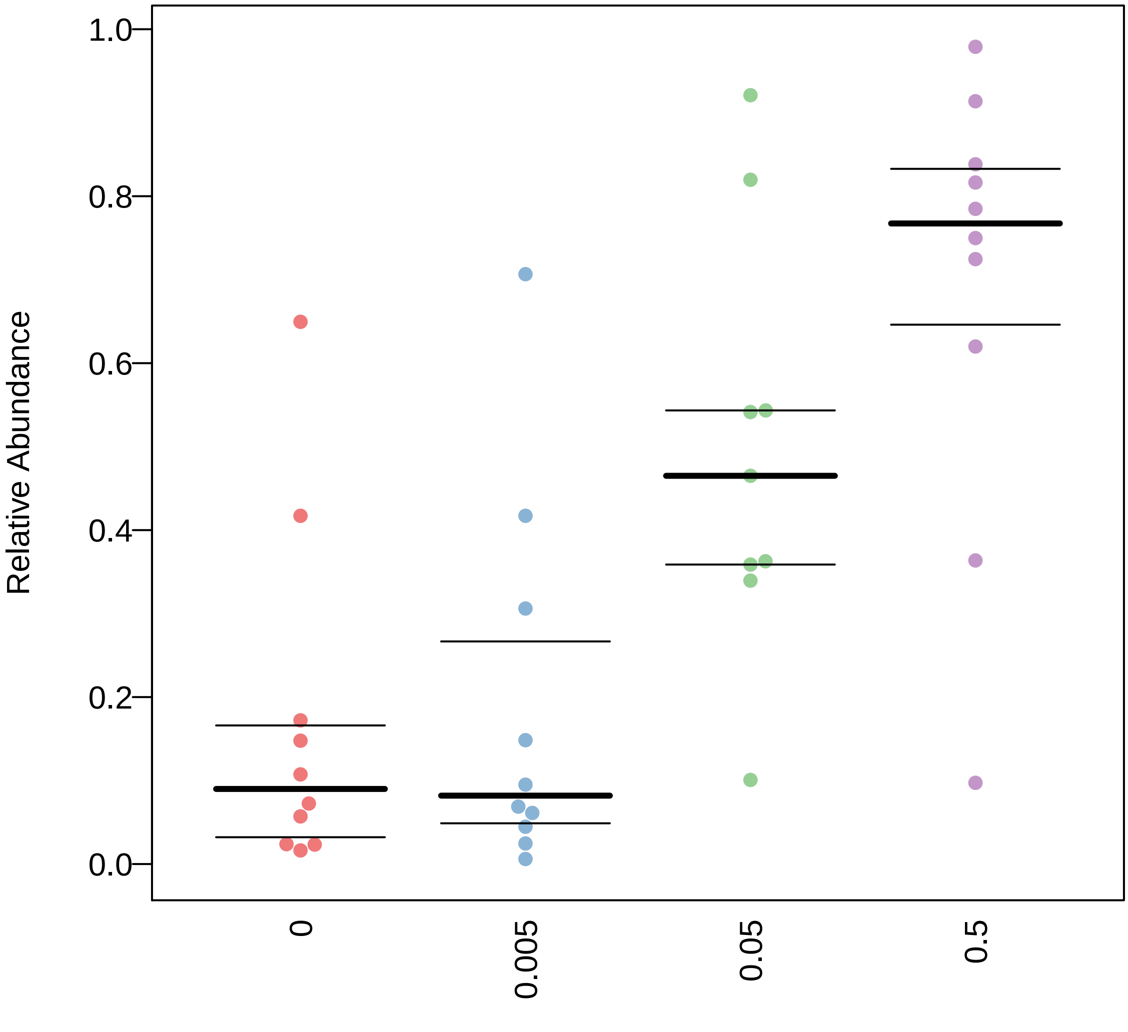

### Facultatively_Anaerobic.pdf

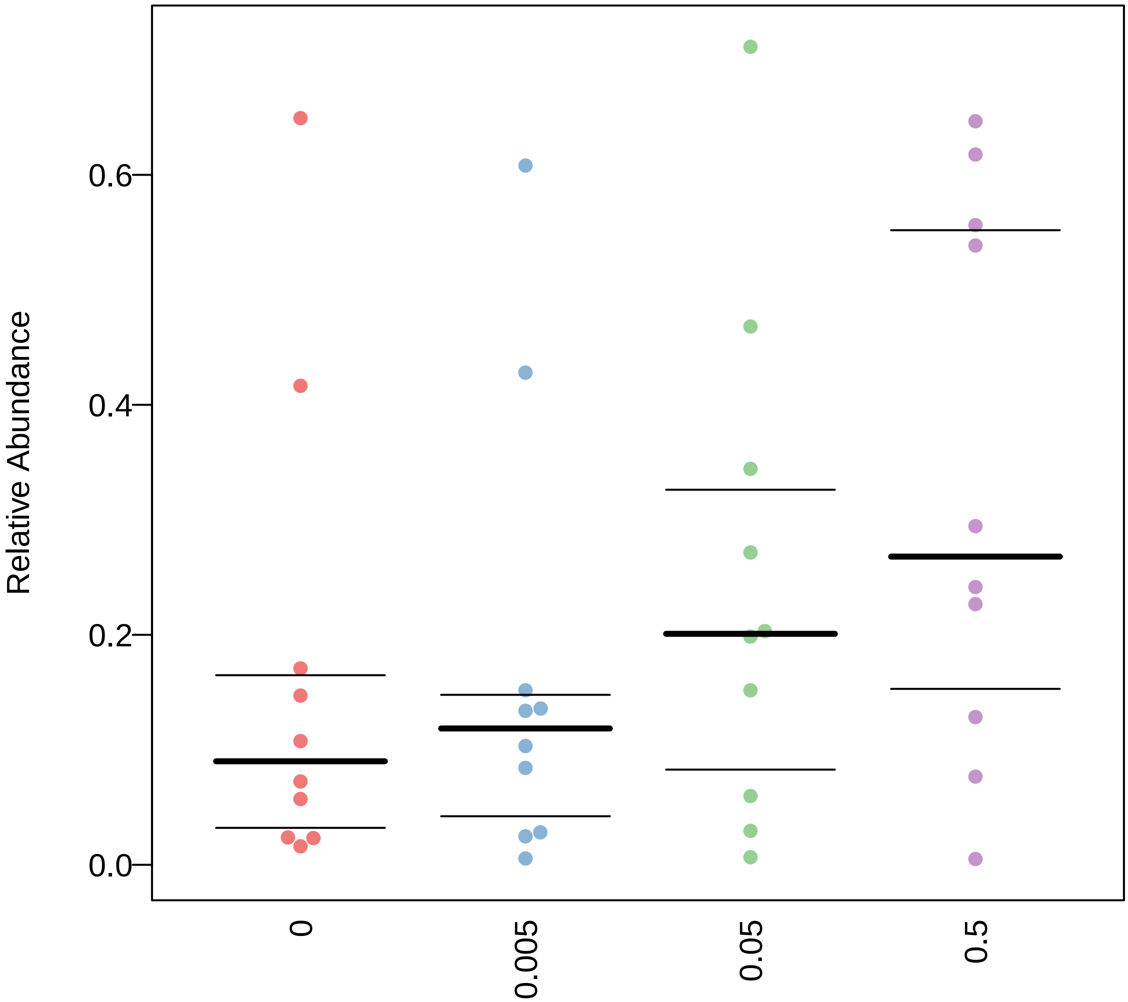

### Forms_Biofilms.pdf

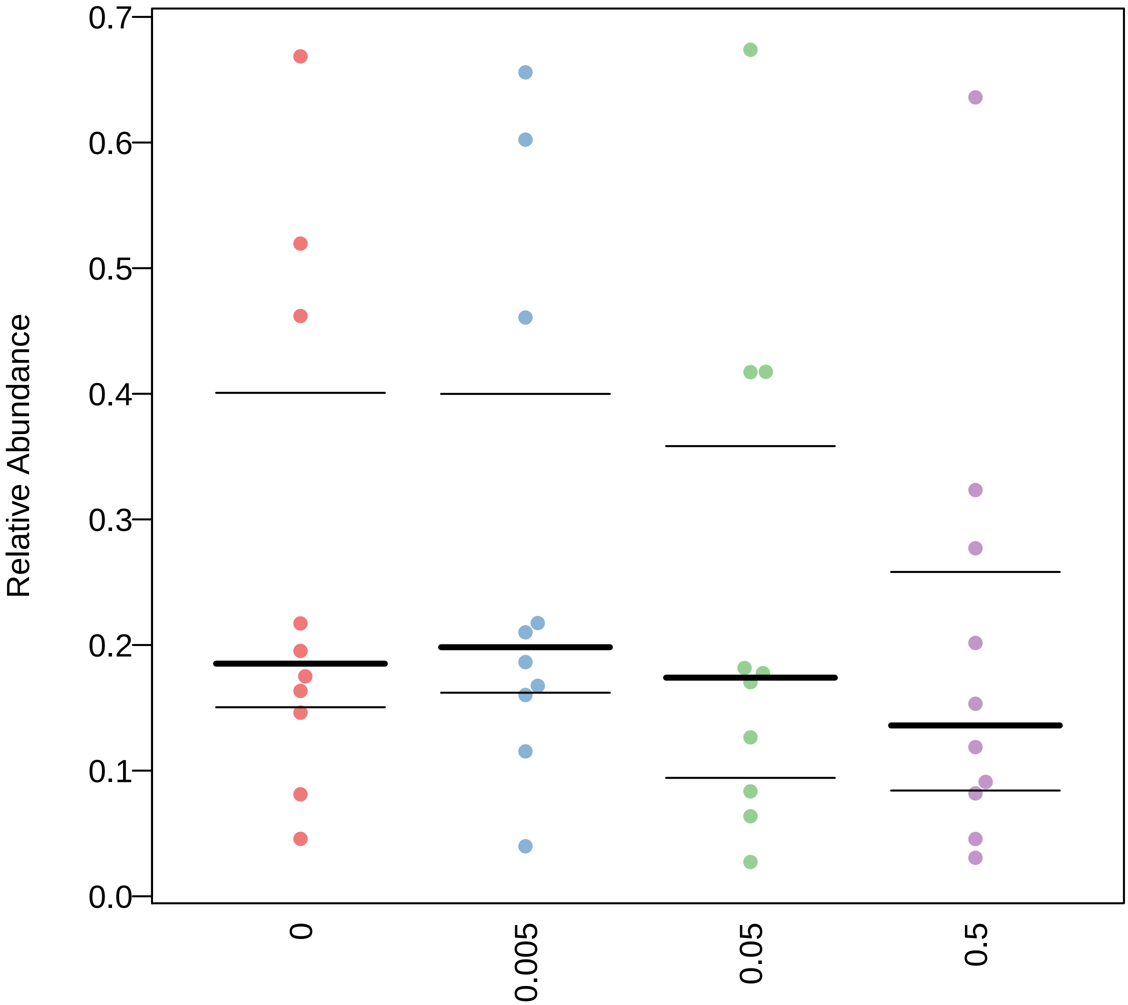

### Forms_Biofilms.pdf

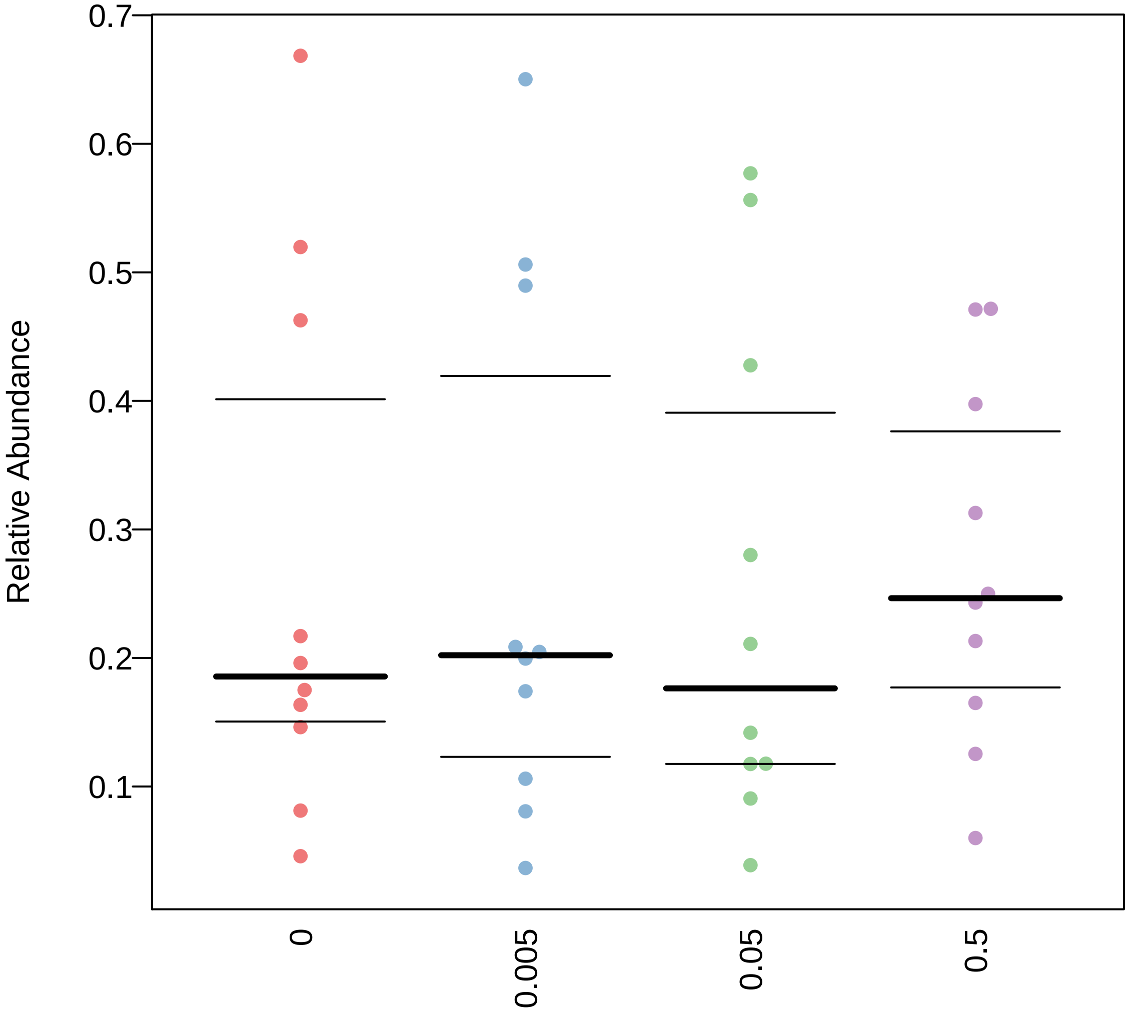

### Forms_Biofilms.pdf

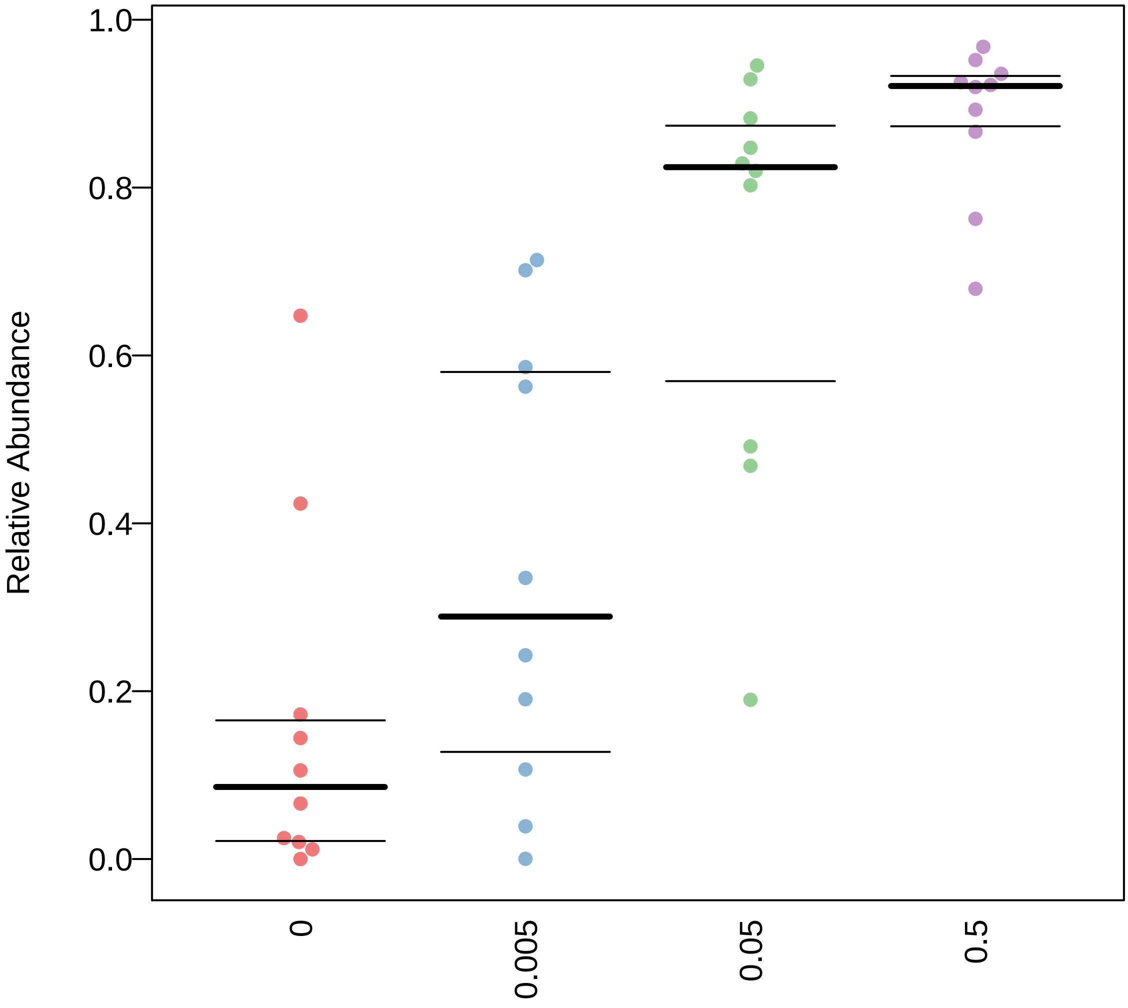

### Forms_Biofilms.pdf

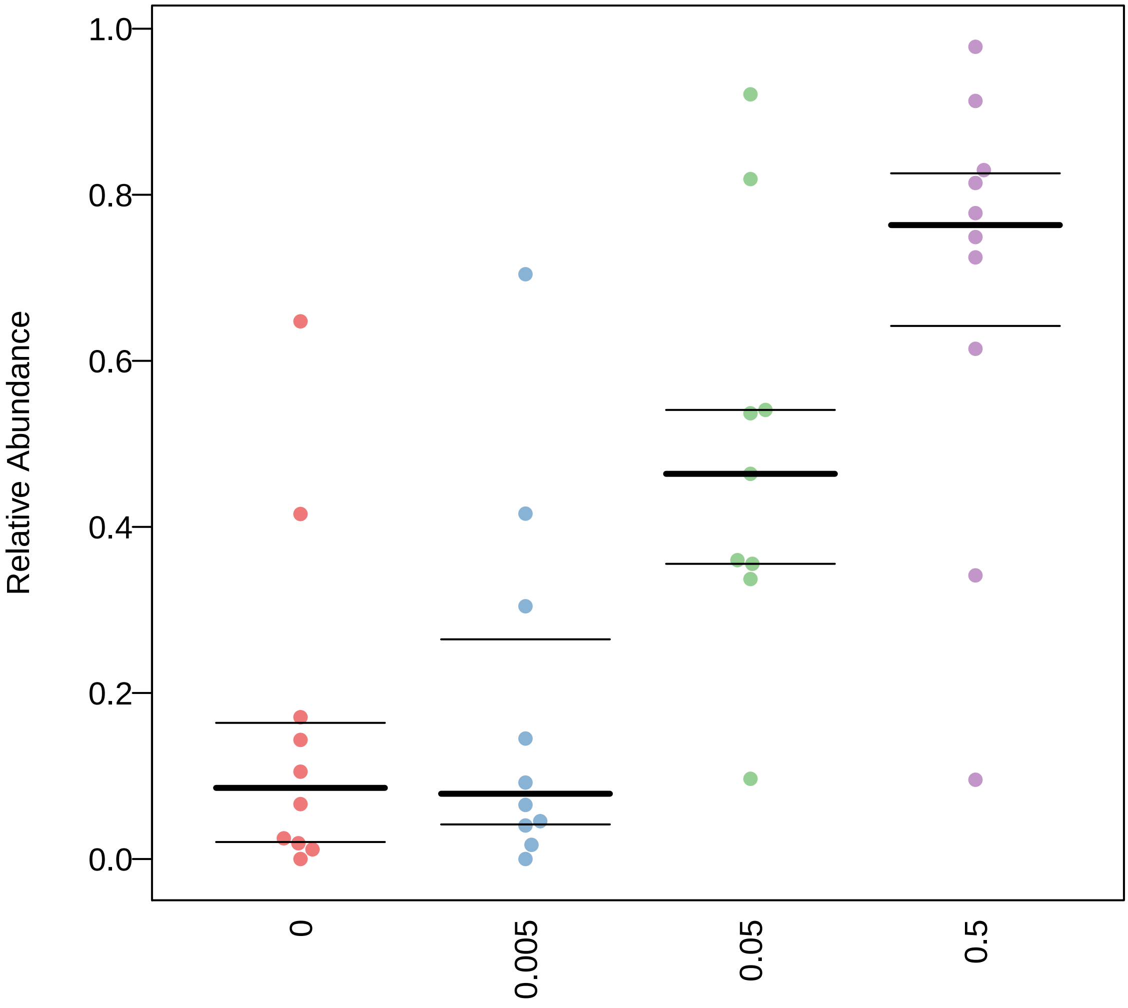

### Forms_Biofilms.pdf

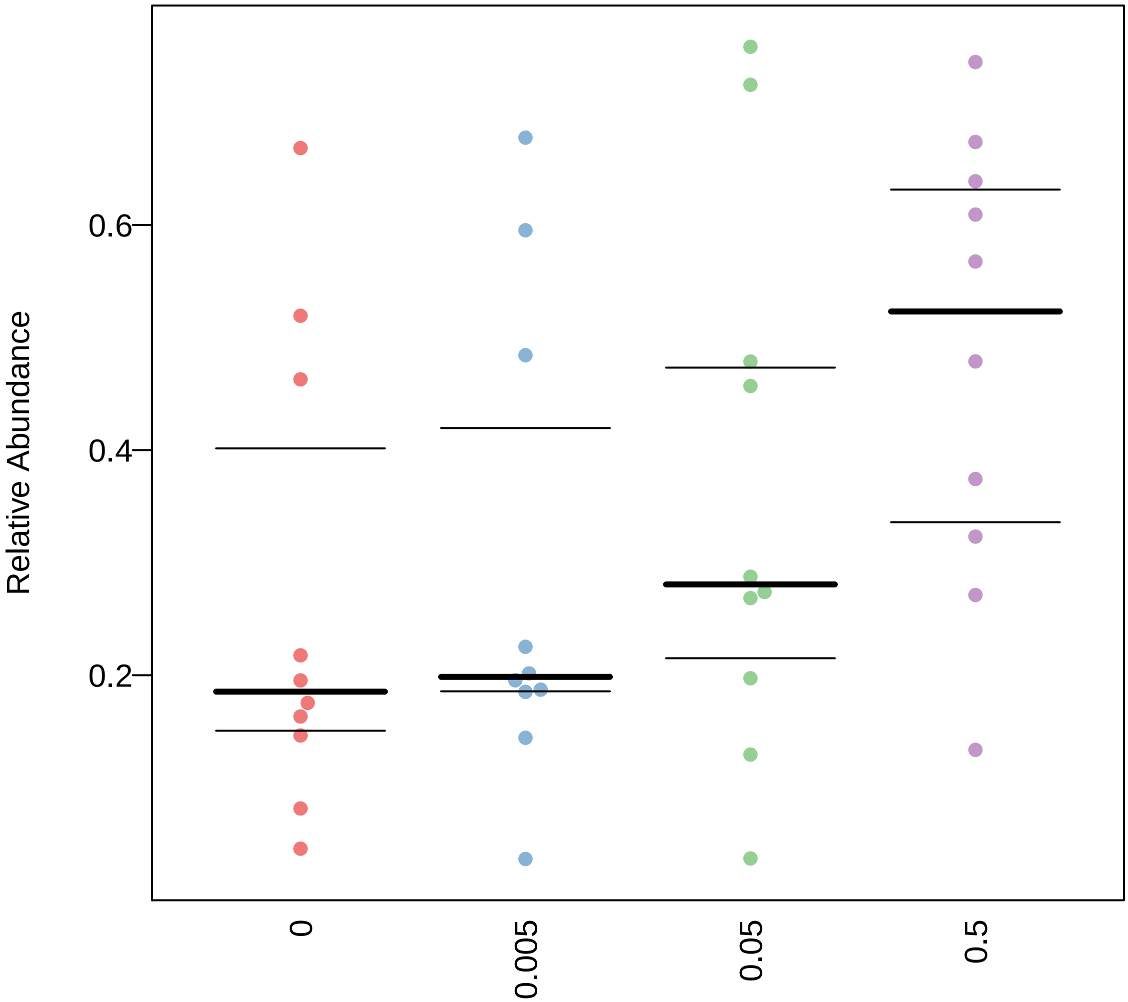

### Gram_Negative.pdf

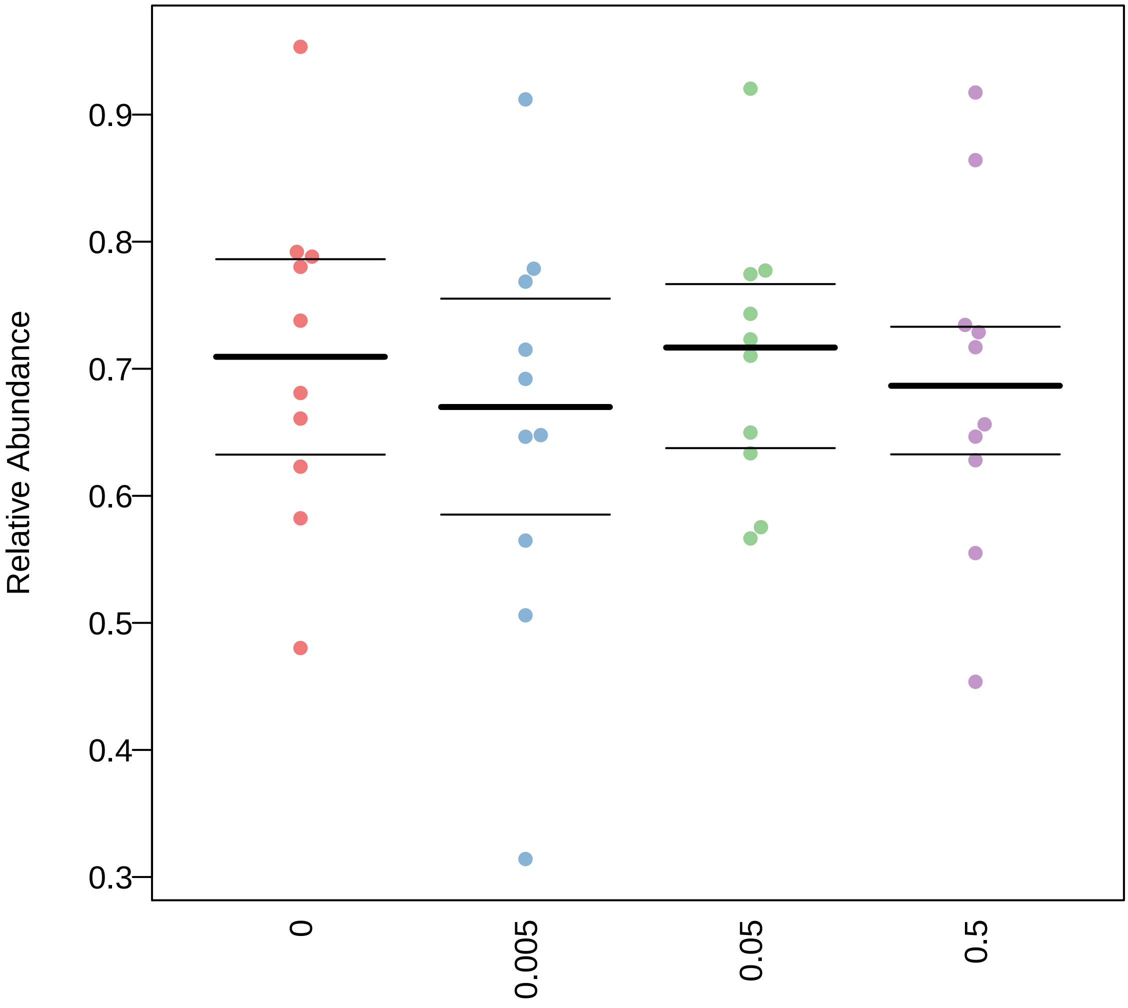

### Gram_Negative.pdf

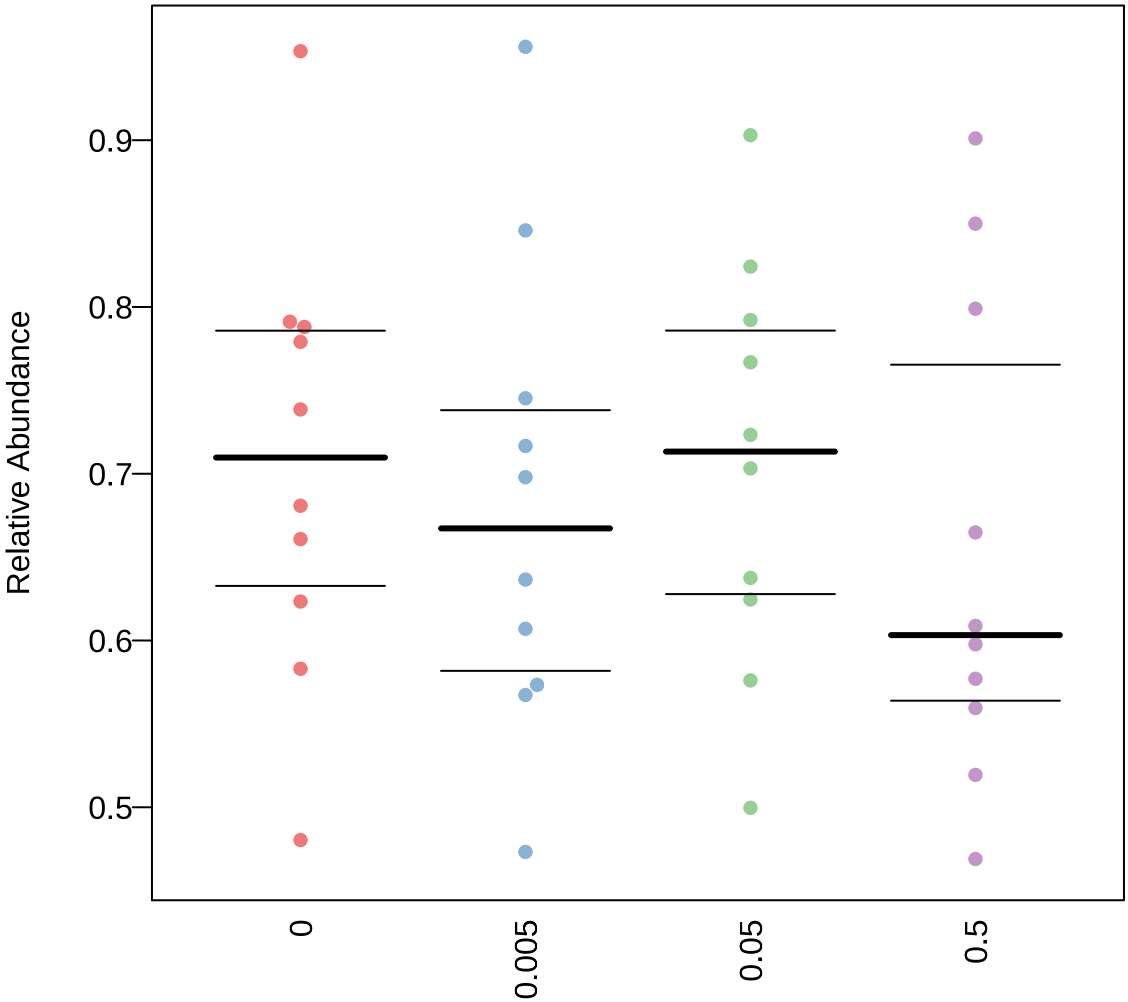

### Gram_Negative.pdf

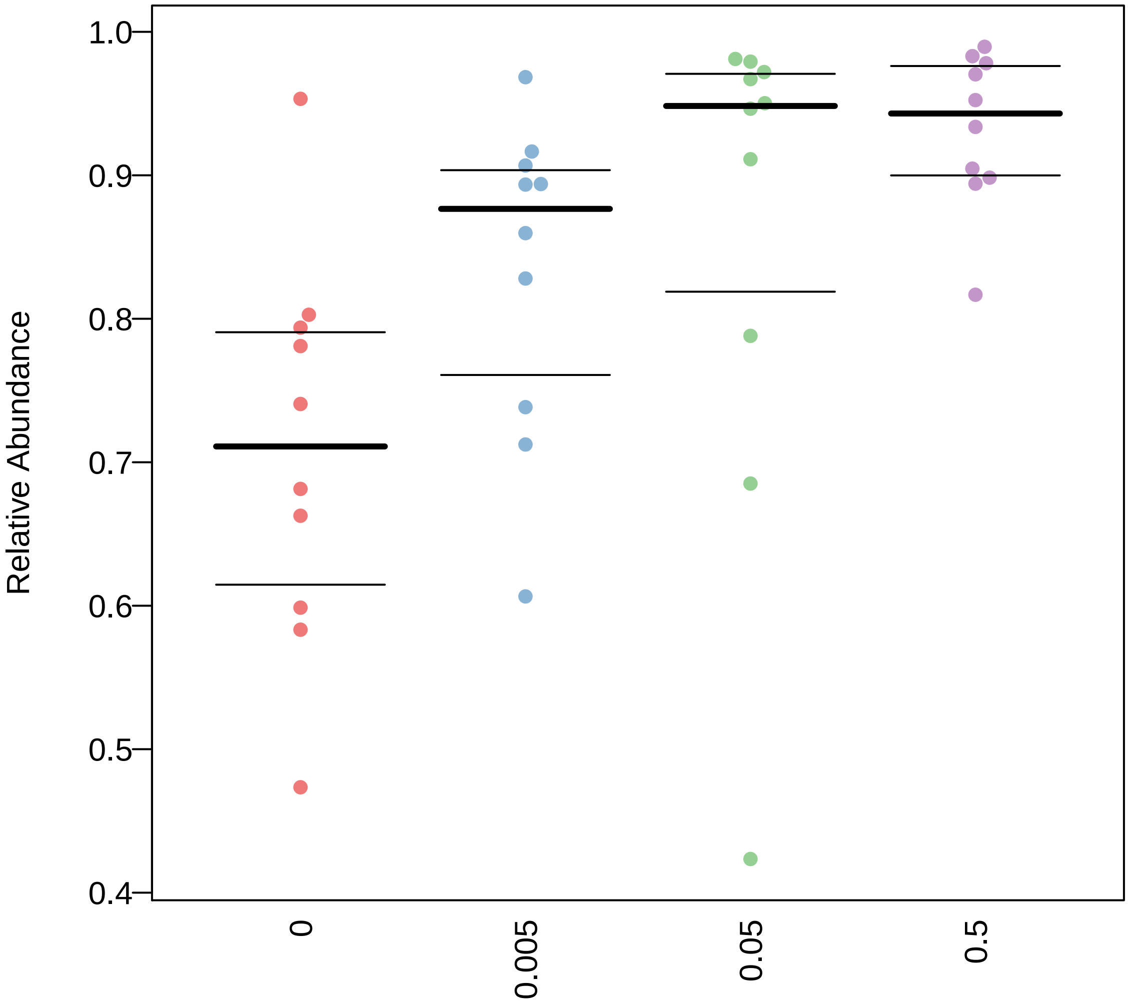

### Gram_Negative.pdf

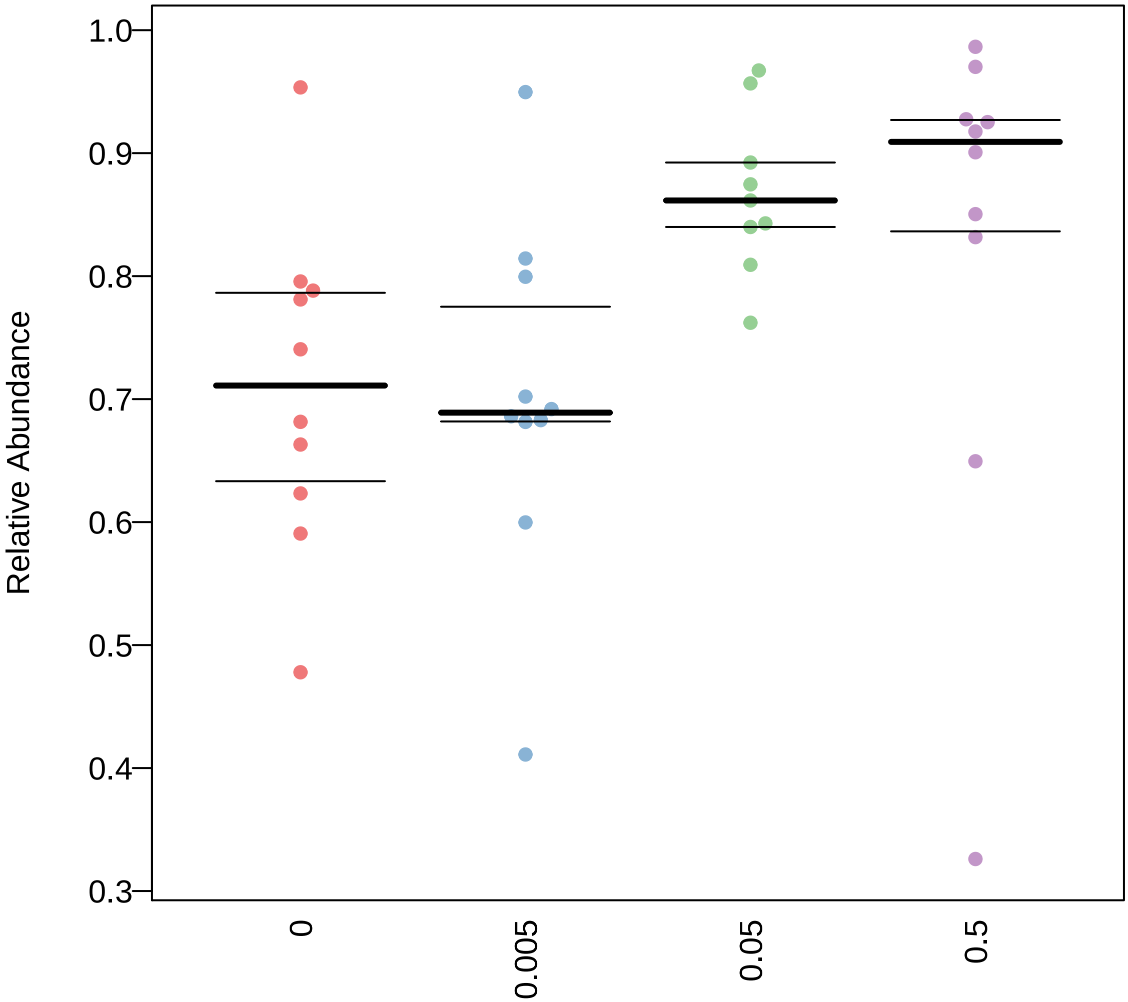

### Gram_Negative.pdf

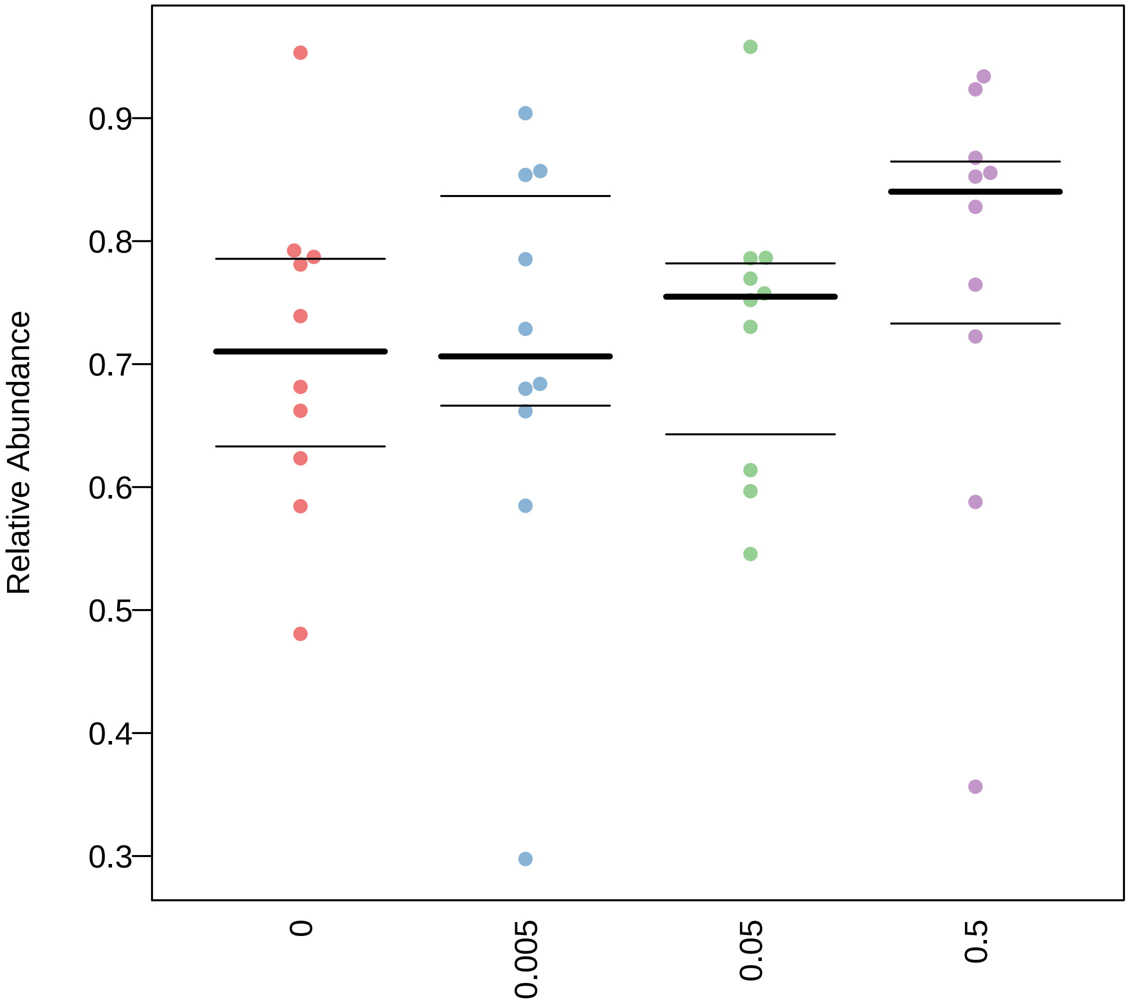

### Gram_Positive.pdf

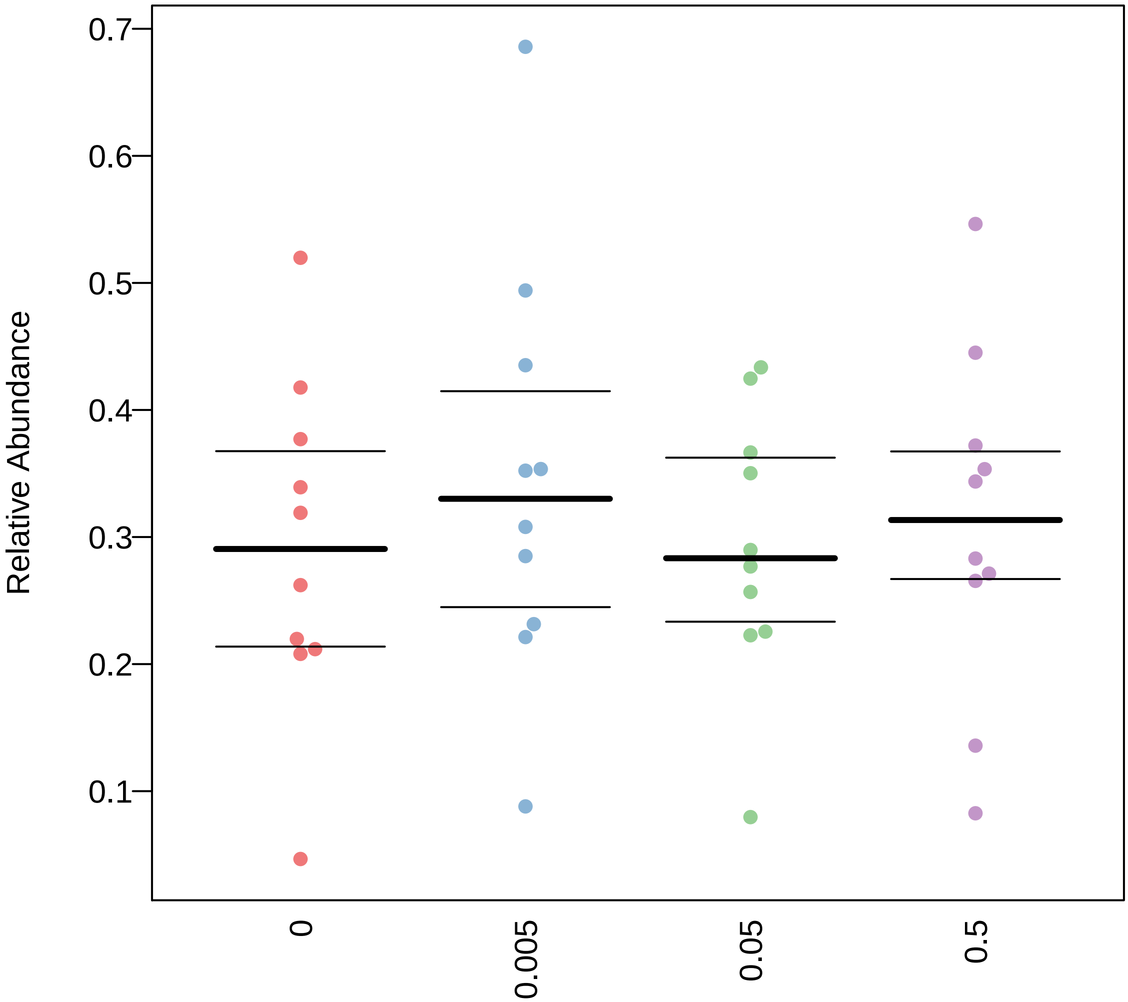

### Gram_Positive.pdf

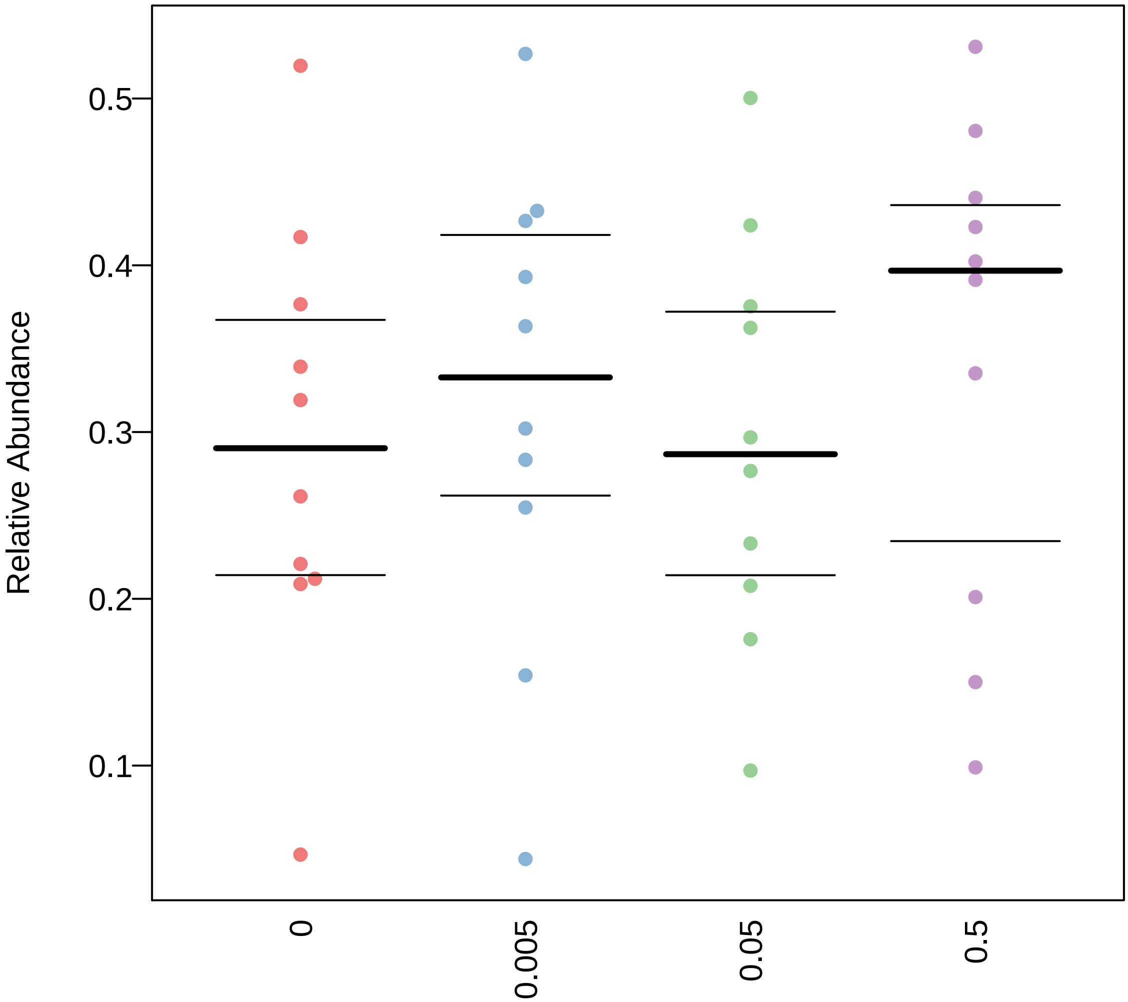

### Gram_Positive.pdf

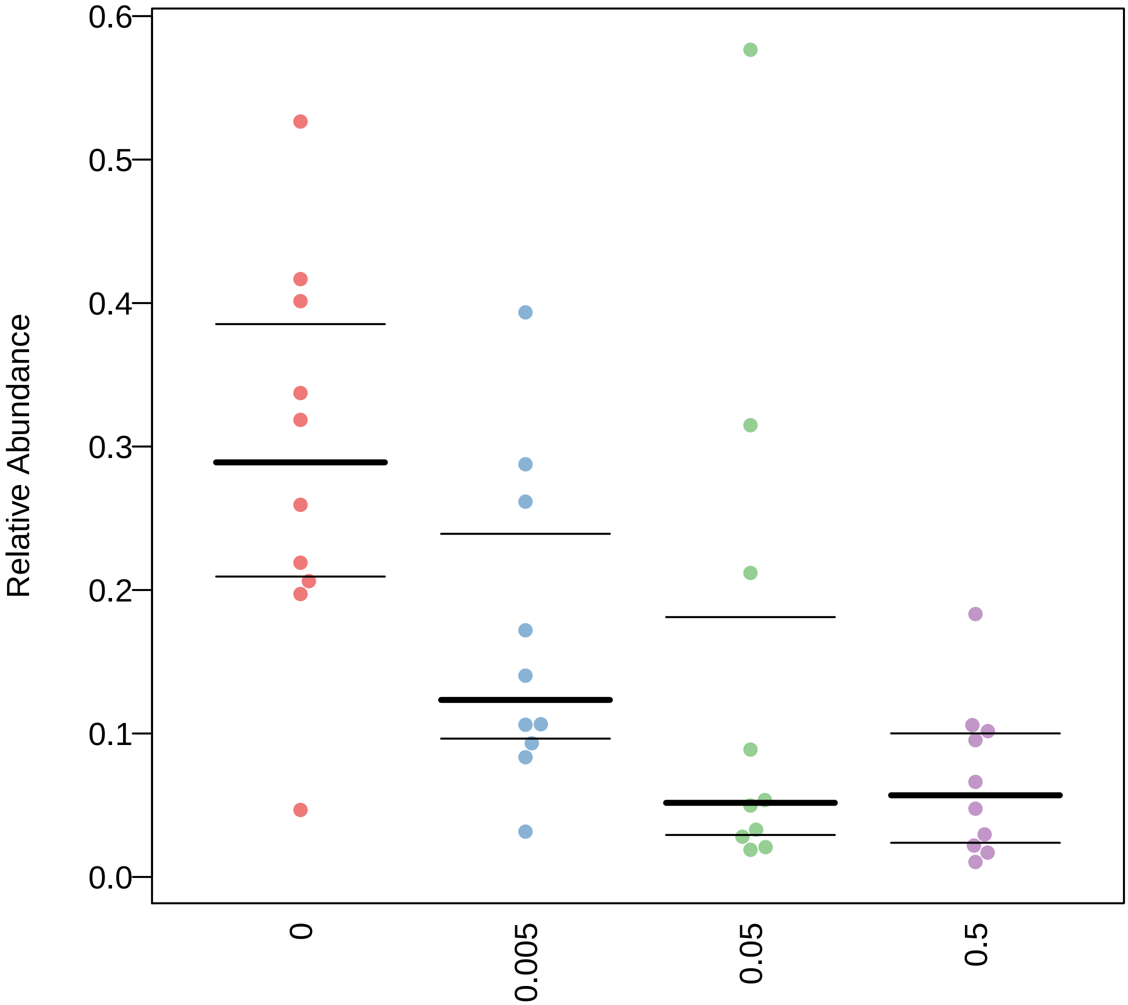

### Gram_Positive.pdf

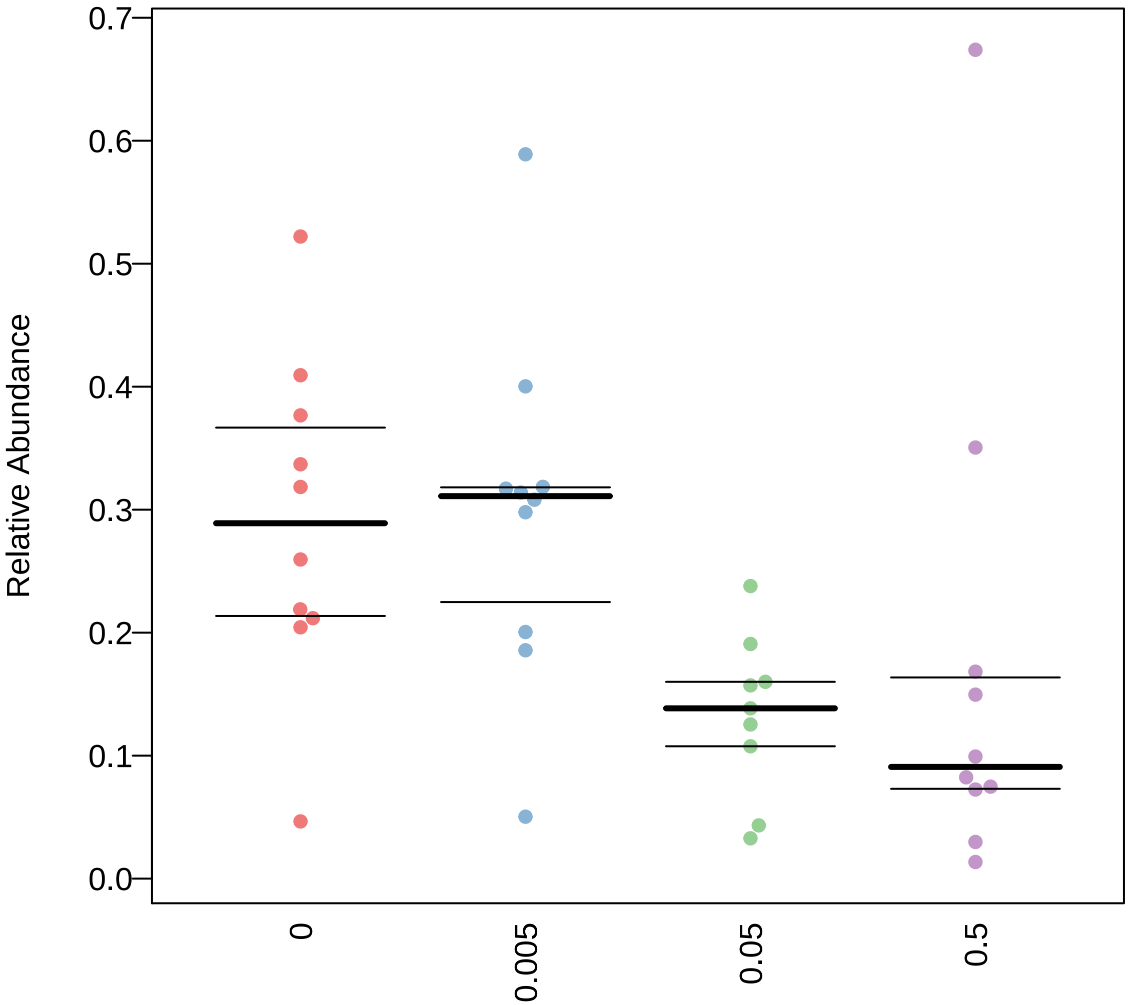

### Gram_Positive.pdf

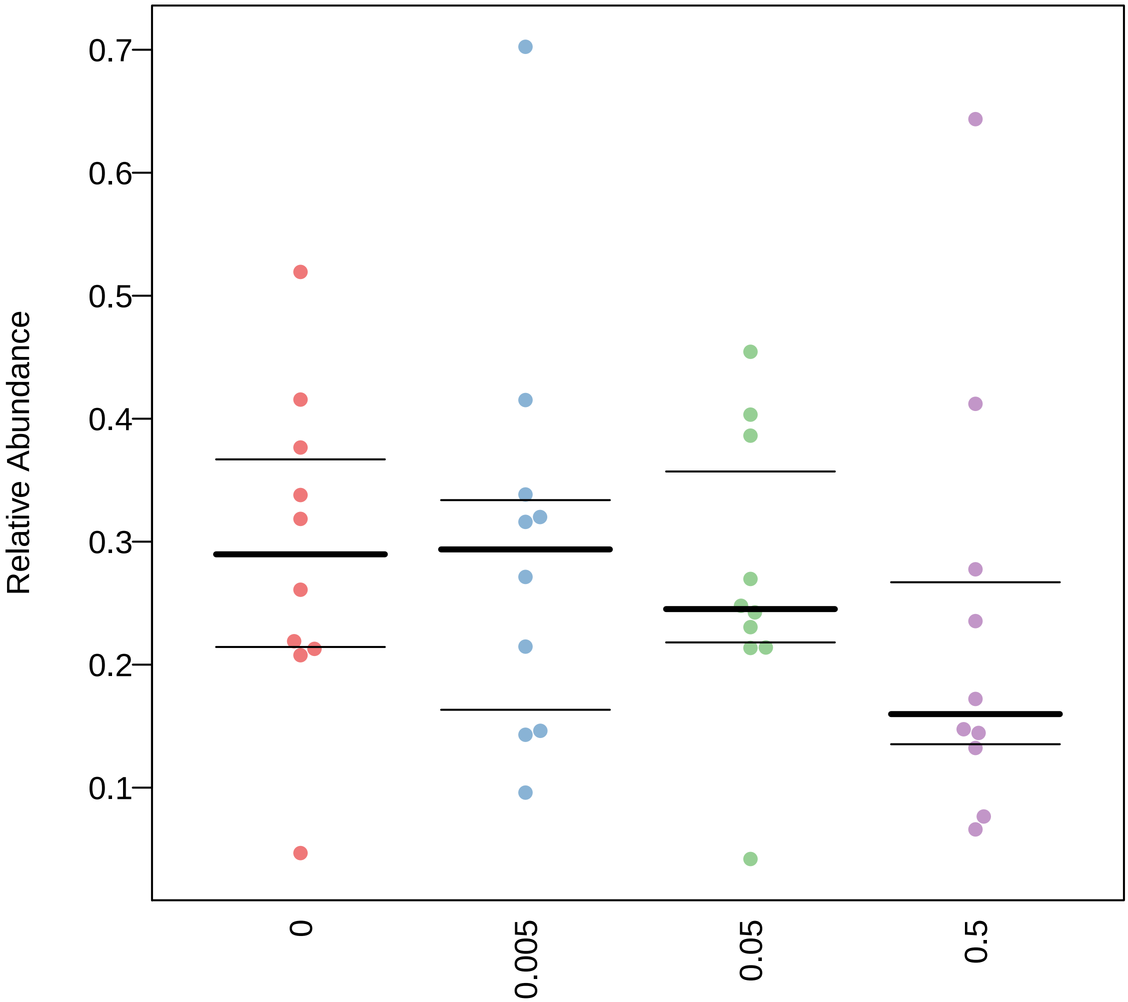

### Potentially_Pathogenic.pdf

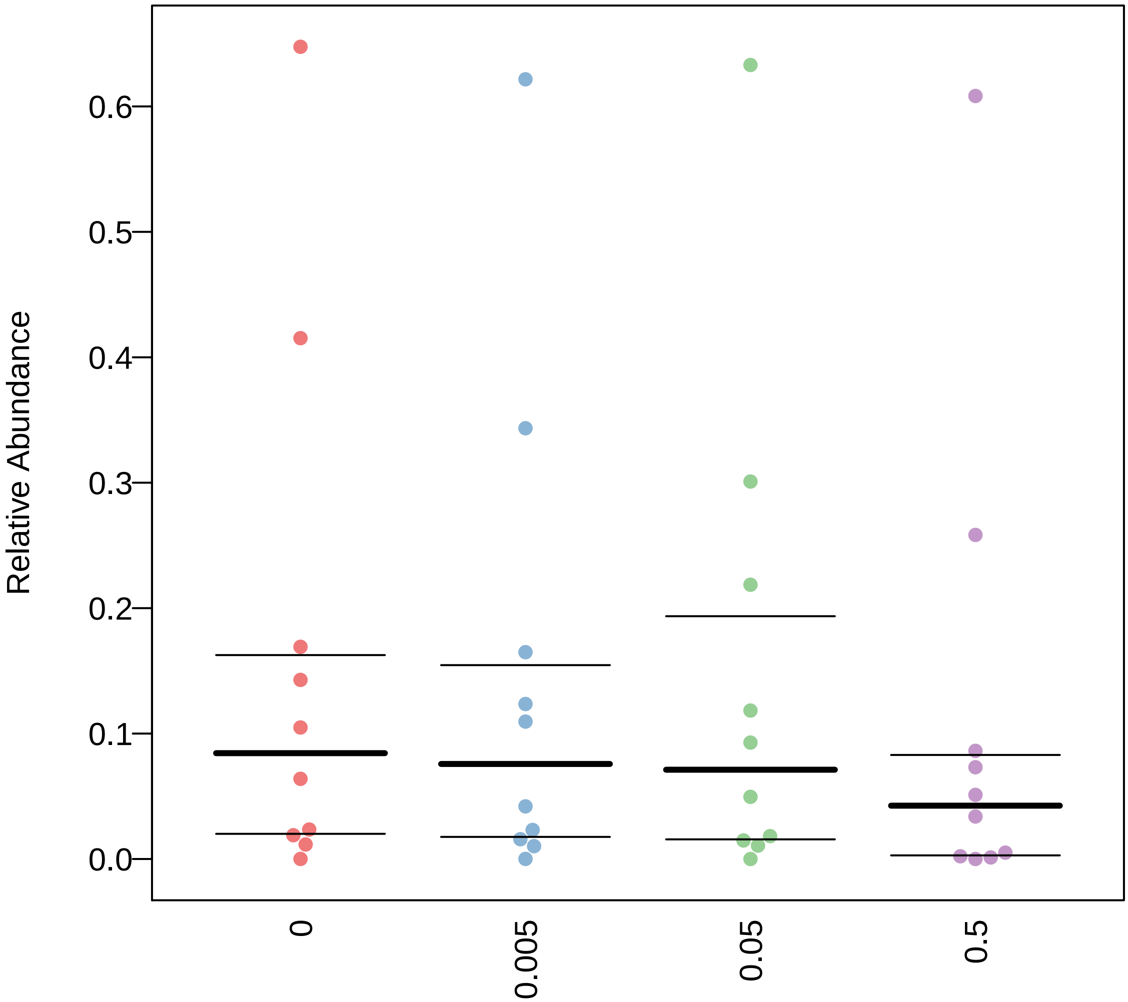

### Potentially_Pathogenic.pdf

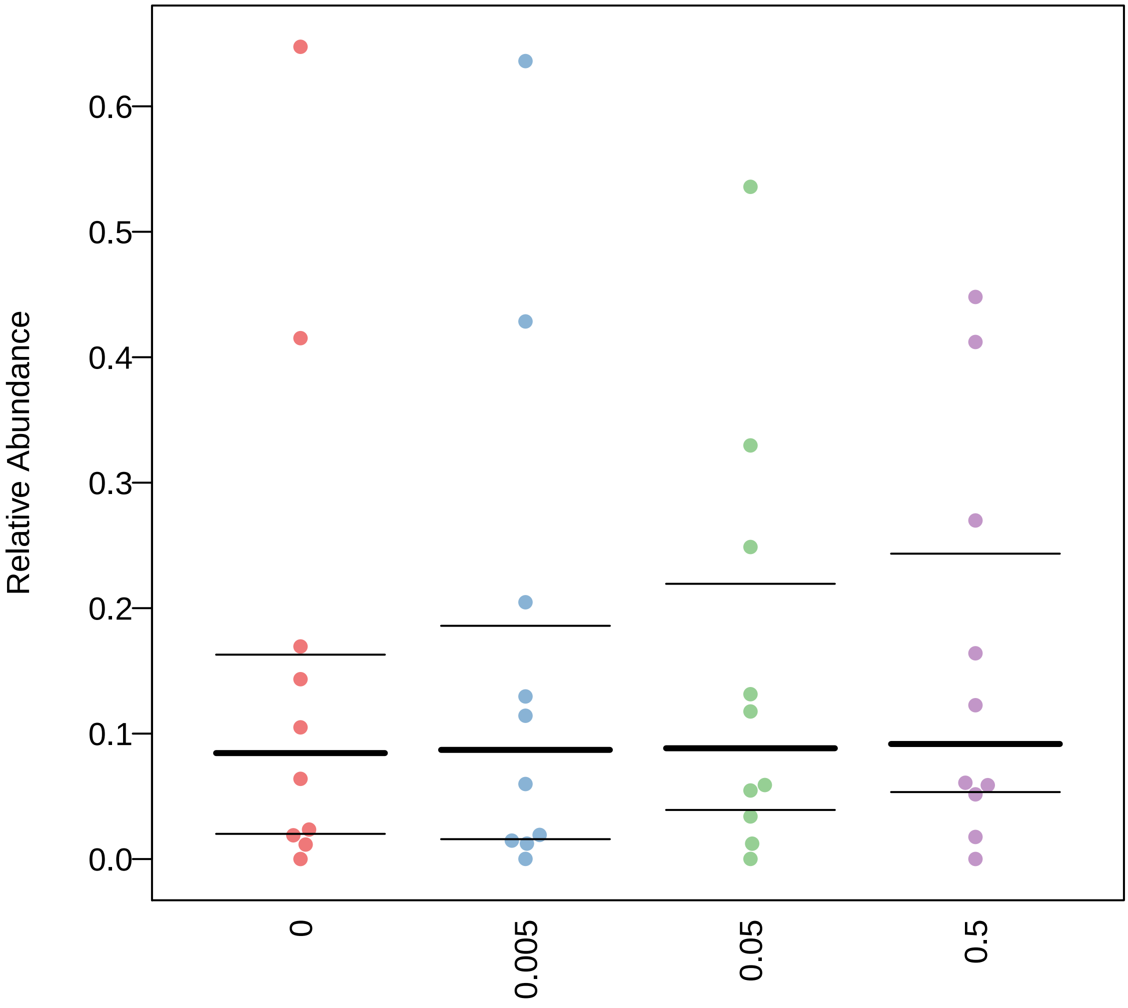

### Potentially_Pathogenic.pdf

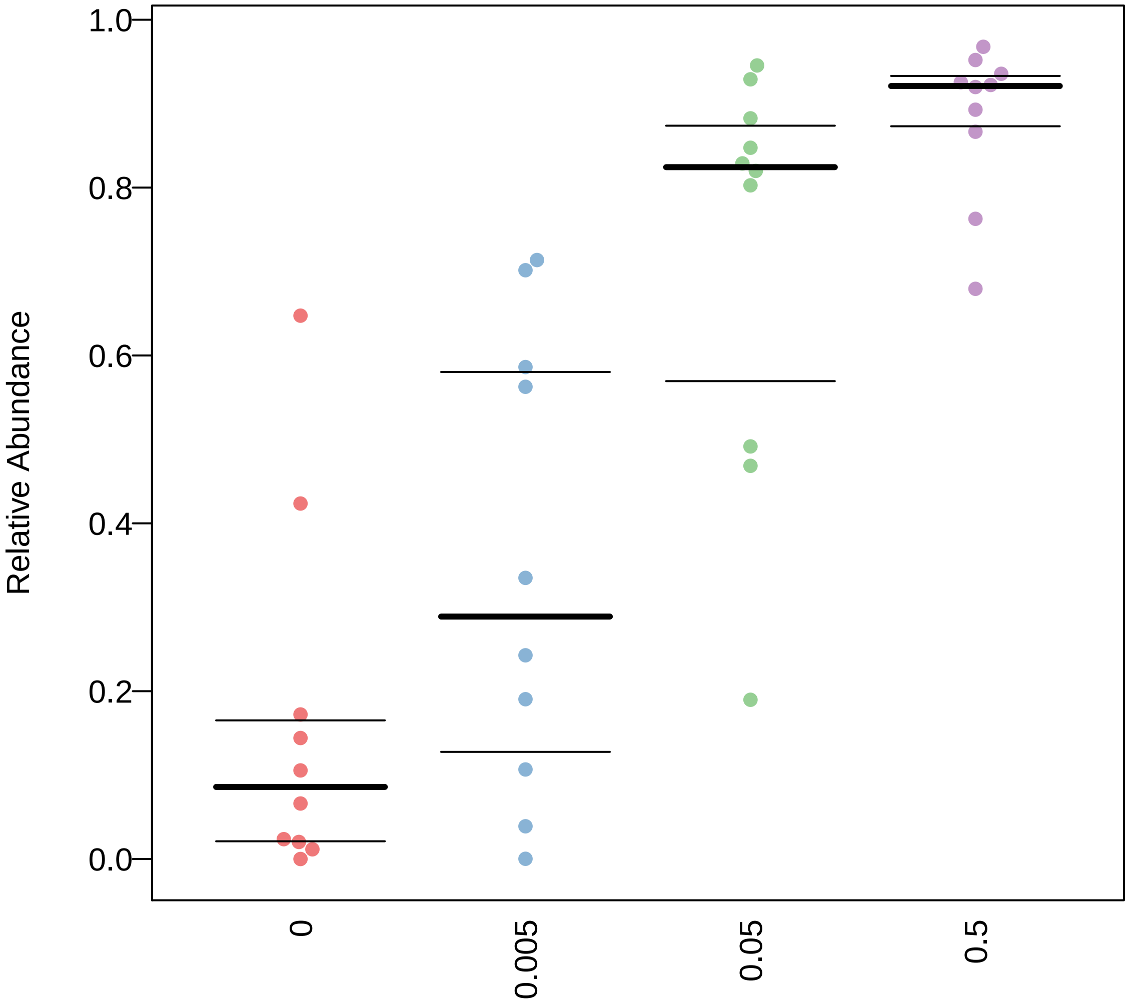

### Potentially_Pathogenic.pdf

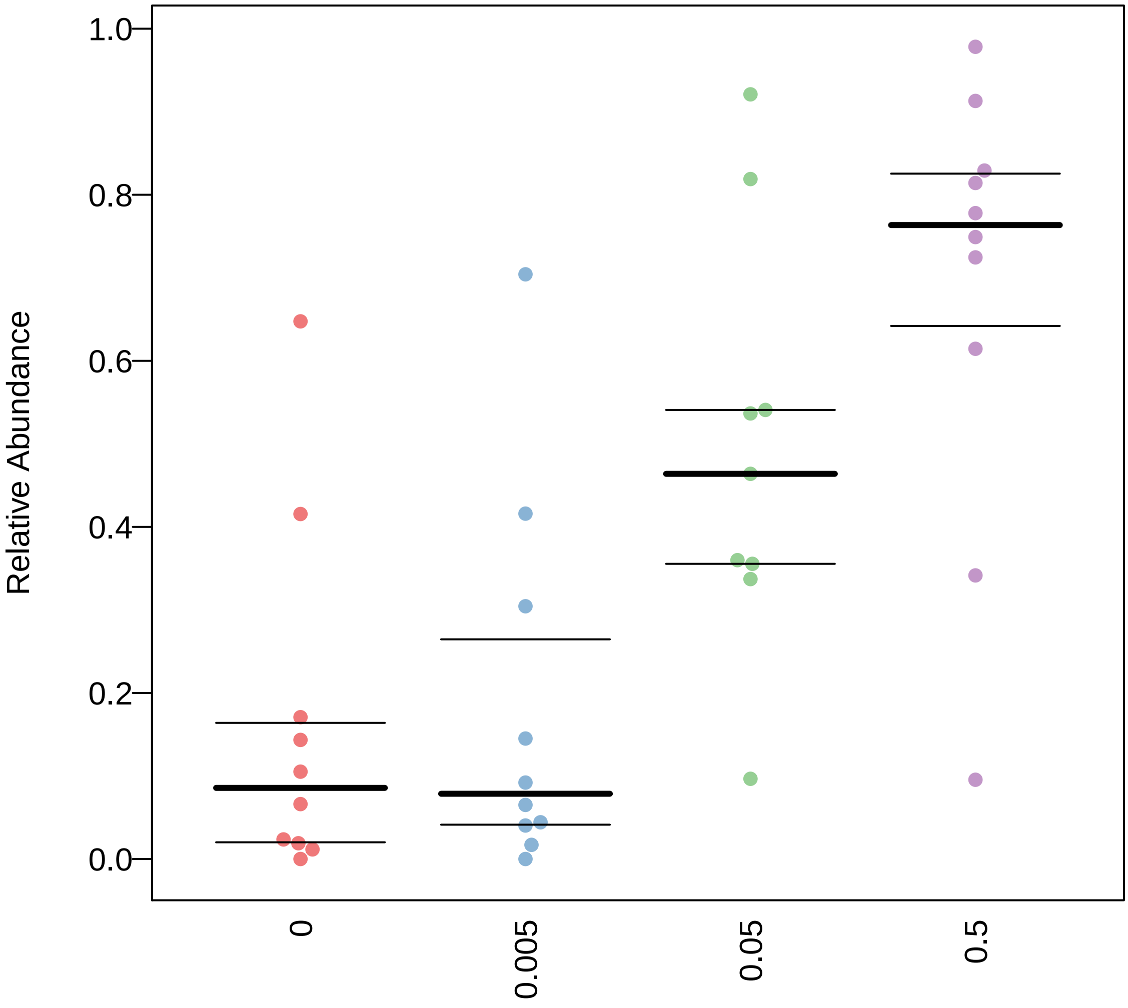

### Potentially_Pathogenic.pdf

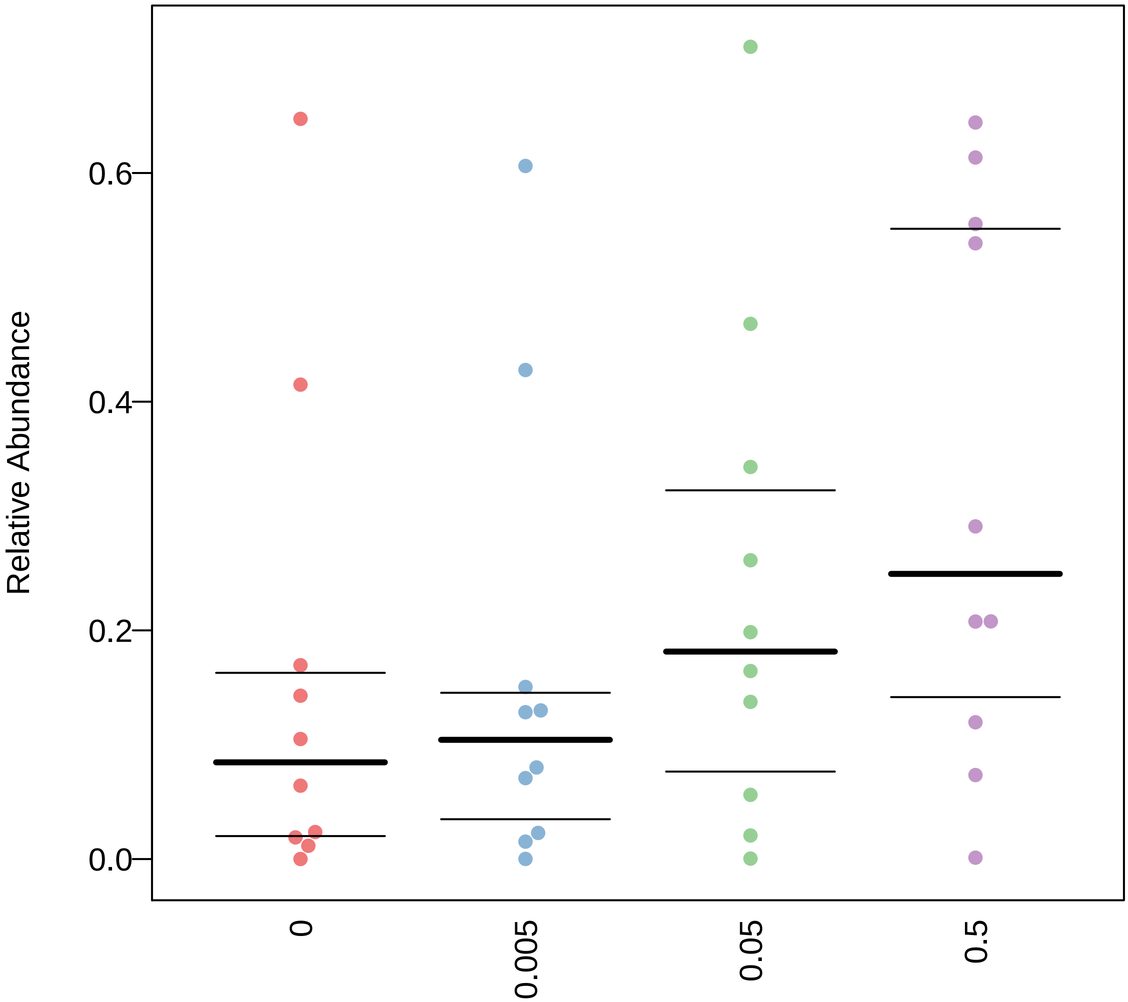

### Stress_Tolerant.pdf

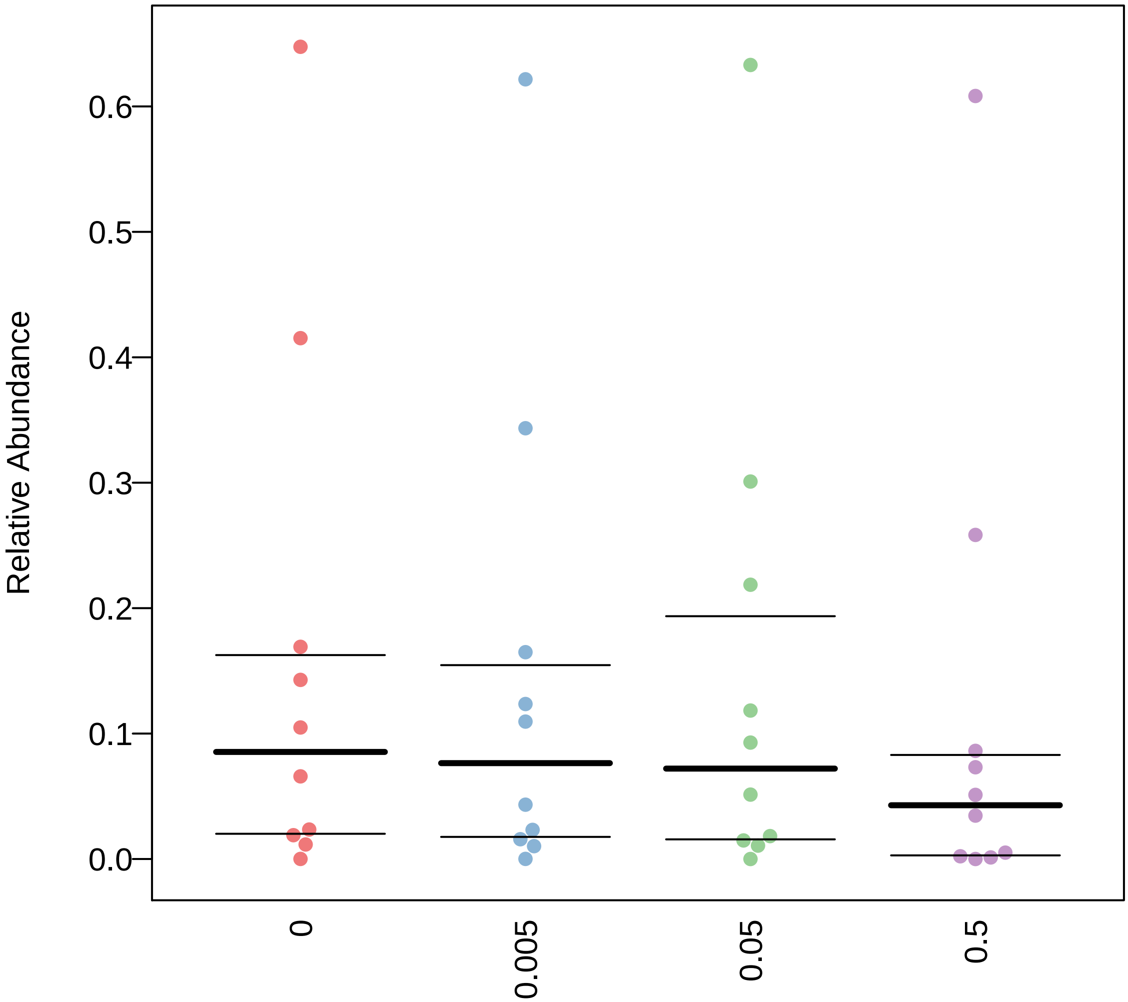

### Stress_Tolerant.pdf

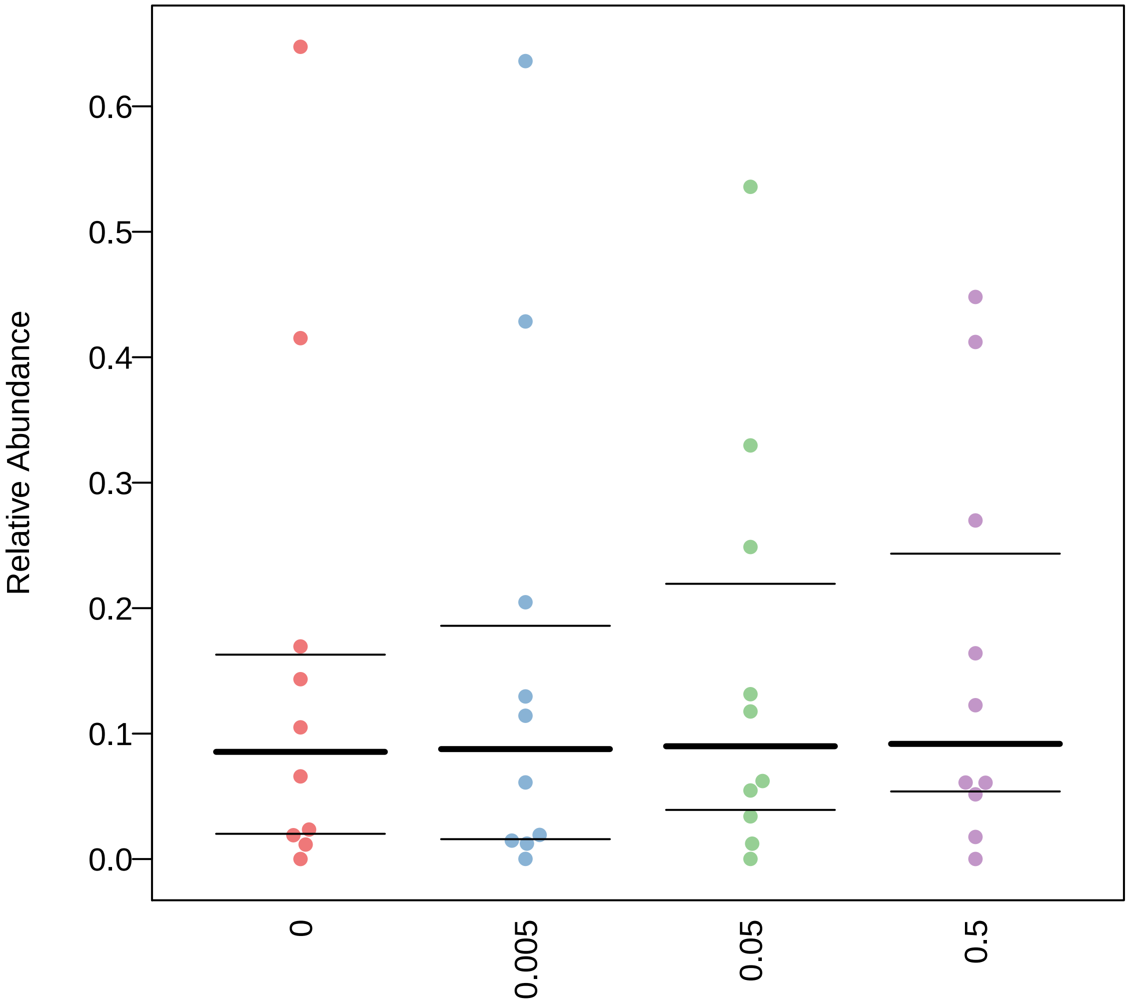

### Stress_Tolerant.pdf

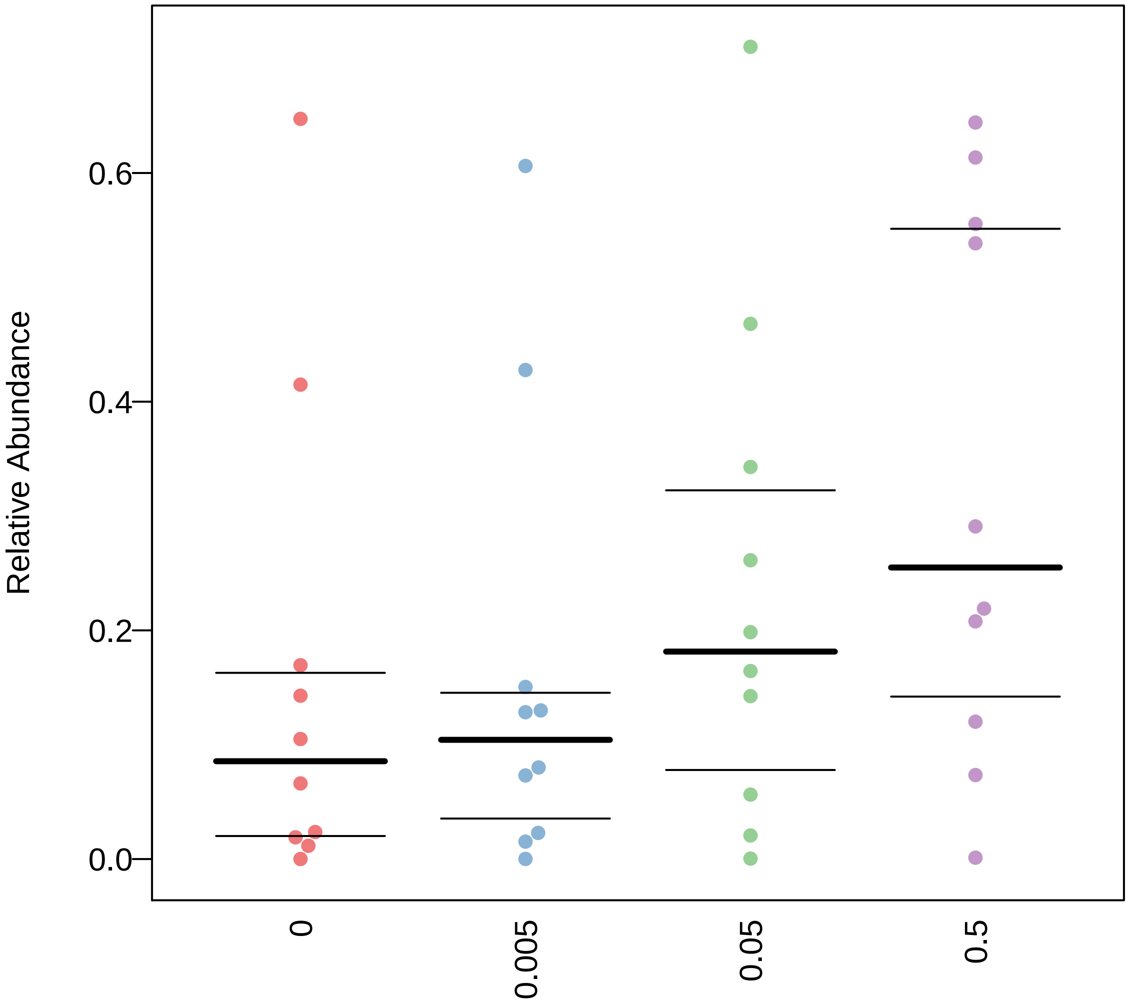
