## Supplementary Data 2 - RL&SL Digestion Chromatography for "Dietary emulsifiers alter composition and activity of the human gut microbiota *in vitro*, irrespective of chemical or natural emulsifier origin": 20_2017102367303.pdf

#### Auto-Scaled Chromatogram

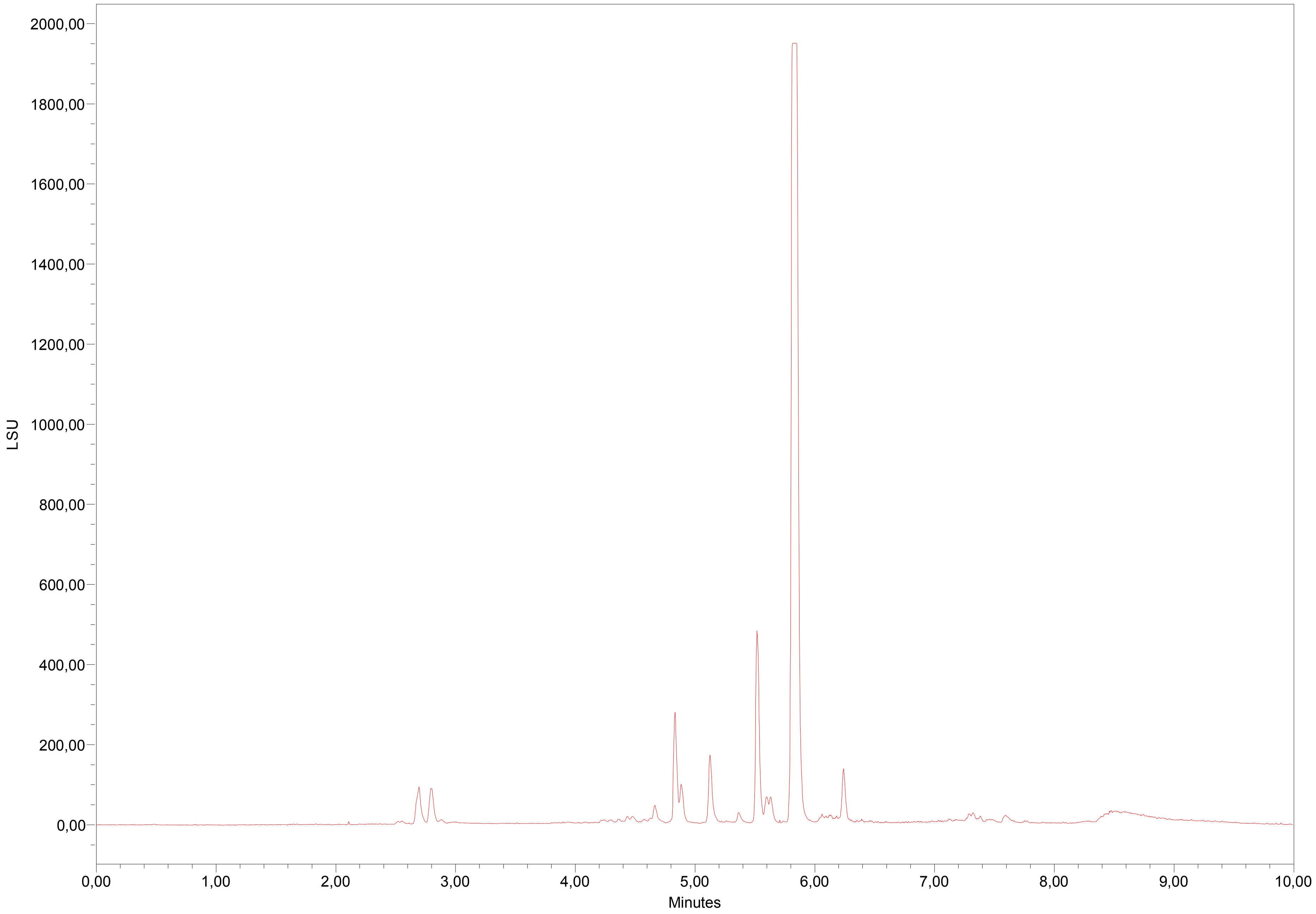


SampleName 20_20171030_S1; Vial 1:A,1; Injection 1; Channel ELSD Signal; Date Acquired 30/10/2017 14:22:49

### Chromatogram Overlay with Z Axis Offset

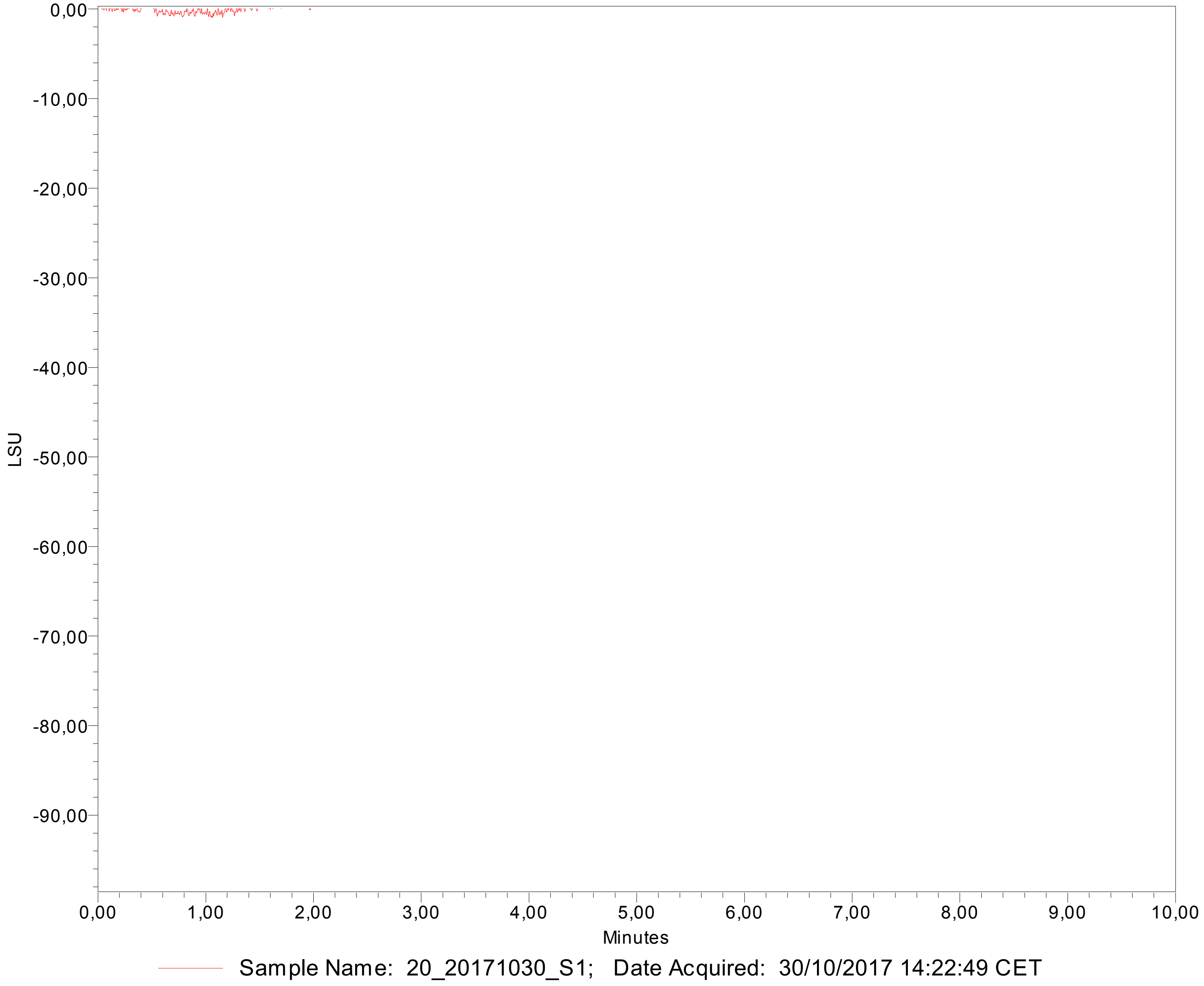

Error Log

All Peaks Table group contains information that doesn't match the data being reported.
Calibration Plot group contains information that doesn't match the data being reported.
Area Component Summary group contains information that doesn't match the data being reported.
