## Supplementary Data 2 - RL&SL Digestion Chromatography for "Dietary emulsifiers alter composition and activity of the human gut microbiota *in vitro*, irrespective of chemical or natural emulsifier origin": 20_2017102367314.pdf

### Auto-Scaled Chromatogram

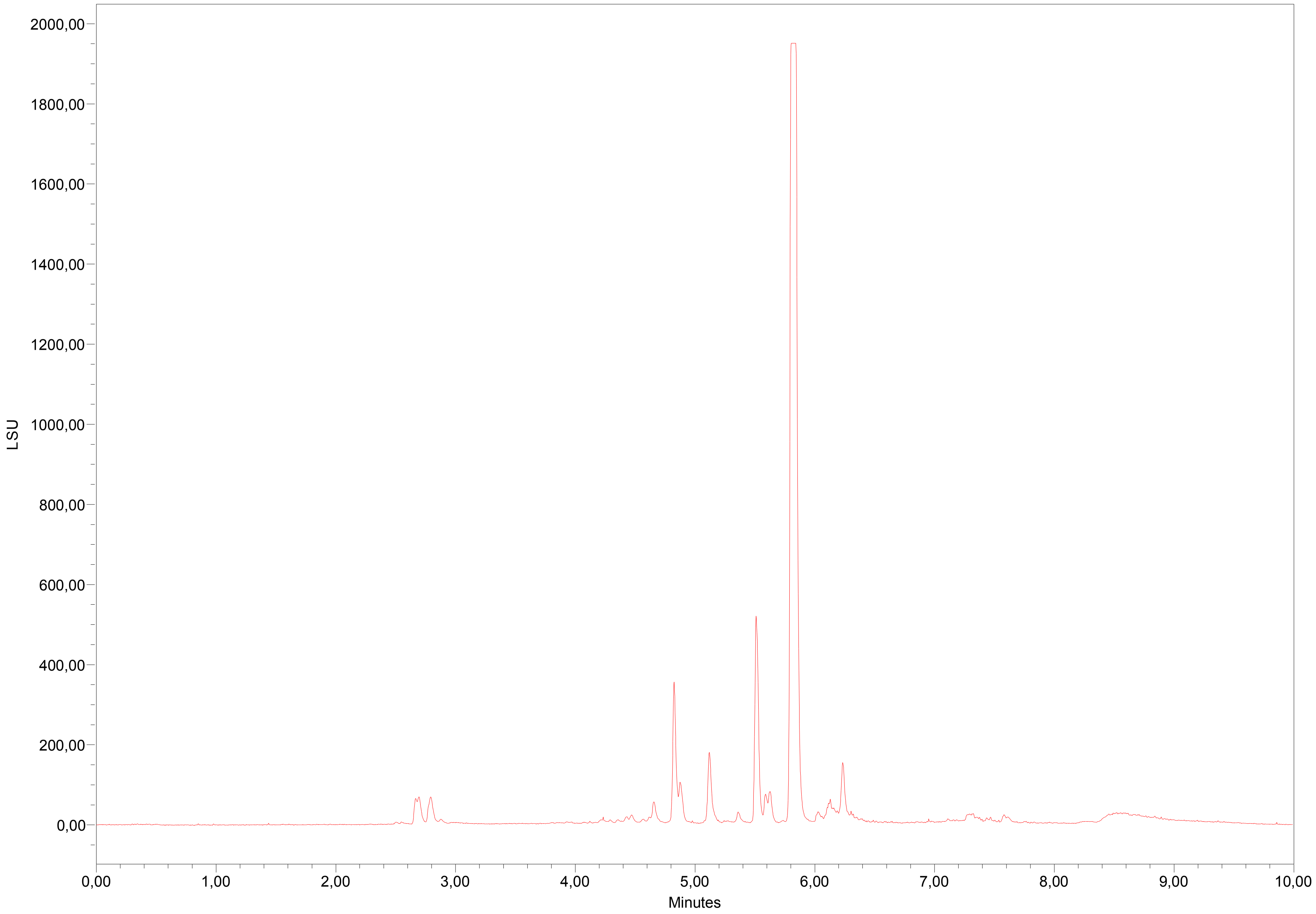


SampleName 20_20171030_S3; Vial 1:A,3; Injection 1; Channel ELSD Signal; Date Acquired 30/10/2017 14:43:52

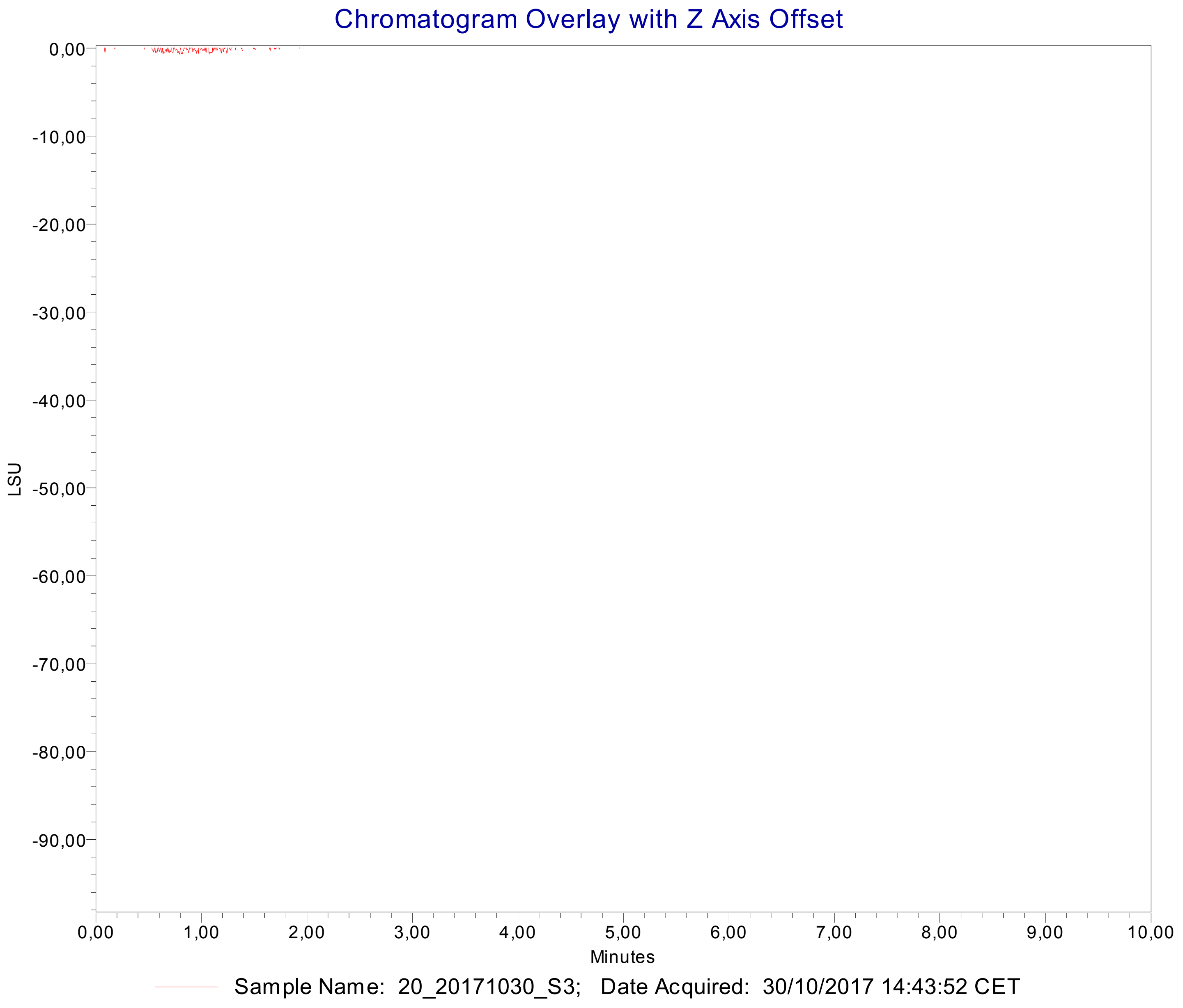

Chromatogram Overlay with Z Axis Offset
LSU
0,00
-10,00
-20,00
-30,00
-40,00
-50,00
-60,00
-70,00
-80,00
-90,00
0,00
1,00
2,00
3,00
4,00
5,00
6,00
7,00
8,00
9,00
10,00
Minutes
Sample Name: 20_20171030_S3; Date Acquired: 30/10/2017 14:43:52 CET

### Error Log

All Peaks Table group contains information that doesn't match the data being reported.
Calibration Plot group contains information that doesn't match the data being reported.
Area Component Summary group contains information that doesn't match the data being reported.
